# Persister cancer cells are characterized by H4K20me3 heterochromatin that defines a low inflammatory profile

**DOI:** 10.1101/2024.01.26.577389

**Authors:** Valentina Ramponi, Laia Richart, Marta Kovatcheva, Camille Stephan-Otto Attolini, Jordi Capellades, Alice E. Lord, Oscar Yanes, Gabriella Ficz, Manuel Serrano

## Abstract

Anti-cancer therapies may induce proliferative arrest in cancer cells in the form of senescence or drug-tolerant persistency, the latter being a reversible arrest with similarities to embryonic diapause. Here, we use mTOR/PI3K inhibition to develop and characterize a model of persistency/diapause-like arrest in human cancer cells of various origins. We show that persister and senescent cancer cells share an expanded lysosomal compartment and hypersensitivity to BCL-XL inhibition. However, persister cells do not exhibit other features of senescence, such as the loss of Lamin B1, senescence-associated b-galactosidase activity, and an inflammatory phenotype. Compared to senescent cells, persister cells have a profoundly diminished senescence-associated secretory phenotype (SASP), low activation of interferon signaling pathways and lack upregulation of MHC-I presentation. Based on a genome-wide CRISPR/Cas9 screen performed in diapause mouse embryonic stem cells (mESC), we discover that persister human cancer cells are hypersensitive to the inhibition of one-carbon metabolism. This finding led us to uncover that the repressive heterochromatic mark H4K20me3 is enriched at promoters of SASP and interferon response genes in persister cells, but not in senescent cells. Collectively, we define novel features and vulnerabilities of persister cancer cells and we provide insight into the epigenetic mechanisms underlying their low inflammatory and immunogenic activity.

## INTRODUCTION

Anti-cancer therapies are based on the infliction of damage or stress to cancer cells with the aim of triggering cell death. In practice, a fraction of cancer cells often evade cell death and enter a state of cell cycle arrest that may contribute to tumor relapse after therapy termination^1,2^. Anti-cancer therapies can lead to the acquisition of two main states of proliferative arrest: therapy-induced senescence (TIS) and drug-tolerant persistency (DTP). A critical distinction between these two states is the reversibility of the process. Persister cells efficiently reacquire their capacity to resume proliferation upon drug withdrawal^2,3^, while, in general, most senescent cells are unable to resume proliferation after therapy^4^. It is worth mentioning the concept of cancer cell dormancy, which is related to drug-persistency^5^. Cancer cells dissociated from their parental tumor can acquire a state of proliferative arrest known as dormancy upon settling in a novel microenvironment, and proliferation can be reacquired when the relation with the microenvironment is perturbed^6,7^.

Extensive research on senescence has identified a number of cellular and molecular features frequently present in this form of cell cycle arrest^8,9^. These include morphological alterations of the cytoplasm that becomes flattened and larger^10^, expansion of the lysosomal compartment^11^ associated to increased activity of the enzyme β-galactosidase^12^, high levels of reactive oxygen species (ROS)^13^, and significant alterations in chromatin structure, characterized by the loss of Lamin B1 (LMNB1) from the nuclear envelope and the formation of senescence-associated heterochromatin foci (SAHF)^14–16^. Senescent cells are also characterized by the senescence-associated secretory phenotype (SASP), which is composed of a complex and dynamic mixture of cytokines, chemokines, and matrix remodeling proteins^17,18^. The SASP produced intratumorally by senescent cells has been demonstrated to be partly responsible for the exacerbation of immunosuppression post-chemotherapy, mostly by producing myeloid-attracting chemokines^4,18,19^ Consequently, there has been a strong focus on developing therapies known as senolytics, which selectively target and eliminate senescent cells^20,21^. Indeed, multiple preclinical studies have shown that senolytic drugs when combined with standard cancer therapies improve therapeutic outcome (reviewed in^22^).

In comparison, less is known about the molecular and cellular features of persister cells. Recent studies have identified specific characteristics of persister cells, including slow cell cycle^23–28^, altered metabolism with an increase in autophagy^29,26^ and dependency on oxidative phosphorylation (OXPHOS)^30^, altered epigenetic traits with increased repressive chromatin histone marks H3K9me3, H3K27me3 and decreased H3K4me3^23,25^, a stem-cell like phenotype ^24^, and immune evasion^31,32^. Nevertheless, there is still no consensus about hallmarks of persister cells. Utilizing *in vitro* cell lines represents a valuable tool for uncovering features of persister cells, but the current *in vitro* models have been built around therapies that can also induce senescence^23,26,33–42^. In this regard, an experimental model that induces only persistency, and not senescence, may facilitate the discovery of discriminating features.

A remarkable recent finding has been the observation cancer persister cells and embryonic diapause cells share an extensive transcriptional program^26,27^. Embryonic diapause is a reversible state of suspended development triggered by unfavorable nutrient conditions that can be mimicked in human and mouse embryonic stem cells (ESC) *in vitro* using a dual mTOR/PI3K inhibitor, INK128^43,44^. Interestingly, the transcriptomes of persister colorectal and breast cancer cells revealed a high similarity with the transcriptional signatures of *in vitro* and *in vivo* embryonic diapause^26,27^. Diapause ESC and persister cells also share an increased autophagic activity^26^. Based on these evidences, cancer persister cells are also referred to as diapause-like cancer cells (DLCC)^26,27^.

Here, we exploit the similarities between cancer persistency and embryonic diapause to establish an *in vitro* model of cancer diapause-like arrest based on the dual inhibition of mTOR/PI3K. Using this system, we uncover shared and unique traits of senescent and persister cancer cells. We leverage the resemblance of cancer persistency to embryonic diapause and use a genome-wide CRISPR/Cas9 screening approach in mouse ESC. In this manner, we identify and validate novel vulnerabilities specific to persister/diapause-like cells. Finally, we identify a discriminatory epigenetic state between senescent and persister cells that explains their differential expression of inflammatory programs. Collectively, our work highlights the unique molecular features of persister/diapause-like cancer cells, and uncovers potential therapeutic approaches for improved tumor control.

## RESULTS

### Dual inhibition of mTOR/PI3K induces a persister/diapause-like state in cancer cells

Previous investigators have reported that standard chemotherapeutic treatments, such as irinotecan and docetaxel can induce a diapause-like state in cancer cells^26,27^. However, these chemotherapeutic treatments also trigger alternative fates, such as apoptosis and senescence. With the goal of achieving a more uniform diapause-like response, we treated a variety of human cancer cell lines with INK128, a dual inhibitor of mTOR and PI3K, previously reported to trigger diapause in mouse and human embryonic stem cells^43,44^. All tested cancer cell lines (melanoma SK-Mel-103, lung cancer A549, H1299 and H226, and breast cancer MCF7 and MDA-MB-231) underwent a profound and stable suppression of proliferation upon treatment with INK128 for 7 days (**Figures 1A-B** and **S1A-B**). Analysis of DNA content indicated that INK128-induced arrest occurred primarily at the G1 phase of the cell cycle (**Figure S1C**). Annexin V/PI staining did not detect apoptotic (Annexin V+ PI+) cells at 48h of INK128 treatment; however, after 7 days (with fresh media every 2-3 days and always in the presence of the drug), INK128-treated cells presented a mild, albeit significant, increase in the number of apoptotic cells compared to proliferating control cells (from ≤2% in proliferating cells to ≤4% in INK128-treated cells) (**Figure S1D**). Importantly, upon removal of INK128, cultures resumed proliferation (**Figures 1A-B** and **S1B**), thereby recapitulating a key hallmark of the diapause-like state, namely, its efficient reversibility. INK128-treated cells were also more resistant to doxorubicin than proliferating cells (**Figure S1F**), thereby recapitulating a defining feature of drug-tolerant persistency^2^. Cancer dormancy may share similarities with drug-tolerant persistency (or diapause-like arrest) (see Introduction). Interestingly, we observed that *DEC2* (also known as *BHLHE41*) and *p27* (*CDKN1B*), two markers of *in vivo* cancer dormancy^45,46^, were upregulated in INK128-treated cells (**Figures 1C** and **S1E**). To further explore the similarities between INK128-induced diapause-like cancer cells and embryonic diapause, we compared their transcriptomes. We performed RNA-sequencing of A549 and SK-Mel-103 cells with or without INK128 treatment. Gene Set Enrichment Analysis (GSEA) revealed that INK128 induced the downregulation of the mTOR pathway and the unfolded protein response (UPR), which is consistent with the inhibition of mTOR and a reduction in protein synthesis (**Figure 1D**). We also observed downregulation of cell cycle pathways and MYC target pathways, which are hallmarks of embryonic diapause^47^. When focusing on the previously reported signatures of mouse embryonic diapause (a signature of downregulated genes and a signature of upregulated genes)^26^, both were significantly modified as in embryonic diapause in the transcriptomes of INK128-treated cells (**Figures 1E** and **S1G**). Moreover, based on the A549 and SK-Mel-103 transcriptomes, we generated INK128 signatures of downregulated and upregulated genes (**Table S1** and **S2**) and tested them against previously published RNA sequencing datasets from persister colorectal cancer cells^26^. Interestingly, the INK128 signatures were significantly changed in DTP colorectal cancer cells in the same way as in INK128-treated A549 and SK-Mel-103 cells (**Figure 1F**). Of note, INK128-treated cancer cells presented a significant enrichment of the TGF-β (TFGβ) and WNT/β-Catenin (WNT/βCAT) pathways, both of which have been previously linked to tumor dormancy *in vivo*^46,48,49^, again suggesting the similarity with dormancy. Collectively, these data indicate that mTOR/PI3K inhibition in cancer cells triggers a diapause-like arrest that resembles embryonic diapause and related states in cancer cells known as DTP and tumor dormancy. Based on this, we will refer to INK128-treated cancer cells as persister/diapause-like cancer cells.

**Figure 1.**
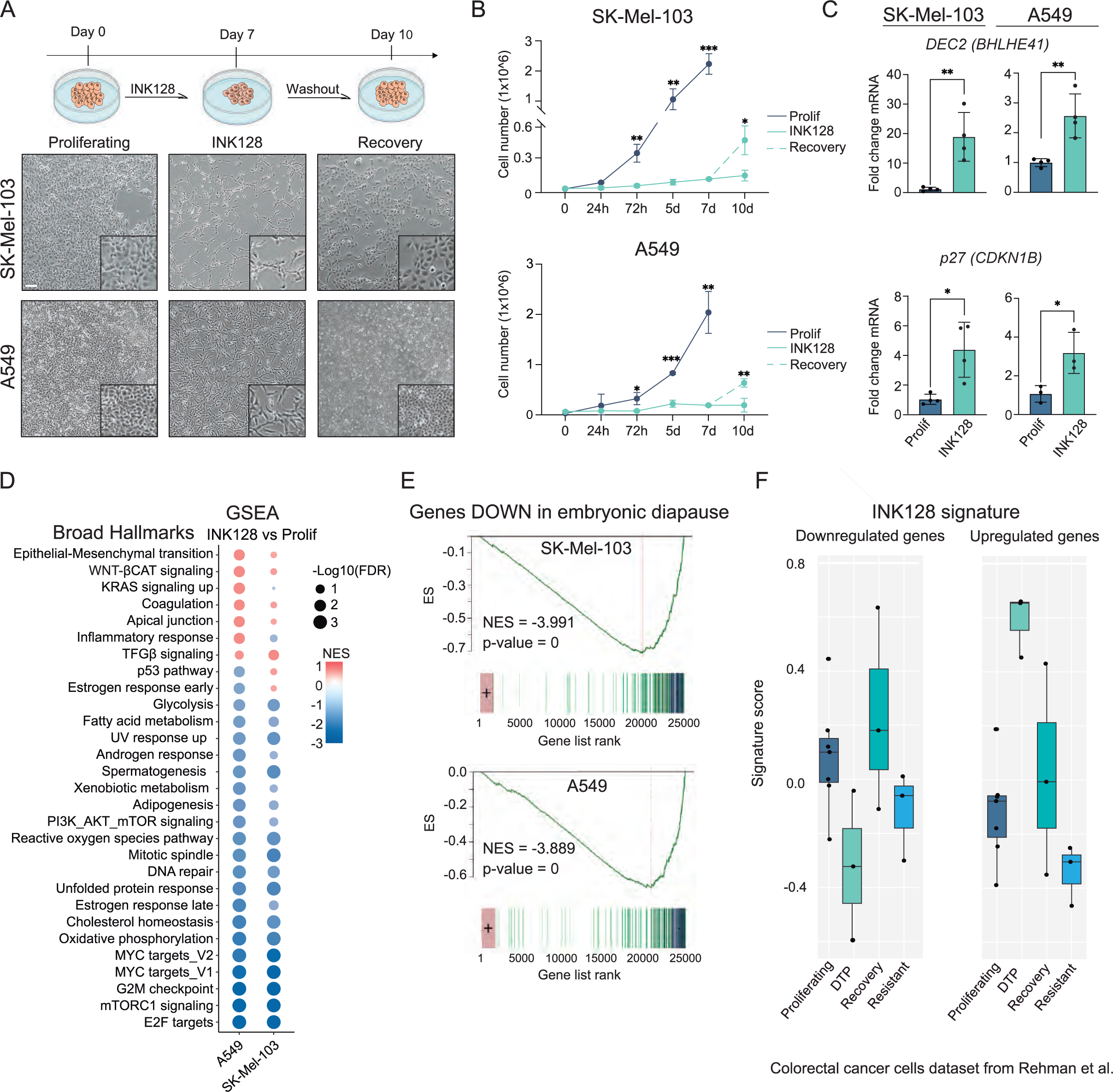
INK128-treated cells undergo a reversible cell cycle arrest with features of drug-tolerant persistency and embryonic diapause. (A) Brightfield pictures of SK-Mel-103 and A549 cells untreated (proliferating), treated with INK128 (100μM and 200μM respectively) for 7 days, and after INK128 withdrawal for 3 days. Scale bar is 100μm. (B) Proliferation curve of SK-Mel-103 and A549 cells untreated (proliferating), treated with INK128 and after INK128 withdrawal (n=3). Cells were counted using a hemocytometer after 24 hours from plating, 3 days, 5 days, 7 days and 10 days (or 3 days after INK128 withdrawal) (n=3). ***, P < 0.001; **, P < 0.01; *, P < 0.05; multiple unpaired t test. (C) mRNA expression levels of *DEC2* (*BHLHE41*) and *p27* (*CDKN1B*) in untreated (proliferating) and INK128 treated SK-Mel-103 and A549 cells, measured by qRT-PCR (relative to the average expression of housekeeping genes *ACTB* and *GAPDH*). **, P < 0.01; *, P < 0.05; unpaired Student’s t test, compared to proliferating controls (n= 3 or 4). (D) Gene set enrichment analysis (GSEA) of INK128-treated Sk-Mel-103 and A549 cells in comparison to untreated (proliferating) controls. Normalised Enrichment Score (NES) is colour-coded, while statistical significance (FDR: False Discovery Rate) is encoded in the size of the dots. (E) GSEA of the signatures “Embryonic diapause_down” in INK128-treated versus proliferating SK-Mel-103 and A549 cells. Normalised Enrichment Score (NES) is colour-coded, while statistical significance (FDR: False Discovery Rate) is encoded in the size of the dots. (F) Signature scores of control proliferating cells, drug-tolerant persistency in response to irinotecan (DTP), cells that had exited persistency upon withdrawal of irinotecan (recovery), and cells that were resistant to irinotecan and did not undergo DTP (resistant) (published datasets from Rehman et al^26^), when interrogated for the expression of the INK128 downregulated signature and the INK128 upregulated signature (proliferating n=8; all the other conditions n=3).

### Persister/diapause-like cancer cells lack a senescence-associated secretory phenotype

We next wanted to compare the transcriptomes of INK128-induced persister/diapause-like cancer cells with those of therapy-induced senescent cancer cells. We induced senescence using doxorubicin (a genotoxic agent) or palbociclib (a targeted therapy that inhibits the CDK4/6 kinases) in A549 and SK-Mel-103 cells, and compared their transcriptional profiles with those of the untreated parental cells. Principal component analysis (PCA) showed that persister/diapause-like, senescent and proliferative cells formed three distinct and clearly separated clusters of cells (**Figures 2A** and **S2A**). GSEA revealed a number of pathways that were differentially affected in persister/diapause-like cells versus therapy-induced senescent cells. The most notable finding was a large group of pathways related to secretion and inflammation that were upregulated in senescence, but not in persister cells (**Figure 2B**). There were no pathways selectively upregulated in persister/diapause-like cells, but not in senescent cells. Among the commonly regulated pathways, only two pathways were upregulated in both persister/diapause-like cells and therapy-induced senescent cells, related to WNT/βCAT and TGFβ signaling, and numerous pathways related to proliferation were commonly downregulated as expected (**Figure 2B**). Analysis by GSEA of a proteomic-based SASP signature of oncogene-induced senescence and radiation-induced senescence^50^ indicated a complete absence of SASP in persister/diapause-like cells compared to senescent cells (**Figure 2C**). Indeed, when analyzed at the individual level of SASP factors upregulated by therapy-induced senescence in SK-Mel-103 and A549 cells, the large majority were absent in INK128-treated cells (**Figures S2B** and **S2C**). Importantly, the SASP signatures were also significantly absent in persister cells from human primary colorectal cancers compared to resistant or proliferating cancer cells (**Figure 2D**. Again, this reinforces the concept that INK128-arrest is a good model of drug persistency in real cancers. We conclude that the absence of SASP in persister/diapause-like cancer cells is a key distinguishing feature from senescent cancer cells.

**Figure 2.**
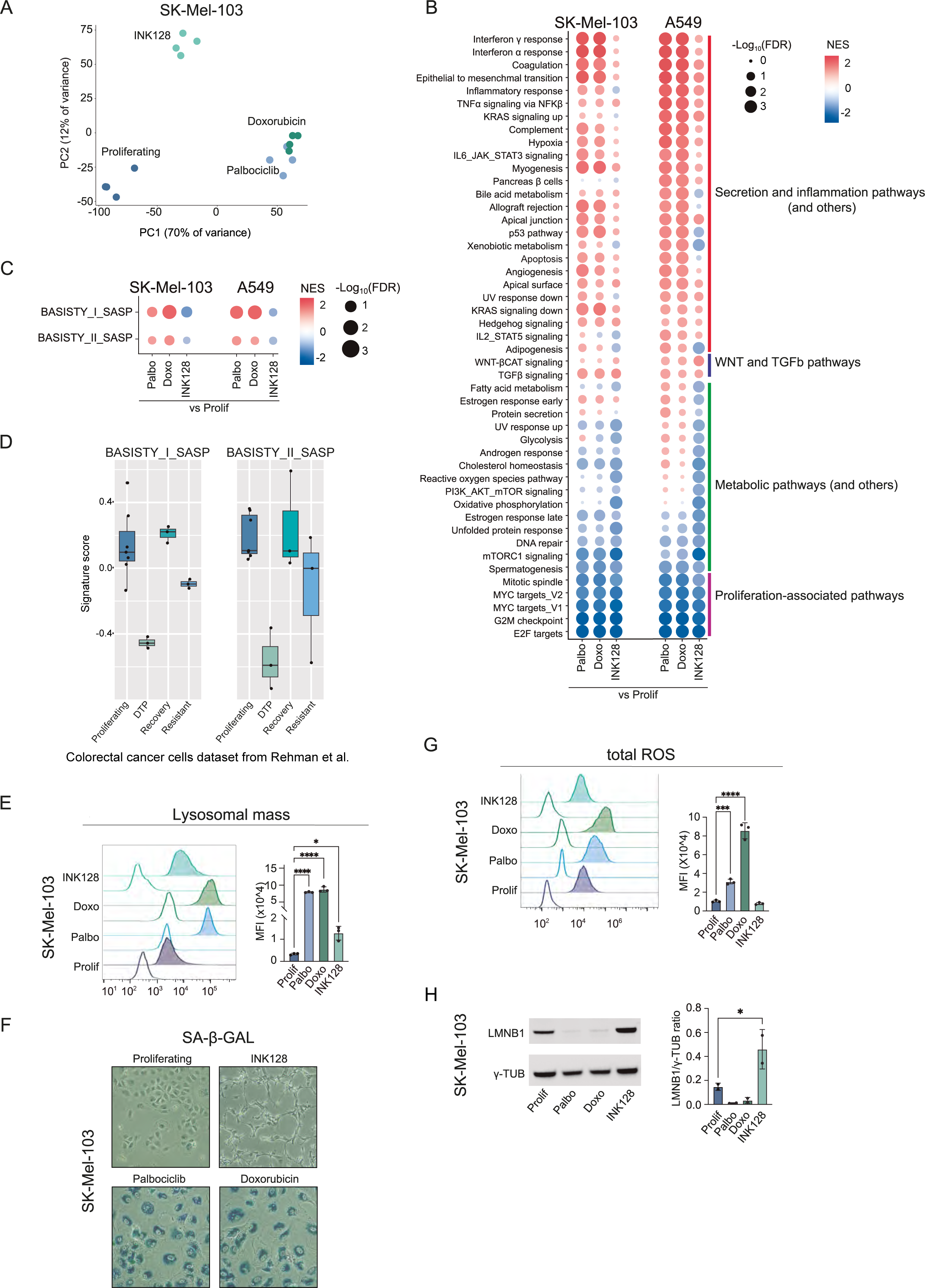
Hallmarks of senescence in persister/diapause-like cancer cells. (A) Principal Component Analysis (PCA) of proliferating, palbociclib-treated, doxorubicin-treated and INK128-treated SK-Mel-103 (n=4). (B) Gene set enrichment analysis (GSEA) of palbociclib-treated, doxorubicin-treated and INK128-treated versus proliferating SK-Mel-103 and A549 cells. Only pathways significantly differentially regulated in SK-Mel-103 or A549 cells are represented. Normalized enrichment scores (NES) and Log10FDR are indicated. (C) Signature scores of control proliferating cells, drug-tolerant persistency in response to irinotecan (DTP), cells that had exited persistency upon withdrawal of irinotecan (recovery), and cells that were resistant to irinotecan and did not undergo DTP (resistant) (published datasets from Rehman et al26), when interrogated for the expression of the SASP signatures from Basisty et al.50 (proliferating n=8; all the other conditions n=3). (D) Dot plot summarizing the results of pre-ranked GSEA analysis testing the differential enrichment of SASP gene sets^50^ in palbociclib-treated, doxorubicin-treated and INK128-treated versus proliferating Sk-Mel-103 and A549 cells. Normalised Enrichment Score (NES) is colour-coded, while statistical significance (FDR: False Discovery Rate) is encoded in the size of the dots. (E) Flow cytometry analysis of Lysotracker Red in proliferating, TIS (palbo and doxo) and DLCCs SK-Mel-103. Representative histograms showing the fluorescence signal of each stained sample and its unstained control (uncolored histogram). Quantification of the mean fluorescence intensity (MFI) after autofluorescence subtraction (n=3 SK-Mel-103, n=4 A549). ****, P < 0.0001; ***, P < 0.001; **, P < 0.01; *, P < 0.05; one-way ANOVA compared to proliferating control cells. (F) SA-β-GAL staining (blue) of proliferating, palbo-treated, doxo-treated and INK128-treated SK-Mel-103 after 7 days from the beginning of the treatment. (G) Flow cytometry analysis of total ROS in proliferating, palbo-treated, doxo-treated and INK128-treated SK-Mel-103 cells. Representative histograms showing the fluorescence signal of each stained sample and its unstained control (uncolored histogram). Quantification of the MFI after autofluorescence subtraction (n=3). ***, P < 0.001; **, P < 0.01; one-way ANOVA compared to proliferating control cells. (H) LMNB1 and γ-TUBULIN protein levels in proliferating, palbo-treated, doxo-treated, INK128-treated SK-Mel-103 cells. A representative immunoblot is shown (n=2). *, P < 0,05; one-way ANOVA compared to proliferating controls.

### Cellular features distinguishing persister/diapause-like and senescent cancer cells

To further extend the comparison between persister/diapause-like and senescent cancer cells, we analyzed cellular features that have been previously described as hallmarks of senescence. Senescent cells are characterized by an increased lysosomal content and this is also reflected in high levels of the lysosomal enzyme β-galactosidase (SA-β-GAL)^11,12,51,52^. Both persister/diapause-like and senescent cancer cells showed increased lysosomal mass (**Figures 2E, S2D-E**), but only senescent cells had increased SA-β-GAL activity (**Figure 2F** and **S2G**). These results were confirmed in additional cell lines (MCF7, MDA-MB-231, H1299) (**Figures S2F** and **S2G**). Elevated levels of total ROS are another feature of senescent cells. Interestingly, persister/diapause-like cells had total ROS levels comparable to those of proliferating cells, and clearly lower than senescent cells (**Figure 2G** and **S2H**). Another hallmark of senescence is the loss of Lamin B1 (LMNB1), a component of the nuclear lamina. Notably, persister/diapause-like cells had increased levels of LMNB1 compared to proliferating and senescent cells (**Figure 2H** and **S2I**). In summary, persister/diapause-like arrest is characterized by absence of the SASP, high lysosomal content, absence of SA-β-GAL activity, normal ROS levels and retention of LMNB1 (see in **Table 1**).

**Table 1.**
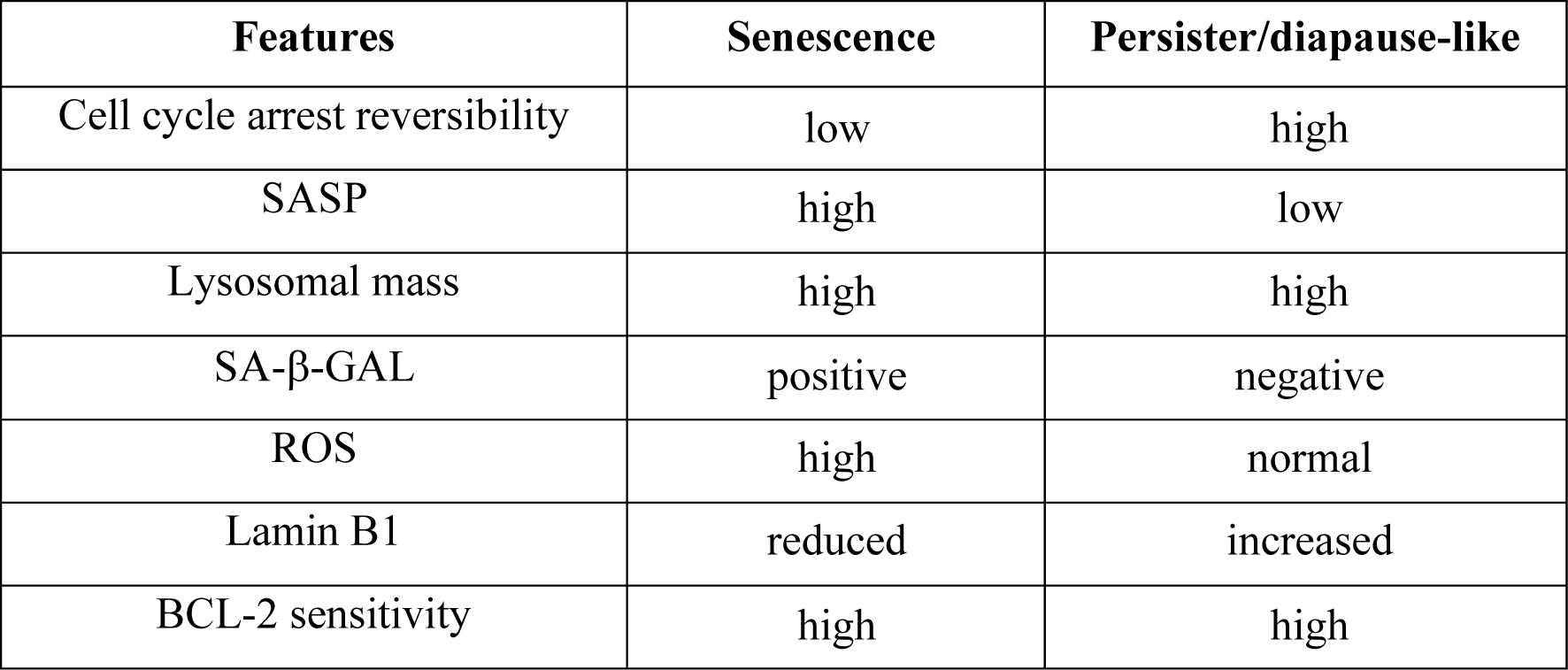
Senescent features in therapy-induced senescence and persistency/diapause-like arrest. Summary of the senescent features analyzed in both TIS and DLCC in A549 and SK-Mel-103 cells.

### Persister/diapause-like cancer cells are sensitive to Bcl-2 family inhibition

One of the most general features of senescent cells is their high sensitivity to inhibitors of the antiapoptotic BCL-2 family proteins and, in particular, to navitoclax, a highly specific inhibitor of BCL-2, BCL-XL (encoded by *BCL2L1* gene) and BCL-W (encoded by *BCL2L2*)^53^. We wondered whether diapause-like cancer cells share this feature with senescent cells. Proliferating, diapause-like and senescent cancer cells were treated with navitoclax for 48h and cell viability was assessed. As expected, senescent cells were sensitive to navitoclax, while proliferating cells were not (**Figures 3A** and **S3A**). Interestingly, persister/diapause-like cancer cells were also sensitive to navitoclax (**Figures 3A** and **S3A**). In general, BCL-XL is the most important antiapoptotic protein for senescent cells^54–58^ and we wondered if this was also the case for diapause-like cells. Cells undergoing INK128-diapause-like arrest (for 7 days) were treated for 5 additional days with siRNA pools targeting BCL-2, BCL-XL and BCL-W (**Figure 3B**). Interestingly, si*BCL-XL* reduced the viability of SK-Mel-103 and A549 under diapause-like arrest (**Figures 3C** and **S3B**), while si*BCL-2* or si*BCL-W* had no effect on the viability of INK128-treated SK-Mel-103 (**Figure 3C**). We also tested if navitoclax could have activity on *in vivo* tumor xenografts treated with a clinical grade mTOR/PI3K dual inhibitor, BEZ235 (also known as dactolisib). We injected mice subcutaneously with SK-Mel-103 cells and when tumors reached approximately 100 mm^3^ in size, we initiated treatment with BEZ235 (to induce the persister/diapause-like state *in vivo*) and/or navitoclax, for 7 additional days (**Figure 3D**). Interestingly, while navitoclax had no effect on tumor growth when used as single agent, it had a significant impact in reducing tumor growth when used in combination with BEZ235 (**Figures 3E** and **S3C**). We conclude that persister/diapause-like cancer cells are hypersensitive to BCL-XL inhibition, a feature that is shared with senescent cells (**Table 1**).

**Figure 3.**
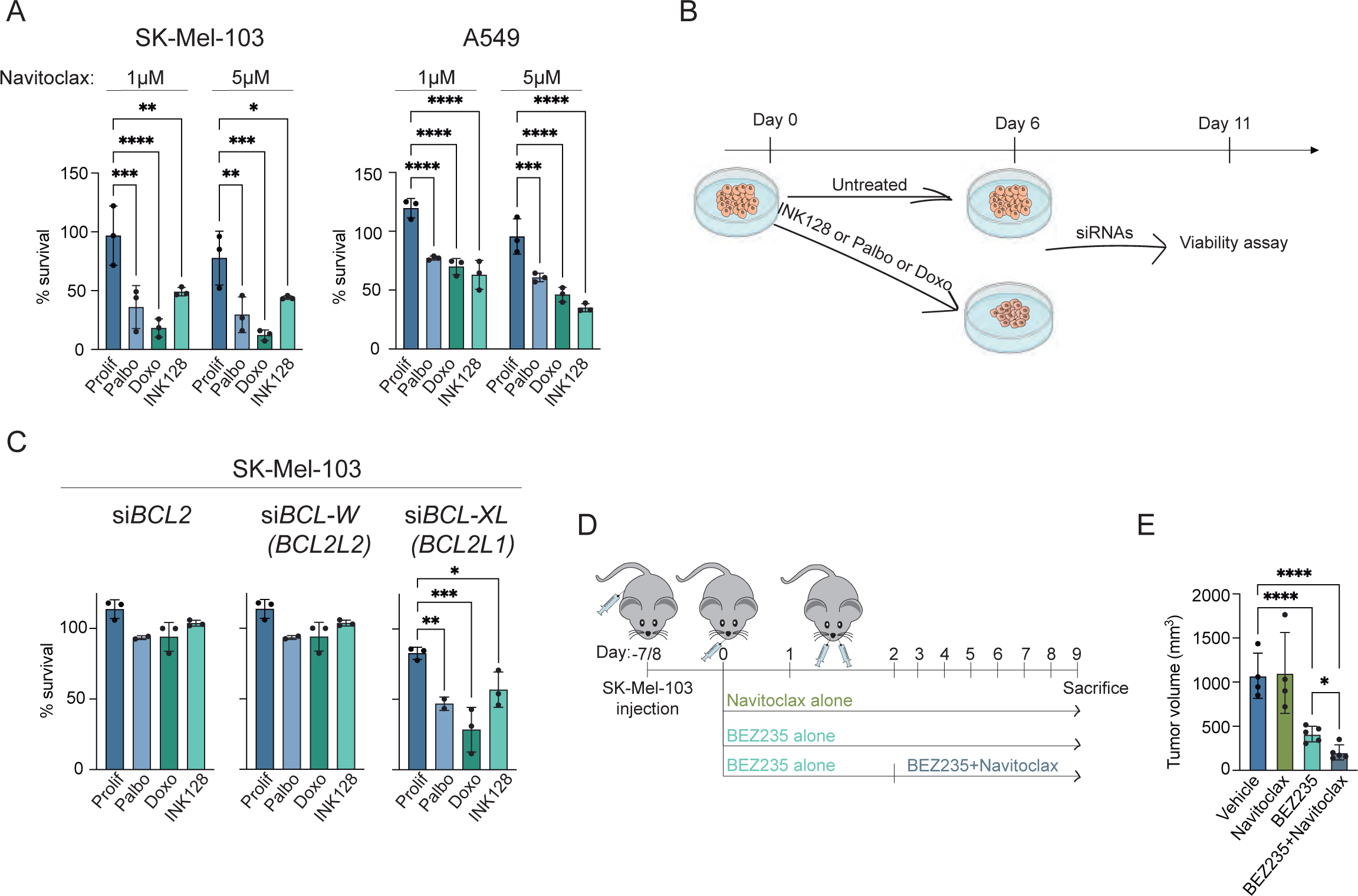
Senescent and persister cancer cells are sensitive to BCL-XL inhibition. (A) Crystal Violet viability assay of proliferating, palbo-treated, doxo-treated and INK128-treated SKMel-103 and A549 after Navitoclax treatment (72hrs) at 5μM and 1μM. % of survival was calculated in comparison to each respective untreated control (n=3). Quantification was done using the SYNERGY HTX Absorbance microplate reader. ***, P < 0.001; **, P < 0.01; two-way ANOVA compared to proliferating controls. (B) Schematic of the protocol used for siRNA transfection and viability assessment. (C) CellTiterGlo viability assay of proliferating, palbo-treated, doxo-treated and INK128-treated SK-Mel-103 cells treated with siRNAs targeting: *BCL2*, *BCL-W* (*BCL2L2*), *BCL-XL* (*BCL2L1*). Viability was assessed 5 days after siRNA transfection (n= 3). Raw data for quantification were acquired by measuring luminescence in a VICTOR Multilabel Plate Reader (Pelkin Elmer). ***, P < 0.001; **, P < 0.01; *, P< 0.05; one-way ANOVA compared to proliferating controls. (D) Schematics of the cancer treatment protocol used in this study. (E) Tumor volume at sacrifice of SK-Mel-103 tumor-bearing animals treated with vehicle, Navitoclax, BEZ235 and BEZ235+Navitoclax (n=4/5). **** P< 0.001; *, P < 0.05; two-way ANOVA analysis considering all the groups and all the days (for full graphs see Figure S4).

### Identification of persister/diapause-like cancer cells-specific survival genes

We reasoned that persister/diapause-like cancer cells may exhibit other vulnerabilities, beyond their sensitivity to BCL-2 family inhibition, that could open novel approaches for tumor control. To identify novel vulnerabilities, we performed an unbiased genetic screen to identify genes whose inactivation is particularly deleterious for diapause arrested cells compared to proliferative cells. Capitalizing on the similarities between diapause-like arrest in cancer cells and embryonic diapause, we performed the primary screening in mouse embryonic stem cells (mESC) and the targets obtained were validated, in a second phase, in human diapause-like cancer cells. In particular, we used mESC carrying a doxycycline-inducible caspase 9 (Cas9) and an exome-wide single guide (sgRNA) library^59^. These cells were treated with INK128 for 5 days to induce embryonic diapause and doxycycline was then administered for 3 more days to induce Cas9/sgRNA editing (**Figure 4A**). A similar procedure was performed on mESC that were not treated with INK128 and therefore correspond to proliferating cells. The relative abundance of sgRNAs (ratio after and before editing) was obtained for diapause cells and for proliferating cells. We reasoned that sgRNAs depleted specifically in diapause cells but not in proliferating cells represented potential genes whose activity was critical specifically for persister/diapause-like cancer cell survival (**Figure 4A**). We prioritized those sgRNA-targeted genes that were depleted by at least two-fold following Cas9 induction (log2 fold change ≤ −1.00). In this manner, we identified 824 genes specific for the survival of diapause mESC, 83 commonly involved in the survival of diapause and proliferating mESC, and 229 specifics for the survival of proliferating mESC (**Figure 4B**).

**Figure 4.**
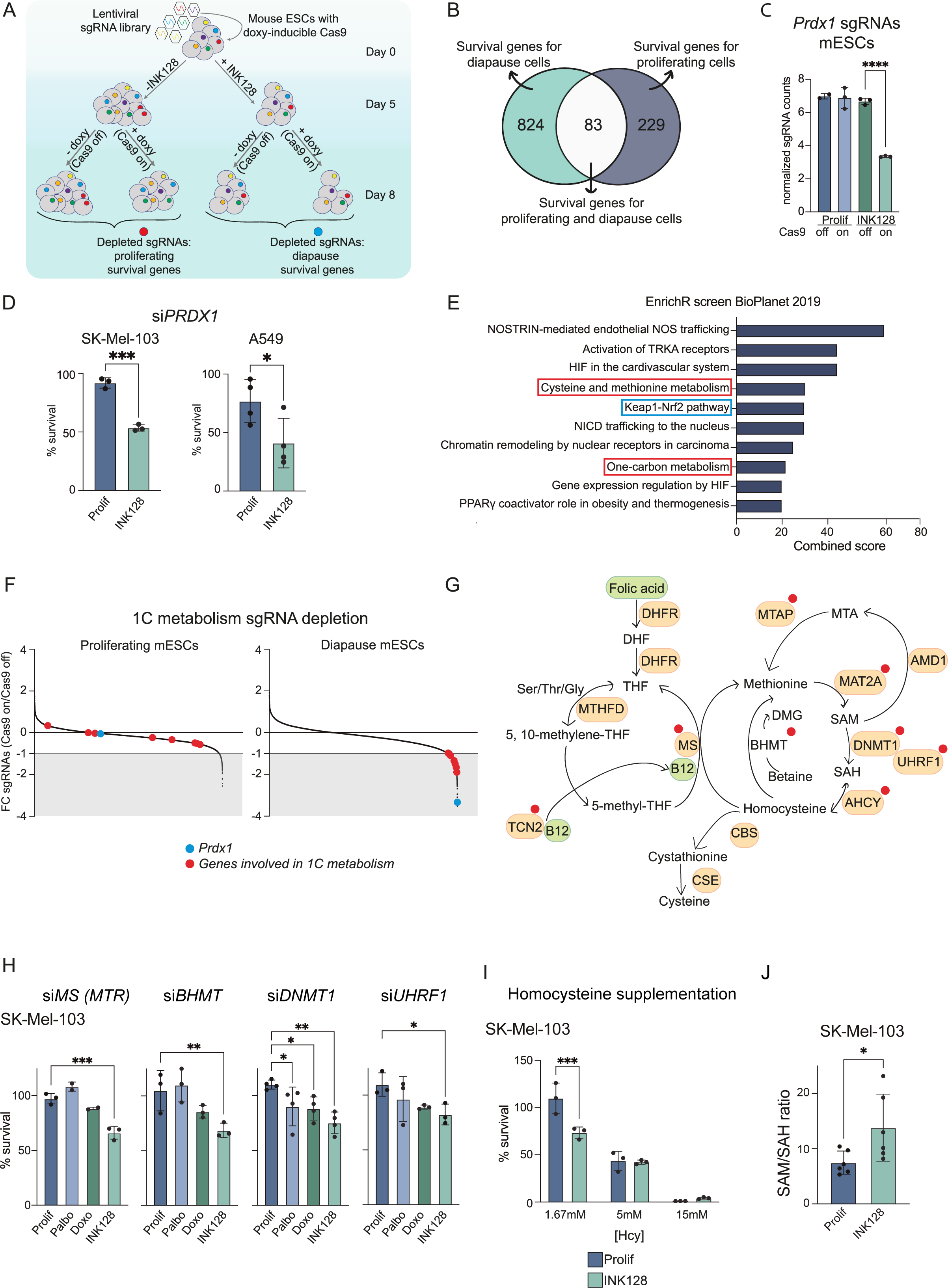
Whole genome CRISPR/Cas9 screening identifies 1C metabolism and DNAme as specific vulnerabilities of persister cells. (A) Schematic of the CRISPR/Cas9 screening experiment used in this study. (B) Venn diagram displaying sgRNAs depleted in proliferating and diapause mouse embryonic stem cells with a log2FC cutoff of −1. (C) *Prdx1* sgRNA counts in proliferating and diapause mESCs untreated (Cas9 off) and treated (Cas9 on) with doxycycline (Proliferating Cas9 off mESCs n=2; all the other conditions n=3). ****, P < 0.0001, one-way ANOVA versus Cas9 off diapause mESC. (D) Crystal violet viability assay of proliferating and INK128-treated SK-Mel-103 transfected with si*PRDX1* for 5 days. % of survival was calculated in comparison to the respective siSCR. Raw data for quantification were acquired using the SYNERGY HTX Absorbance microplate reader. **, P < 0.01; unpaired t test compared to proliferating controls. (E) Top 10 significantly enriched terms from EnrichR BioPlanet 2019 when analyzing the genes found depleted in the diapause mESCs treated with doxycycline. Combined score is performed by the EnrichR software by multiplying the log p-value from the Fisher extact test by the z-score of the deviation from the expected rank. (F) Representation of the Log2FC of all the sgRNAs detected in the screening in proliferating mESC and diapause mESC (fold change of Cas9 on vs Cas9 off for each condition). Red circles show the genes associated to 1C metabolism and the blue circle represents Prdx1, the most depleted gene in the diapause mESC. (G) Schematic of the 1C metabolism. Red circles highlight the genes found as depleted in the screening in diapause mESC. (H) Crystal violet viability assay of proliferating, palbo-treated, doxo-treated and INK128-treated SK-Mel-103 transfected with siMS (MTR), siBHMT, siDNMT1, siUHRF1 or 5 days (n=3 or 4).% of survival was calculated in comparison to the respective siSCR. Raw data for quantification were acquired using the SYNERGY HTX Absorbance microplate reader. ***, P < 0.001; **, P < 0.01; *, P < 0.05; one-way ANOVA compared to proliferating controls. (I) Ratio of SAM/SAH abundance in proliferating and INK128-treated SK-Mel-103 cells. *, P < 0.05; unpaired t test compared to proliferating controls. (J) CellTiterGlo viability assay of proliferating and INK128-treated SK-Mel-103 with supplementation of Homocysteine at 3 different concentrations: 1.67mM, 5mM, 15mM (n=3). Viability was assessed 72hrs after Hcy supplementation. Raw data for quantification were acquired by measuring luminescence in a VICTOR Multilabel Plate Reader (Pelkin Elmer). ***, P < 0.001; two-way ANOVA compared to proliferating controls.

We focused on the 824 genes specific for the survival of diapause cells. Encouragingly, we found that *Prdx1* (encoding PRDX1) was the gene with the highest sgRNA depletion in diapause cells, whereas *Prdx1* sgRNAs were not altered in proliferating mESC (**Figure 4C** and **Table S2**). This suggests that inactivation of *Prdx1* represents a vulnerability specific to the dormant state. Peroxiredoxin-1 (PRDX1) is an activator and downstream effector of NRF2 (encoded by *Nfe2l2*), a transcription factor reported to be critical for tumor dormancy^60^. Interestingly, when *siPRDX1* was tested in human cancer cells (SK-Mel-103 and A549), it also led to a reduction in the survival of persister/diapause-like cells, while it had no effect on proliferating cells (**Figures 4D**). The abundance of *Nfe2l2* sgRNAs was also reduced specifically in diapause mESC (**Figure S4A**), and, again, its downregulation in diapause-like SK-Mel-103 led to reduced survival (**Figure S4B**). These results validated the potential of our screening platform to identify diapause-specific survival genes and confirmed the existence of functional similarities between embryonic diapause and persistency/diapause-like arrest in cancer cells.

To discover novel mechanisms of survival selective to persister/diapause-like cancer cells, we performed pathway analysis of the 824 depleted survival candidate genes using the EnrichR software. This analysis highlighted once again the importance of the KEAP1-NRF2 pathway as a top vulnerability of diapause-like cells (**Figure 4E**). Interestingly one carbon (1C) metabolism was among the most significantly enriched pathways, identified by several independent databases (**Figures 4E** and **S4C**). 1C metabolism, comprised of both the methionine and the folate cycles (**Figure 4G**), shuttles methyl groups to generate substrates for biosynthetic processes including nucleotide biosynthesis, redox balance, and the methylation of both histones and DNA^61^. Interestingly, several other enriched pathways pointed to a diapause dependency on 1C metabolism, such as cysteine and methionine metabolism (which feed 1C metabolism), cobalamin metabolism and vitamin digestion (vitamin B12 is a critical co-factor for the methionine cycle), and the methionine de novo salvage pathway (**Figures 4E** and **S4C**). In particular, diapause mESC presented significant depletion of sgRNAs targeting multiple genes involved in 1C metabolism and DNA methylation (*Bhmt*, *Mtr*, *Mat2a*, *Mtap*, *Tcn2*, *Ahcyl2, Dnmt1, Uhrf1*), while the abundance of the same sgRNAs was not altered in proliferating mESC (**Figure 4F**).

We tested the effect of the downregulation of some of these genes in proliferating, persister and senescent human cancer cells. The inclusion of senescent cells in these analyses was to discriminate whether the potential survival genes represent persister/diapause-like-specific vulnerabilities or are shared between persister and senescent cancer cells. siRNA mediated knockdown of *DNMT1*, *UHRF1*, *BHMT* and *MTR* significantly reduced the survival of INK128-treated SK-Mel-103, but had no effect on proliferating cancer cells. Senescent cancer cells were not sensitive to the knockdown of these genes, except in the case of si*DNMT1*, which produced a milder albeit significant reduction in survival (**Figure 4H**). Although most of the hits identified in diapause mESC were validated in human SK-Mel-103 undergoing persistency/diapause-like arrest, there were some exceptions. For example, inhibition of *AHCYL2* had no effect on either persister/diapause-like, senescent or proliferating SK-Mel-103 cells (**Figure S4D**), perhaps due to compensation by paralogs AHCY and AHCYL1. In the case of A549 cells, we could only validate a persister/diapause-like-specific survival role for *DNMT1* and *UHRF1* (**Figure S4E**). Interestingly, persister/diapause-like cancer cells were hypersensitive to homocysteine (Hcy) compared to proliferating cells (**Figure 4I**). At high concentrations, Hcy is converted into S-adenosyl-homocysteine (SAH), a strong competitor of the universal methyl-donor S-adenosyl-methionine (SAM), thereby compromising methyl-transferase reactions^62,63^.

To substantiate the role of 1C metabolism in persister/diapause-like cells, we performed a targeted metabolomic analysis. The SAM/SAH ratio is defined as the “methylation index” because it indicates the methylation capacity of a cell. We found that the SAM/SAH ratio was significantly increased in persister SK-Mel-103 cells compared to proliferating cells (**Figure 4J**), suggesting a higher methylation demand. We also found an increase in the ratios between methionine and its precursors 5-methyl-THF and betaine, again suggesting an elevated demand for methionine generation (**Figure S4F**). The deviation of homocysteine (Hcy) out of the methionine cycle and into the formation of cysteine (Cys) is critical for the synthesis of glutathione and thereby for anti-oxidant capacity. Interestingly, we also observed depletion of Cys in persister cells (**Figure S4G**). This is consistent with a high methylation demand and a lower deviation of Hcy into Cys.

Collectively, the above data suggest that persister/diapause-like cells are highly dependent on 1C metabolism as compared to proliferating or senescent cancer cells.

### Persister/diapause-like cancer cells present H4K20me3 heterochromatic foci

Based on the importance of 1C metabolism for histone methylation reactions, we looked at the levels of selected histone methylation marks. In particular, we focused on the canonical heterochromatin marks H3K9me3 and H4K20me3^64–67^. Redistribution of these marks has been implicated in other types of cell-cycle arrest, particularly in quiescence^68,69^ and senescence^14,16,70^, and we hypothesized that they could underlie the repression of the inflammatory programs in persister cells. Immunoblot analysis revealed an increase in H3K9me3 and H4K20me3 in INK128-treated SK-Mel-103 and A549 cells compared to proliferating cells (**Figures 5A-B** and **S5A-B**). As reported^70^, we also detected an increase in the global levels of H4K20me3 in therapy-induced senescent cells, while the total or relative levels of H3K9me3 were not increased (**Figures 5A-B** and **S5A-B**).

**Figure 5.**
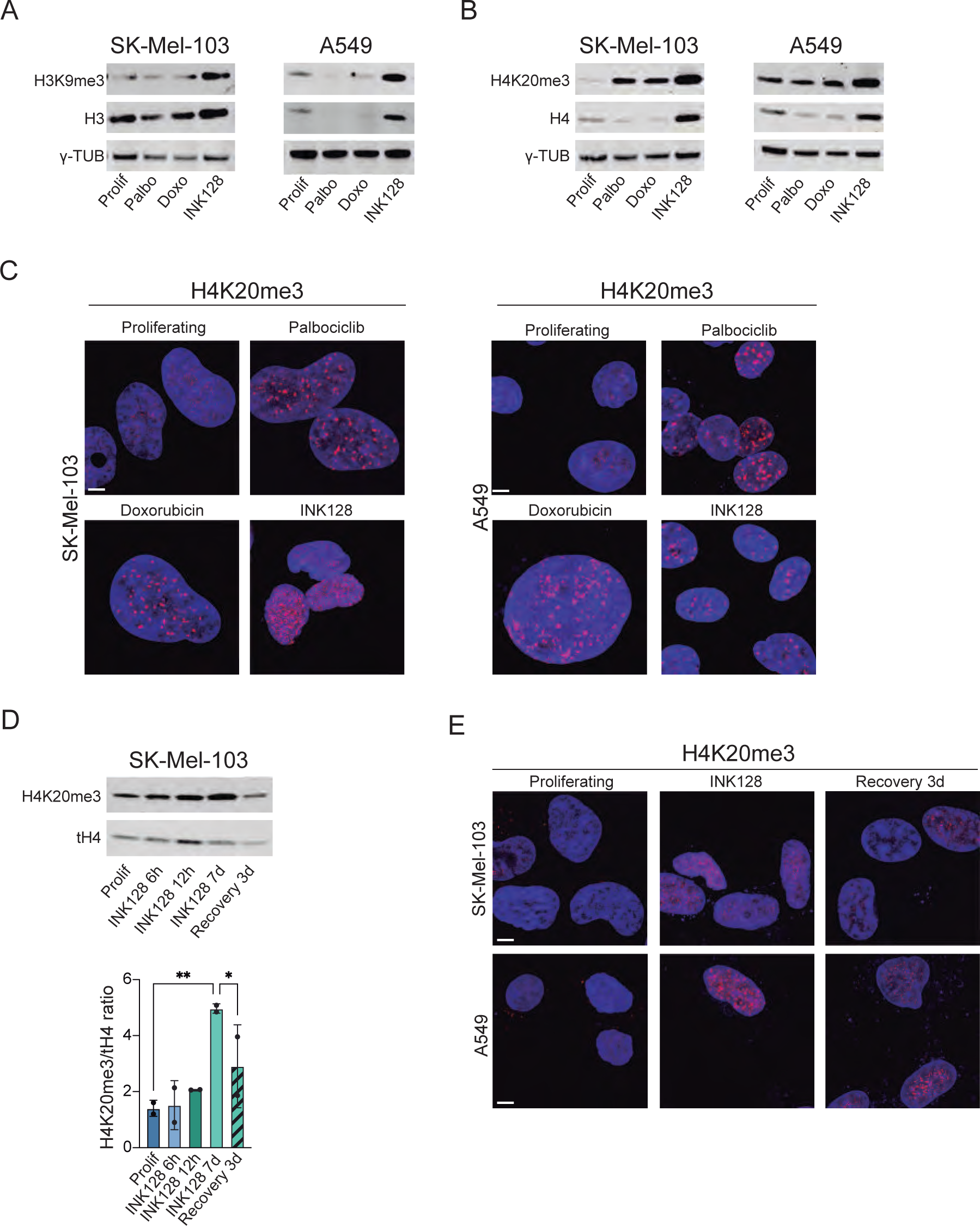
Persister/diapause-like cells present H4K20me3 heterochromatin foci. (A) H3K9me3, total H3 and γ-Tubulin protein levels in proliferating, palbo-treated, doxo-treated, INK128-treated SK-Mel-103 and A549 cells. A representative immunoblot is shown. Quantification in Figure S6. (B) H4K20me3, total H4 and α-Tubulin protein levels in proliferating, palbo-treated, doxo-treated, INK128-treated SK-Mel-103 and A549 cells. A representative immunoblot is shown. (C) Immunostaining of H4K20me3 (red) and dapi (blue) in proliferating, palbo-treated, doxo-treated and INK128-treated SK-Mel-103 and A549 cells. Scale bar is 5μm. Quantification in Figure S6. (D) H4K20me3, total H4 and γ-Tubulin protein levels in proliferating, INK128-treated (6hrs, 12hrs, 7days) and INK128-recovery (3 days of withdrawal) SK-Mel-103 cells. A representative immunoblot (left) and quantification (right) from n=2 is shown. H4K20me3 levels have been normalized on total H4 levels. **, P < 0.01; *, P < 0,05; one-way ANOVA compared to INK128 7 days. (E) Immunostaining of H4K20me3 (red) and dapi (blue) of SK-Mel-103 and A549 cells in the following conditions: proliferating, INK128-treated, and cells after 3 days from INK128 withdrawal. Representative images are shown. Scale bar is 5μm. Quantification in Figure S6.

The elevated levels of H3K9me3 in drug tolerant persistency have been reported and linked to the silencing of repeated elements^25^. In contrast, H4K20me3 remains unexplored in the context of persistency/diapause-like arrest in cancer cells. Immunofluorescence staining revealed the formation of H4K20me3 foci in persister/diapause-like and senescent cells (**Figures 5C** and **S5C**). Interestingly, when persister cells resumed proliferation upon withdrawal of INK128, H4K20me3 levels returned to the same levels as proliferating cells, as detected by immunoblot and immunofluorescence (**Figures 5D-E** and **S5D**). This reinforces the association between persistency/diapause-like arrest and high H4K20me3, and also suggests the reversibility of this epigenetic mark.

### H4K20me3 at promoters of inflammatory genes distinguishes persistency/diapause-like arrest and senescence

To gain a more comprehensive understanding of the role of H3K9me3 and H4K20me3 in persister/diapause-like cancer cells, we performed Cleavage Under Targets and Tagmentation (CUT&Tag) for these epigenetic marks in proliferating, senescent and persister/diapause-like SK-Mel-103 and A549 cells. Pairwise Pearson correlation indicated that the biological replicates (n=3) per condition clustered closely together, indicating the reproducibility of the assay (**Figure S6A**). To analyse the data, we used ChromHMM (Chromatin Hidden Markov Model)^71^ that identifies in an unbiased manner recurrent configurations (called “states”) of a given set of epigenetic marks (in our case H3K9me3 and H4K20me3) across a set of samples. In the case of senescent cells, we observed notable changes in the abundance of H3K9me3 compared to proliferating cells (**Figure 6A**). These changes consisted in losses (states 1 and 2) and gains (states 4 and 6) in H3K9me3, compared to proliferating cells. In relation to this, previous reports have described remarkable changes in the localization of H3K9me3 associated with the loss of Lamin B1^14^. In contrast, persister/diapause-like cells did not show any alteration in the abundance of H3K9me3 across the different states defined by ChromHMM and compared to proliferating cells (**Figure 6A**).

**Figure 6.**
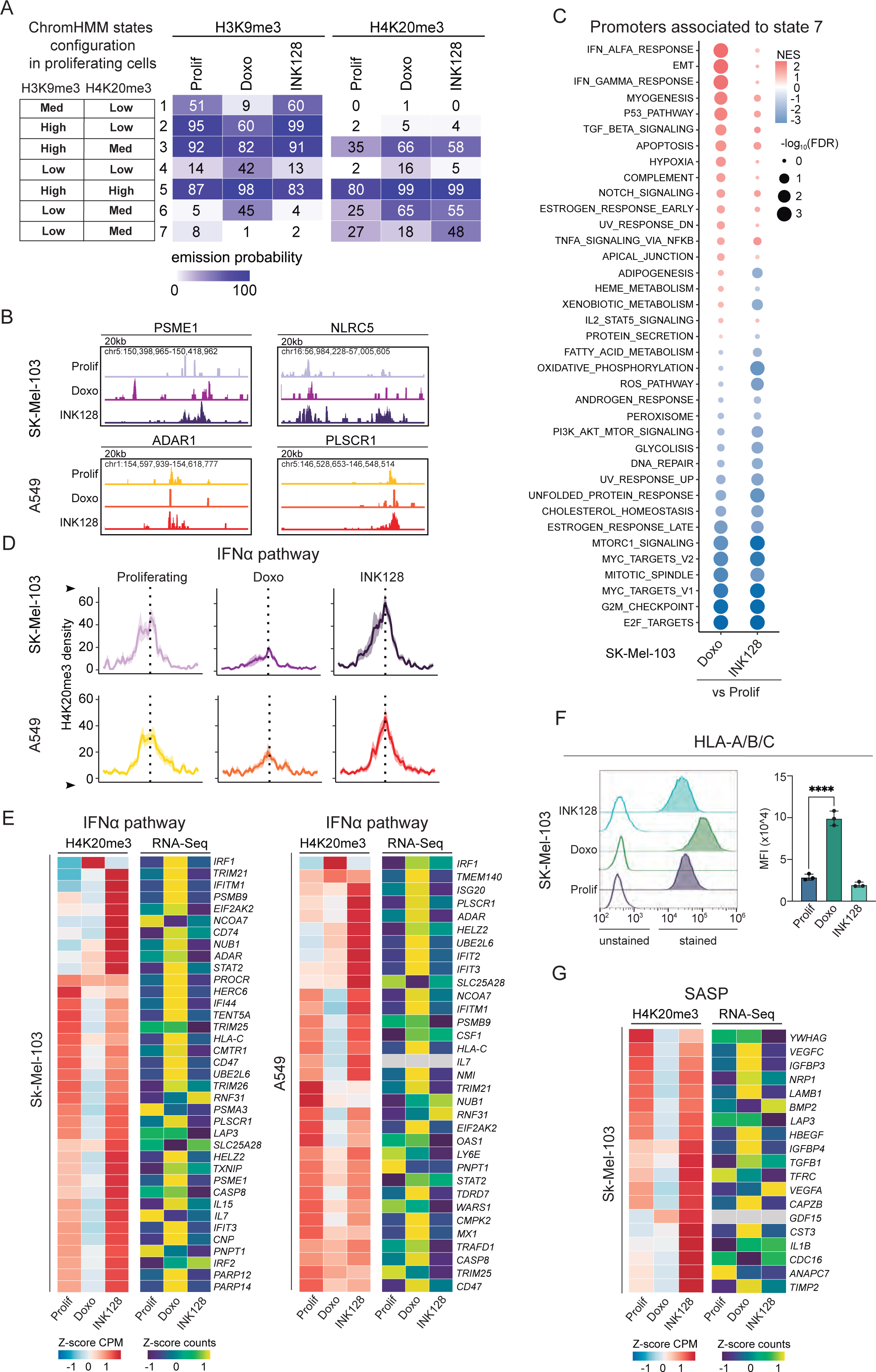
H4K20me3 is differentially associated to IFN and SASP-associated promoter regions in senescent and persister cancer cells. (A) Heatmap summarizing the ChromHMM emission matrix of H3K9me3 and H4K20me3 over the ChromHMM states 1-7 in proliferating, doxo-treated, INK128-treated SK-Mel-103 and A549 (n=3). (B) Bigwigs of promoter regions that exhibit increased H4K20me3 signal in chromatin from INK128-treated compared to proliferating and doxo-treated SK-Mel-103 and A549 cells. (C) Dotplots of Pre-ranked GSEA associated to the ChromHMM state 7 of SK-Mel-103 and A549 cells, where genes were ranked according to their fold change between treated and controls cells. Normalised Enrichment Score (NES) is colour-coded, while statistical significance (FDR: False Discovery Rate) is encoded in the size of the dots. (D) H4K20me3 signal over promoters of genes in the GSEA hallmark gene sets related to IFNα pathway that intersect with ChromHMM state 7 in proliferating, doxo-treated, INK128-treated A549 and SK-Mel-103 cells. (E) Heatmaps plotting the levels of H4K20me3 over promoters in gene set ‘Hallmark interferon alpha response’ that intersect with ChromHMM state 7 in proliferating, doxo-treated, INK128-treated SK-Mel-103 and A549 cells. This is accompanied by a heatmap plotting the normalized RNA-Seq counts for these genes, averaging replicates. (F) Flow cytometry analysis of human (HLA-A/B/C) MHC-I expression in proliferating, palbo-treated, doxo-treated and INK128-treated SK-Mel-103 cells. Representative histograms showing the fluorescence signal of each stained sample and its unstained control (uncolored histogram). Quantification of the MFI after autofluorescence subtraction (n=3). ****, P < 0.0001; one-way ANOVA compared to proliferating control cells. (G) Heatmaps plotting the levels of H4K20me3 over promoters of SASP genes (previously defined as TIS SK-Mel-103 SASP) that intersect with ChromHMM state 7 in proliferating, doxo-treated and INK128-treated SK-Mel-103 cells. This is accompanied by a heatmap plotting the normalized RNASeq counts for these genes, averaging replicates.

Analysis of the abundance of H4K20me3, on the other hand, revealed a number of chromatin states that gained this mark in persister/diapause-like cancer cells and/or therapy-induced senescence. In particular, states 3, 5 and 6 were enriched in H4K20me3 in both conditions. Notably, state 7 was the only one in which persister/diapause-like cells behaved differently from both proliferating and senescent cells. In particular, this state was enriched in H4K20me3 in persister/diapause-like cells relative to the other two cell groups (**Figure 6A**). We then interrogated the composition of genomic features in the above states and state 7 was the only one that contained a large proportion of promoter regions, while the remaining states were mainly composed of intergenic and intronic regions (**Figure S6B**). For illustrative purposes, we show two examples for each cell line (SK-Mel-103 and A549) of promoters that gain H4K20me3 selectively in persister/diapause-like cancer cells compared to therapy-induced senescent or proliferating cells (**Figure 6B**). We next interrogated whether the promoters in state 7 were enriched in specific biological pathways. Remarkably, among the pathways enriched in state 7, we identified some related to secretory and inflammatory processes that were highly expressed in therapy-induced senescence, but not in persistency/diapause-like arrest (**Figure 6C** and **S6C**). These data suggest that the differences in the transcriptional profiles between senescent and persister cancer cells can be ascribed to H4K20me3 distribution over the promoter regions of genes in pathways that are mostly related to inflammation.

Interestingly, the IFNα and the IFNγ pathways were among the most differentially regulated pathways in senescent and persister cells compared to proliferative cells (**Figure 6C** and **S6C**). In senescent cells, interferon-regulated genes lacked H4K20me3 at their promoters and were highly expressed, whereas in persister cells, the same genes were enriched for H4K20me3 at their promoters and were not expressed (**Figures 6D-E** and **S6D**). Interferon signaling is a major inducer of antigen presentation by the MHC-I complex in senescent cells^72,73^. We hypothesized that the lack of IFN activation in persister cells might result in the absence of MHC-I expression on the plasma membrane surface. We measured MHC-I levels on the plasma membrane of proliferating, senescent and persister cells. As expected, both senescent SK-Mel-103 and A549 cells upregulated MHC-I (as detected by an antibody against HLA-A, B and C) (**Figures 6F** and **S6D**). In contrast to senescent cells and in agreement with the H4K20me3 epigenetic data, INK-128-treated cells did not show upregulation of MHC-I compared to proliferating cancer cells (**Figures 6F** and **S6D**).

Finally, we wondered if the absence of SASP in persister/diapause-like cells could be associated with presence or absence of H4K20me3 at the promoter regions of SASP genes. Indeed, all the SASP promoter regions of senescent cells lost H4K20me3, which is consistent with their transcriptional activation. In contrast, persister/diapause-like cells retained or gained the repressive mark H4K20me3 at SASP promoters (**Figures 6G** and **S6E**). Collectively, these data reveal that H4K20me3 is associated with the expression of the SASP: it is lost in senescence where the SASP is strongly upregulated, and retained or increased in persistency/diapause-like arrest where the SASP is absent.

## Discussion

In this study, we characterize a model of persister/diapause-like cancer cells (DLCC) driven by mTOR/PI3K dual inhibition through INK128 treatment. We show that INK128-treated cancer cells present features of both drug-tolerant persistency and tumor dormancy, and their transcriptional profile resembles embryonic diapause. This is consistent with emerging evidences highlighting the similarity between these three states of reversible arrest, namely, dormancy, persistency and diapause^26,27,74^.

We define a panel of distinct and common features between senescence and persister/diapause-like arrest in cancer cells (**Table 1**), which could be useful for future research. Interestingly, we found a number of features that were differential between these two types of cell cycle arrest. In particular, persister cells lack high levels of reactive oxygen species (ROS) and do not lose LMNB1 levels. An important difference between persister and senescent cells derives from the finding that persister cells lack the senescence-associated secretory phenotype (SASP) and the transcriptional programs associated to interferon signaling, including the absence of MHC-I upregulation, which could contribute to their escape from immunosurveillance.

Among the common traits between senescent and persister/diapause-like cancer cells is the expansion of the lysosomal compartment, which is likely attributable to an increased autophagic activity^11,75^. Interestingly, the senescence-associated β-galactosidase activity (SA-β-GAL) which is a lysosomal enzyme detected at a sub-optimal pH in senescent cells was absent persister cells, therefore disconnecting lysosomal expansion from SA-β-GAL. This is consistent with previous reports noting that SA-β-GAL activity in senescent cells is extinguished upon inhibition of mTOR activity^76,77^ Transcriptomic analysis revealed upregulation of the TGFβ and the WNT/βCAT pathways in both senescent and persister/diapause-like cancer cells; these pathways have been reported to be upregulated during tumor dormancy *in vivo*^46,49^. Another interesting trait shared by senescent and persister cells is their sensitivity to the BCL-2 family inhibitor navitoclax, which in both types of cell-cycle arrest was due to the specific inhibition of *BCL-XL*. To support this observation *in vivo*, we treated mice carrying human tumor xenografts with an mTOR/PI3K inhibitor and/or with navitoclax. Interestingly, while navitoclax had no effect on tumor growth as a single agent, it significantly synergized with the mTOR/PI3K inhibitor. This suggests that persister/diapause-like cancer cells are pharmacologically targetable *in vivo* with navitoclax.

To identify additional vulnerabilities of persister/diapause-like cancer cells, we performed a CRISPR-Cas9 screen in mouse embryonic stem cells (mESC), thus leveraging the similarities between embryonic diapause and diapause-like arrest in cancer cells. We identified *Prdx1* as the most depleted gene in diapause mESC, and we validated *PRDX1* and its upstream transcriptional activator *NRF2* as survival genes also in our human cancer models of persistency/diapause-like arrest. PRDX1 is one of the mediators of ROS detoxification by the master regulator NRF2^78,79^, and NRF2 has been involved in the survival of dormant cells^60^. Pathway analysis of depleted genes specific to the diapause population highlighted a possible contribution of one-carbon (1C) metabolism and DNA methylation in diapause-like cancer cell survival. The 1C-metabolism is important for providing methyl groups in the form of S-adenosyl-methionine (SAM), which is used for biosynthetic processes and for the methylation of both histones and DNA^61^. In particular, we identified as essential genes for persister SK-Mel-103 cells the enzymes *BHMT* and *MTR*, both involved in separate reactions that generate methionine. In addition, we identified *DNMT1* and *UHRF1* as critical for the survival of persister/diapause-like SK-Mel-103 and A549. Of note, UHRF1 is a binding partner and a critical cofactor of DNA methyltransferase DNMT1^67,80,81^. In support of the importance of methylation reactions, persister cells presented increased SAM/SAH ratios, which are indicative of higher methylation capacity. The deviation of homocysteine (Hcy) out of the methionine cycle and into the formation of cysteine (Cys) is critical for the synthesis of glutathione and thereby for anti-oxidant capacity. Interestingly, we detected decreased levels of cysteine in persister SK-Mel-103 cells compared to proliferating cells, which we interpret as a reflection of the high methylation demand and, consequently, lower deviation of Hcy towards Cys. Interestingly, diapause-like arrested cells were hypersensitive to high concentrations of Hcy compared to proliferative cells. This may be explained by the known capacity of high Hcy to cause both oxidative damage^82^ and inhibition of methyltransferases (via the accumulation of S-adenosyl-homocysteine)^62^. Together these data suggest that 1C metabolism plays an important role in persister/diapause-like survival, which could be ascribed to its function in epigenetic regulation and anti-oxidant protection.

Based on the importance of 1C metabolism for histone methylation reactions, we looked at the levels of selected histone methylation marks. In particular, we focused on the canonical heterochromatin marks H3K9me3 and H4K20me3^64–67^, which have been implicated in quiescence^68,69^ and senescence^14,16,70^. We detected upregulation of both H3K9me3 and H4K20me3 in persister/diapause-like cancer cells by immunoblot. While elevated H3K9me3 in models of drug-tolerant persistency has already been reported^25^, to our knowledge, there is no information of H4K20me3 in persister or diapause-like cancer cells. Interestingly, H4K20me3 foci were increased in both senescent and persister cells compared to proliferating cells, and the increased levels of this mark were lost from persister/diapause-like cells upon INK128 withdrawal, suggesting a strong association of H4K20me3 with persistency/diapause-like arrest.

To further characterize the role of H3K9me3 and H4K20me3 in senescent and persister/diapause-like SK-Mel-103 and A549 cancer cells, we performed CUT&Tag. These data revealed two important differences between senescent and persister cells. Regarding H3K9me3, this mark was largely unchanged in persister cells compared to proliferating cells, while in senescent cells it underwent a partial re-localization that mostly affected intergenic and intronic elements. Re-localization of H3K9me3 in senescent cells does not have a clear impact on the transcriptional profile of senescent cells^83^ and it has been connected to the loss of Lamin B1^14^. In this regard, it is relevant our observation that persister/diapause-like cells do not present loss of Lamin B1.

Regarding the heterochromatic mark H4K20me3, we observed a different behavior in intergenic/intronic regions compared to promoter regions. In the case of intergenic/intronic regions, there was an increase in the abundance of this epigenetic mark observable both in senescent and diapause-like cells, compared to proliferating cells. This elevation may be connected with the emergence of immunofluorescence-detectable H4K20me3 foci in senescent and diapause-like cells. Interestingly, a group of promoter regions with low levels of H4K20me3 in proliferating cells presented a discrepant behaviour between senescent and diapause-like cells: senescent cells loss H4K20me3 at this group of promoters while diapause-like cells elevated their levels of H4K20me3. Although H4K20me3 has been typically associated with constitutive heterochromatin, it has been reported that it could also be located in promoters and consequently have a role in silencing gene expression^84,85^. When we analyzed the biological functions associated to the differential binding of H4K20me3 to promoters in persister/diapause-like and senescent cells, they turned out to be related to inflammatory pathways, notably interferon-regulated genes and SASP genes. The loss of H4K20me3 in the promoters of inflammatory genes may underlie, at least in part, the upregulation of these pathways in senescent cells. Conversely, the gain of H4K20me3 provides an epigenetic explanation for the lack of SASP and IFN gene expression in persister/diapause-like cells. Interestingly, the lack of SASP expression in DTPs was confirmed also by the analysis of a published dataset of colorectal cancer DTP cells, further supporting the idea that the INK128-model of drug tolerant persister cells recapitulate general features of drug-tolerant persistency.

In summary, we provide a characterization of distinct and shared features between senescent and persister cancer cells. Importantly, we used a CRISPR/Cas9-based screen to identify actionable vulnerabilities of persister/diapause-like cells; the results of this screen prompted us to make epigenetic insights into the regulation of the SASP and the “sterile” inflammatory state of the persistency/diapause-like arrest. These findings, including our conserved persister/diapause-like transcriptional signature and the unique vulnerabilities that we have identified, could be instrumental for the study of these subpopulations of cells in post-therapy tumors *in vivo*. We speculate that varying proportions of senescent and persister cells could influence the microenvironment, the immune infiltration of the tumor and the likelihood of relapse. Further studies are warranted to understand the interplay between both states and how each contribute to both the minimal residual disease and tumor relapse.

## Acknowledgements

We are grateful to the IRB Functional Genomics Unit and CRG/CNAG for library preparation and genomic sequencing, and to the IRB microscopy unit. V.R. was recipient of a predoctoral contract from H2020-MSCA-ITN-2018 HealthAge Program. M.K. was funded by the Barcelona Institute of Science and technology (BIST) and Asociación Española Contra el Cáncer (AECC; POSTD18020SERR) and supported by the European Molecular Biology Organization (EMBO). Work in the laboratory of M.S. was funded by the IRB and by a Coordinated-AECC grant (PRYCO211023SERR).

## Author Contributions

V.R conceptualized, designed, performed and analyzed all experiments, contributed to bioinformatic data analysis and co-wrote the manuscript. L.R. performed bioinformatic analysis of the RNA-seq and CUT&Tag data and helped with the interpretation. M.K. helped with the design of the genetic screening, supervised the study, helped with the interpretation of the data and co-wrote the manuscript. C.S.O.A. performed bioinformatic analysis of the RNA-seq and the CRISPR/Cas9 screening data and helped with the interpretation. J.C., O.Y helped design and perform the metabolomic experiment. A.E.L. helped in the generation of the CUT&Tag samples. G.F. supervised the study, secured funding, analyzed the data, and co-wrote the manuscript. M.S. conceptualized, designed and supervised the study, secured funding, analyzed the data, and co-wrote the manuscript. All authors discussed the results and commented on the manuscript.

## Declaration of Interests

M.K. has ongoing or completed research contracts with Galapagos NV, Rejuveron Senescence Therapeutics, and mesoestetic®. M.S. is shareholder of Senolytic Therapeutics, Inc., Life Biosciences, Inc., Rejuveron Senescence Therapeutics, AG, and Altos Labs, Inc. The funders had no role in study design, data collection and analysis, decision to publish, or preparation of the manuscript.

## Data availability

Generated RNA-sequencing datasets have been deposited in GEO under accession number GSE246690. Generated sgRNA-sequencing datasets have been deposited in GEO under accession number GSE251714. Generated CUT&Tag-sequencing datasets have been deposited in GEO under accession number GSE251660.

---

**Figure S1.**
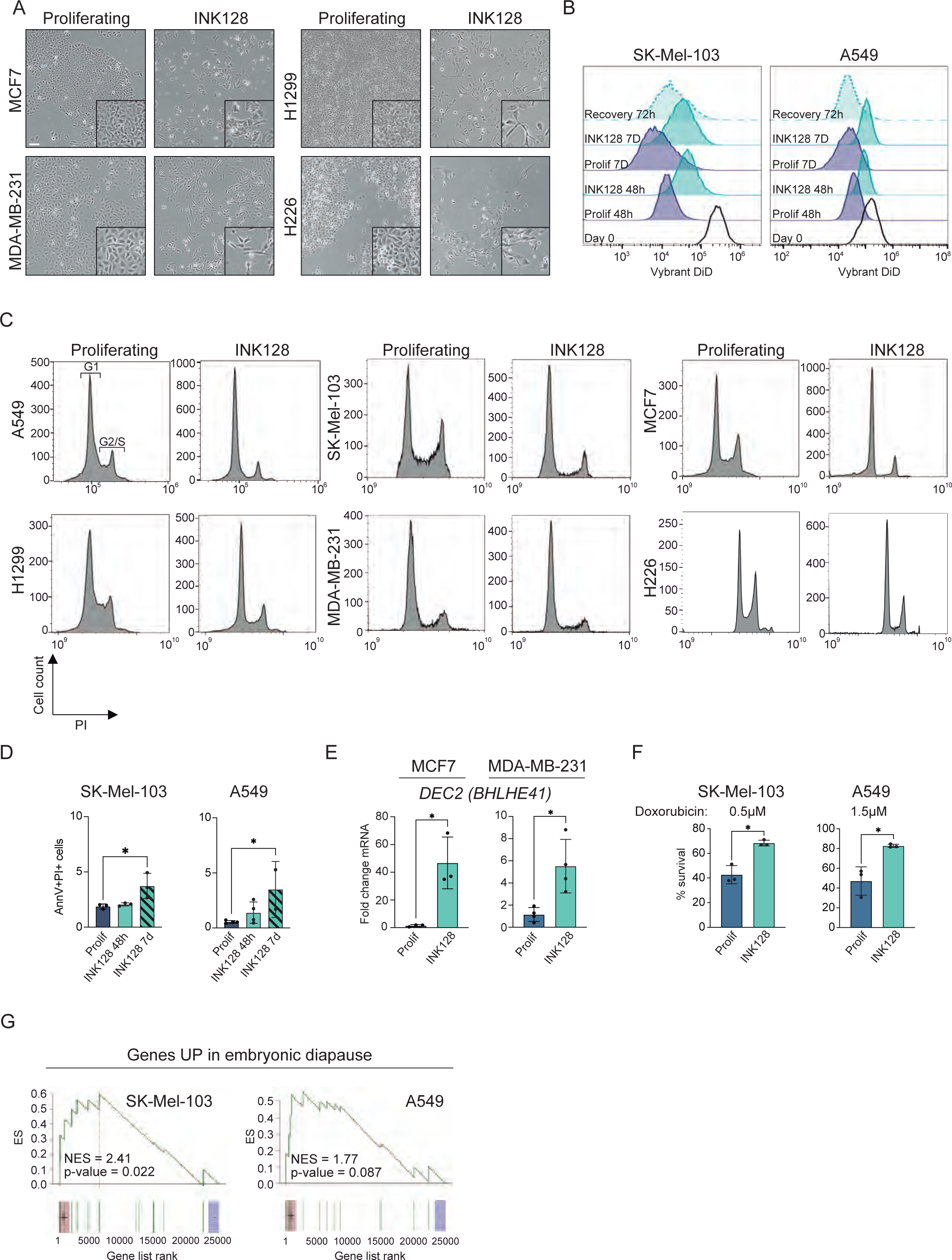
INK128-treated cells undergo a reversible cell cycle arrest with features of drug-tolerant persistency and embryonic diapause. (A) Brightfield pictures of MCF7, MDA-Mb-231, H1299, H226 cells untreated (proliferating) and treated with INK128 (100μM) for 7 days. Scale bar is 100μm. (B) Flow cytometry analysis of label retention experiments performed using the dye Vybrant DiD (n=3). Representative histograms showing the fluorescence signal of each stained sample. (C) Cell cycle flow cytometry analysis through propidium iodide (PI) staining of SK-Mel-103, A549, MCF7, MDA-MB231, H1299, H226 cells untreated (proliferating) and treated with INK128 for 7 days. (D) Flow cytometry analysis of AnnexinV/PI in A549 and SK-Mel-103 cells in the following conditions: proliferating cells, INK128-treated cells after 48 hours, INK128-treated cells after 7 days. n=3, SK-Mel-103, n=4 A549. *, P < 0.05; one-way ANOVA compared to proliferating controls. (E) mRNA expression levels of *DEC2* (*BHLHE41*) in untreated (proliferating) and INK128-treated MCF7 and MDA-MB-231, measured by qRT-PCR (relative to the average expression of housekeeping genes *ACTB* and *GAPDH*) (n=3 MCF7; n=4 MDA-MB-231). *, P < 0.05; unpaired Student’s t test compared to proliferating controls. (F) Crystal violet quantification of proliferating and INK128-treated SK-Mel-103 and A549 cells treated with Doxorubicin at 0.5μM and 1.5μM for 72h. % of survival was calculated in comparison to the respective untreated control (n= 3). Quantification was done using the SYNERGY HTX Absorbance microplate reader. *, P< 0.05; unpaired Student’s t test, compared to proliferating controls. (G) GSEA of the signature “Embryonic_diapause_up”, in INK128-treated versus proliferating SK-Mel-103 and A549 cells. Normalised Enrichment Score (NES) is colour-coded, while statistical significance (FDR: False Discovery Rate) is encoded in the size of the dots.

**Figure S2.**
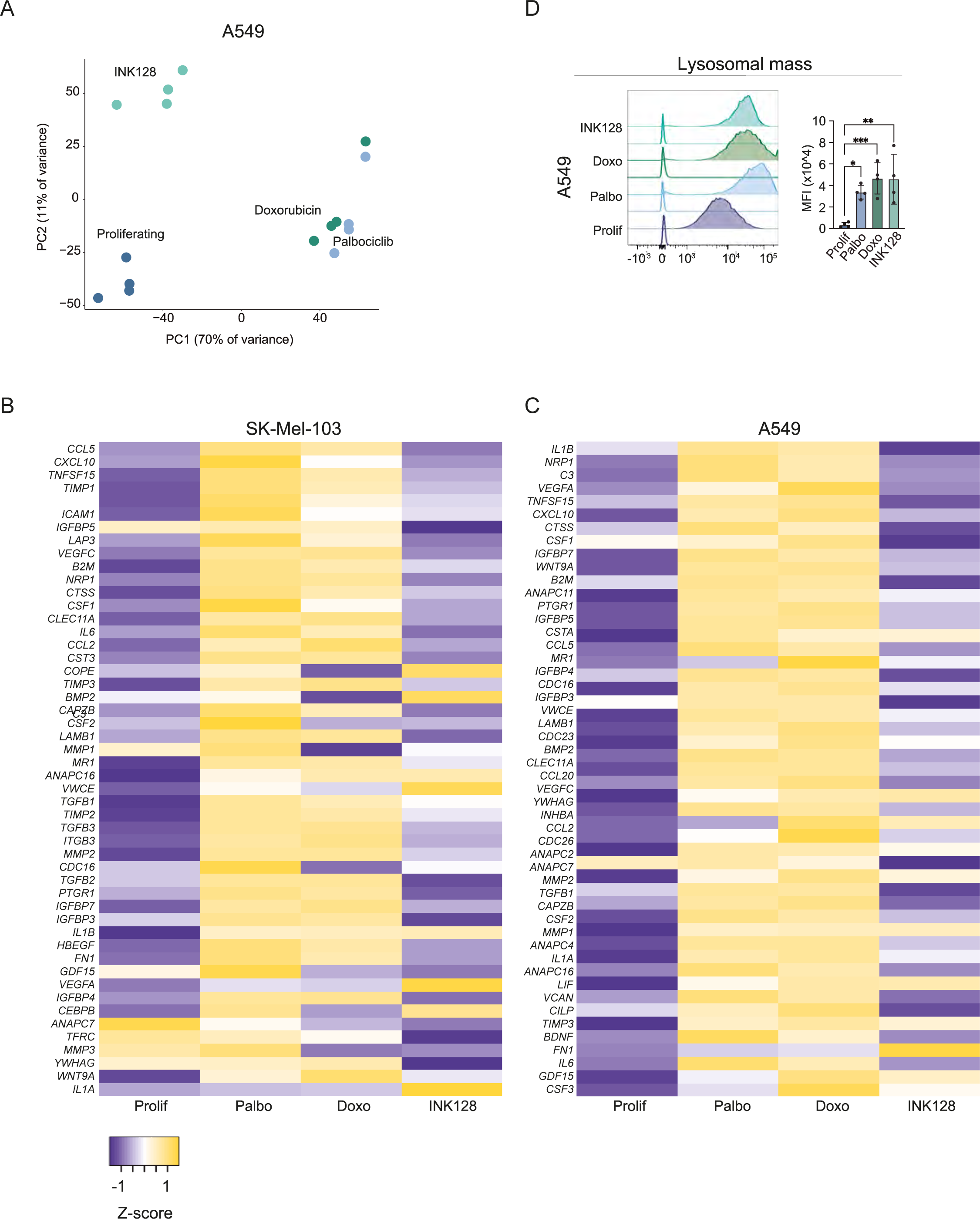
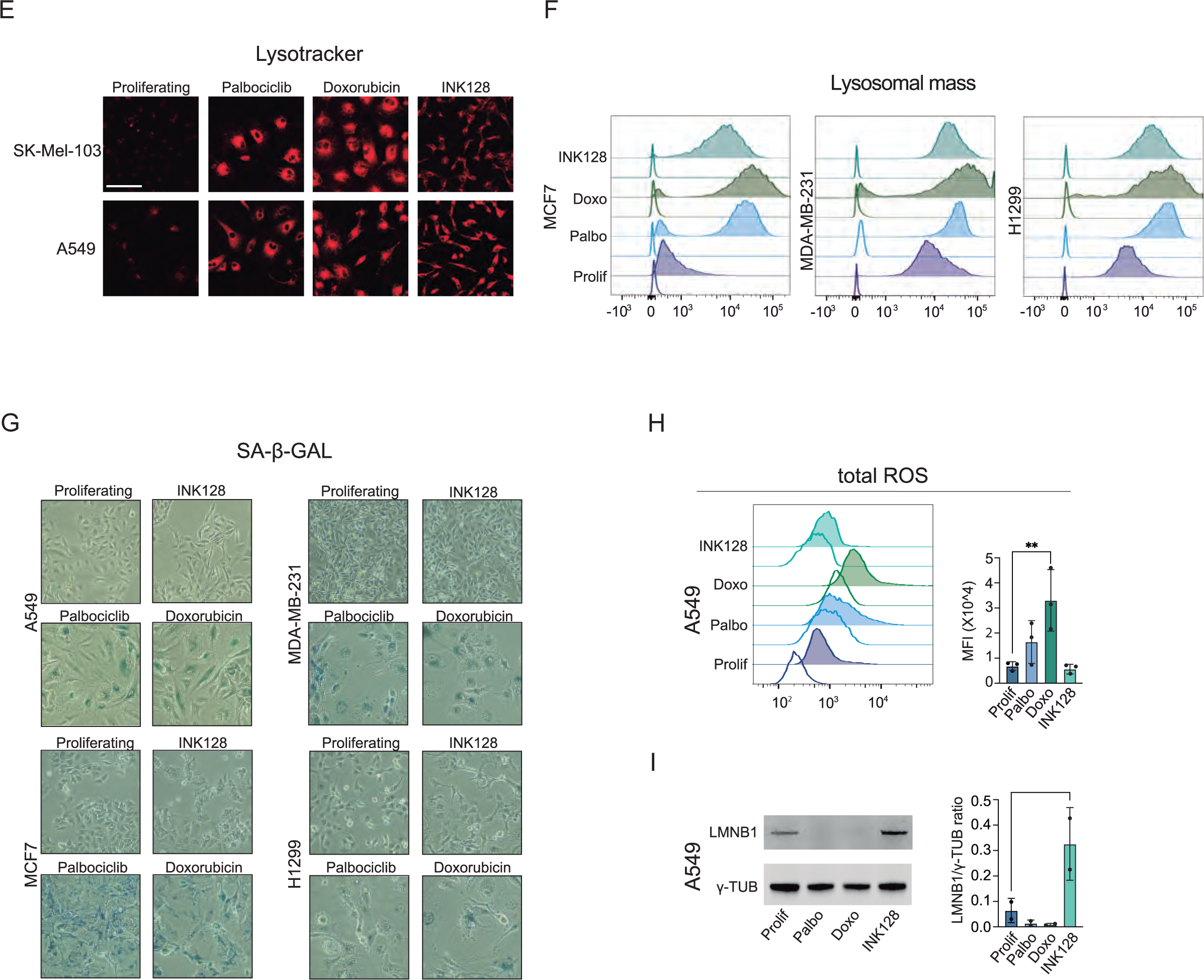
Hallmarks of senescence in persister/diapause-like cancer cells. (A) Principal Component Analysis (PCA) of proliferating, palbociclib-treated, doxorubicin-treated and INK128-treated A549 cells (n=4). (B-C) Row-normalized gene expression levels from RNA-seq datasets of proliferating, palbo-treated, doxo-treated and INK128-treated SK-Mel-103 and A549 of SASP genes selected for their upregulation in doxorubicin-treated or palbociclib-treated SK-Mel-103 (B) and A549 (C). (D) Flow cytometry analysis of Lysotracker Red in proliferating, TIS (palbo and doxo) and DLCCs SK-Mel-103. Representative histograms showing the fluorescence signal of each stained sample and its unstained control (uncolored histogram). Quantification of the mean fluorescence intensity (MFI) after autofluorescence subtraction (n=3 SK-Mel-103, n=4 A549). ****, P < 0.0001; ***, P < 0.001; **, P < 0.01; *, P < 0.05; one-way ANOVA compared to proliferating control cells. (E) Representative microscopy images of Lysotracker Red staining in proliferating, palbo-treated, doxo-treated and INK128-treated SK-Mel-103 and A549 cells. Scale bar is 100µm. (F) Flow cytometry analysis of Lysotracker Red in proliferating, TIS (palbo and doxo) and DLCCs MCF7, MDA-MB-231 and H1299 cells. Representative histograms showing the fluorescence signal of each stained sample and its unstained control (uncolored histogram). (G) SA-β-GAL staining (blue) of proliferating, palbo-treated, doxo-treated and INK128-treated A549, MCF7, MDA-MB-231, H1299 cells after 7 days from the beginning of the treatment. (H) Flow cytometry analysis of total ROS in proliferating, palbo-treated, doxo-treated and INK128-treated A549 cells. Representative histograms showing the fluorescence signal of each stained sample and its unstained control (uncolored histogram). For SK-Mel-103 and A549, quantification of the MFI after autofluorescence subtraction (n=3). ****, P<0.0001; ***, P < 0.001; **, P < 0.01; one-way ANOVA compared to proliferating control cells. (I) LMNB1 and γ-TUBULIN protein levels in proliferating, palbo-treated, doxo-treated, INK128-treated A549 cells. A representative immunoblot is shown (n=2). *, P < 0,05; one-way ANOVA compared to proliferating controls.

**Figure S3.**
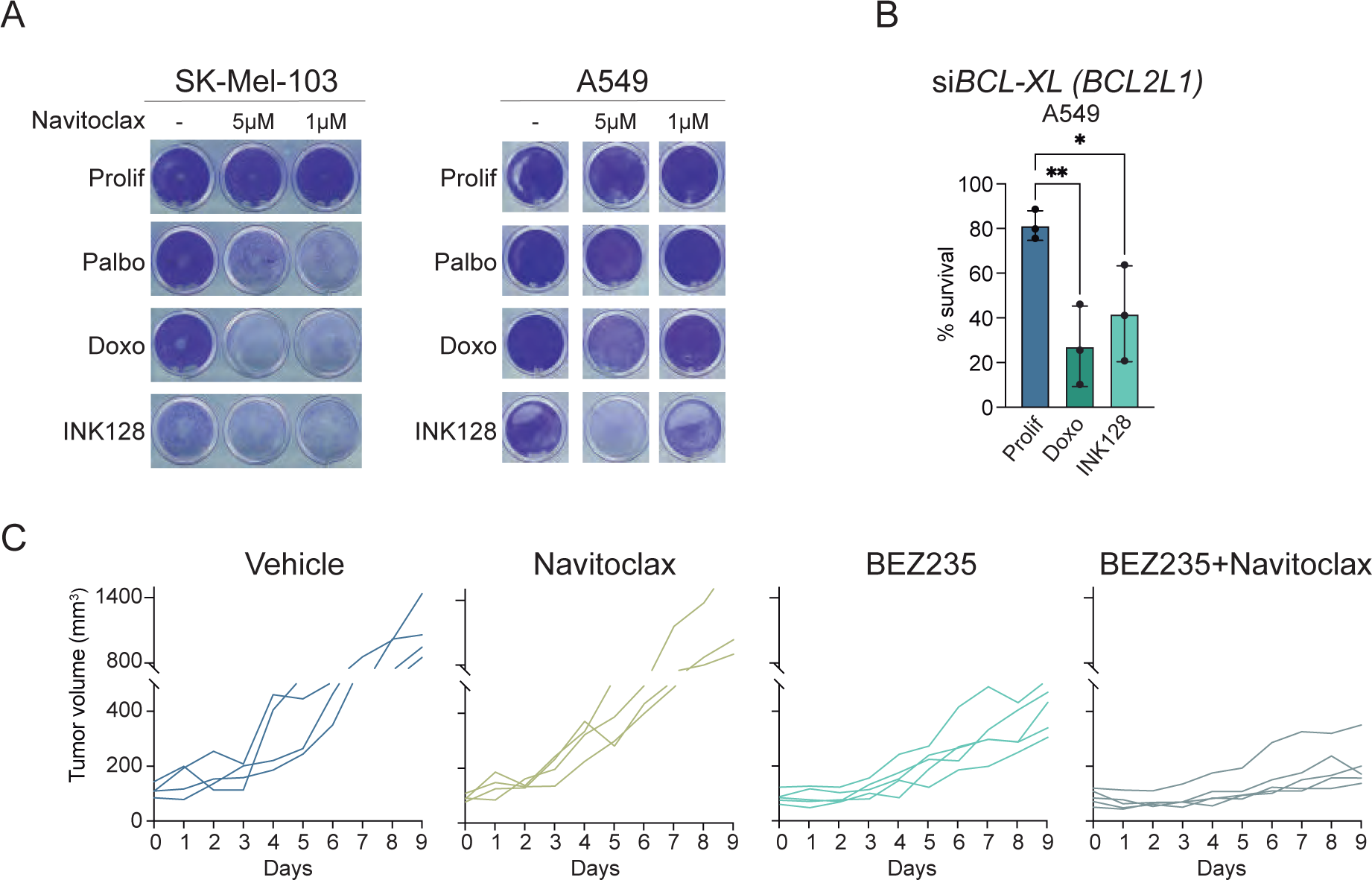
Senescent and persister cancer cells are sensitive to BCL-XL inhibition. (A) Crystal violet assay representative pictures of proliferating, palbo-treated, doxo-treated and INK128-treated SKMel-103 and A549 after Navitoclax treatment (72hrs) at 5μM and 1μM (n=3). (B) CellTiterGlo viability assay of proliferating, doxo-treated and INK128-treated A549 cells treated with siRNAs targeting *BCL-XL* (*BCL2L1*) (n=3). Viability was assessed 5 days after siRNA transfection. Raw data for quantification were acquired by measuring luminescence in a VICTOR Multilabel Plate Reader (Pelkin Elmer). ***, P < 0.001; **, P< 0.01; one-way ANOVA compared to proliferating controls. (C) Individual tumor growth curves from vehicle-treated tumor bearing mice or tumor bearing mice treated with Navitoclax, BEZ235, or BEZ235 + Navitoclax. **** P<0.0001, ***, P < 0.001; **, P < 0.01; two-way ANOVA test (n=4 or 5).

**Figure S4.**
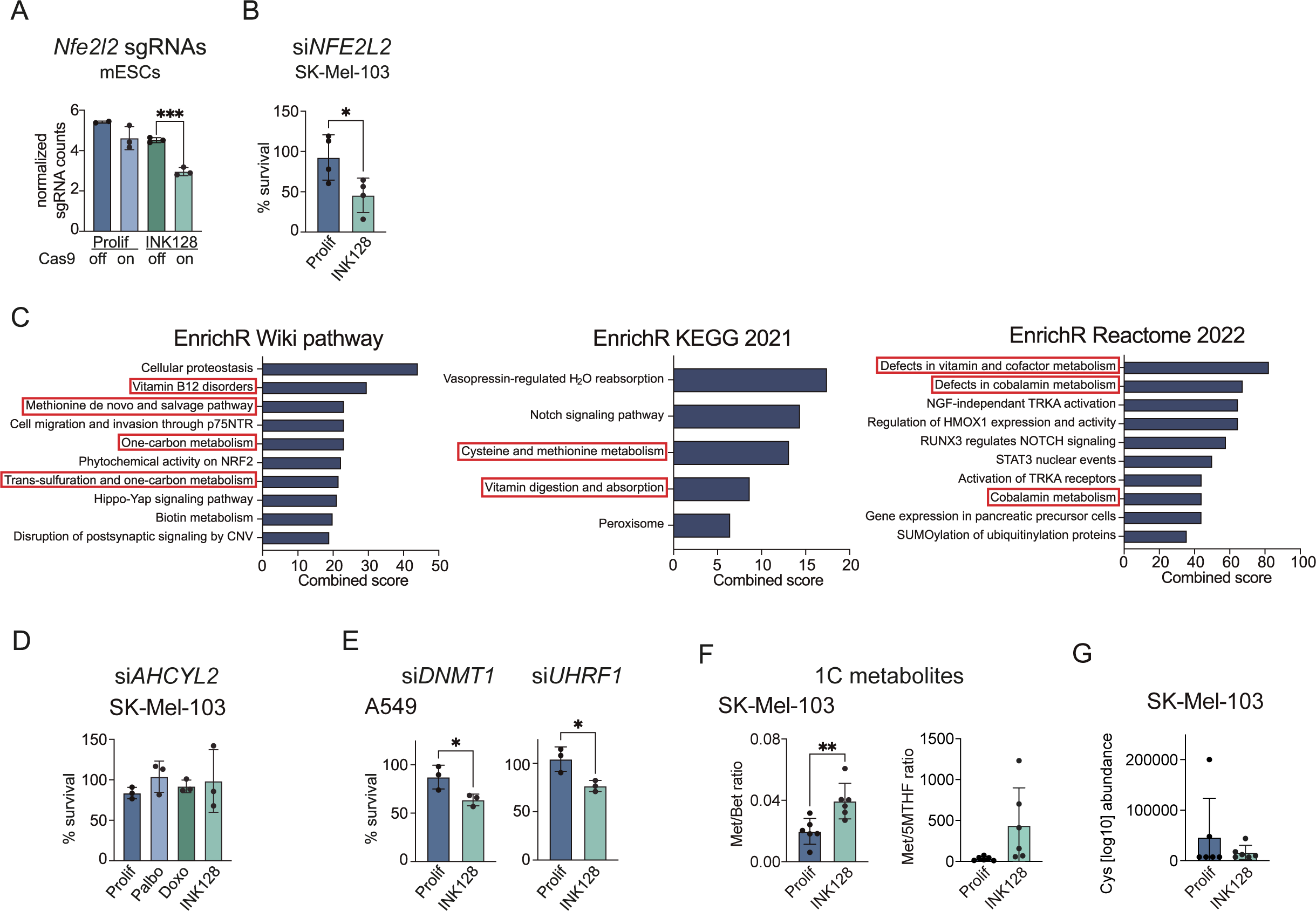
Whole genome CRISPR/Cas9 screening identifies 1C metabolism and DNAme as specific vulnerabilities of persister cells. (A) *Nfe2l2* sgRNA counts in proliferating and diapause mESCs untreated (Cas9 off) and treated (Cas9 on) with doxycycline. ****, P < 0.001, one-way ANOVA versus Cas9 off diapause mESCs (n=2 mESCs prolif Cas9 off, n=3 for all the other conditions). (B) Crystal violet viability assay of proliferating and INK128-treated SK-Mel-103 transfected with si*NRF2* (*NFE2L2*) for 5 days. % of survival was calculated in comparison to the respective siSCR. Raw data for quantification were acquired using the SYNERGY HTX Absorbance microplate reader. *, P < 0.05; unpaired t test compared to proliferating controls (n=4). (C) Top 10 significantly enriched terms from EnrichR Wiki pathway and EnrichR Reactome 2022 when analyzing the genes found depleted in the diapause mESCs treated with doxycycline. Top 5 significantly enriched terms from EnrichR KEGG 2021. Combined score is performed by the EnrichR software by multiplying the log p-value from the Fisher extact test by the z-score of the deviation from the expected rank. (D) Crystal violet viability assay of proliferating, palbo-treated, doxo-treated and INK128-treated SK-Mel-103 transfected with si*AHCYL2*. % of survival was calculated in comparison to the respective siSCR. Raw data for quantification were acquired using the SYNERGY HTX Absorbance microplate reader (n=3). (E) Crystal violet viability assay of proliferating, and INK128-treated A549 transfected with si*DNMT1* and si*UHRF1*. % of survival was calculated in comparison to the respective siSCR. Raw data for quantification were acquired using the SYNERGY HTX Absorbance microplate reader. *, P < 0.05; unpaired t test compared to proliferating controls (n=3). (F) Ratio of Methionine/Betaine (Met/Bet) and Methionine/5-methyltetrahydrofolate (Met/5MTHF) abundance in proliferating and INK128-treated SK-Mel-103 cells. **, P < 0.01; unpaired t test compared to proliferating controls. (G) Total abundance of Cysteine (Cys) in proliferating and INK128-treated SK-Mel-103 cells.

**Figure S5.**
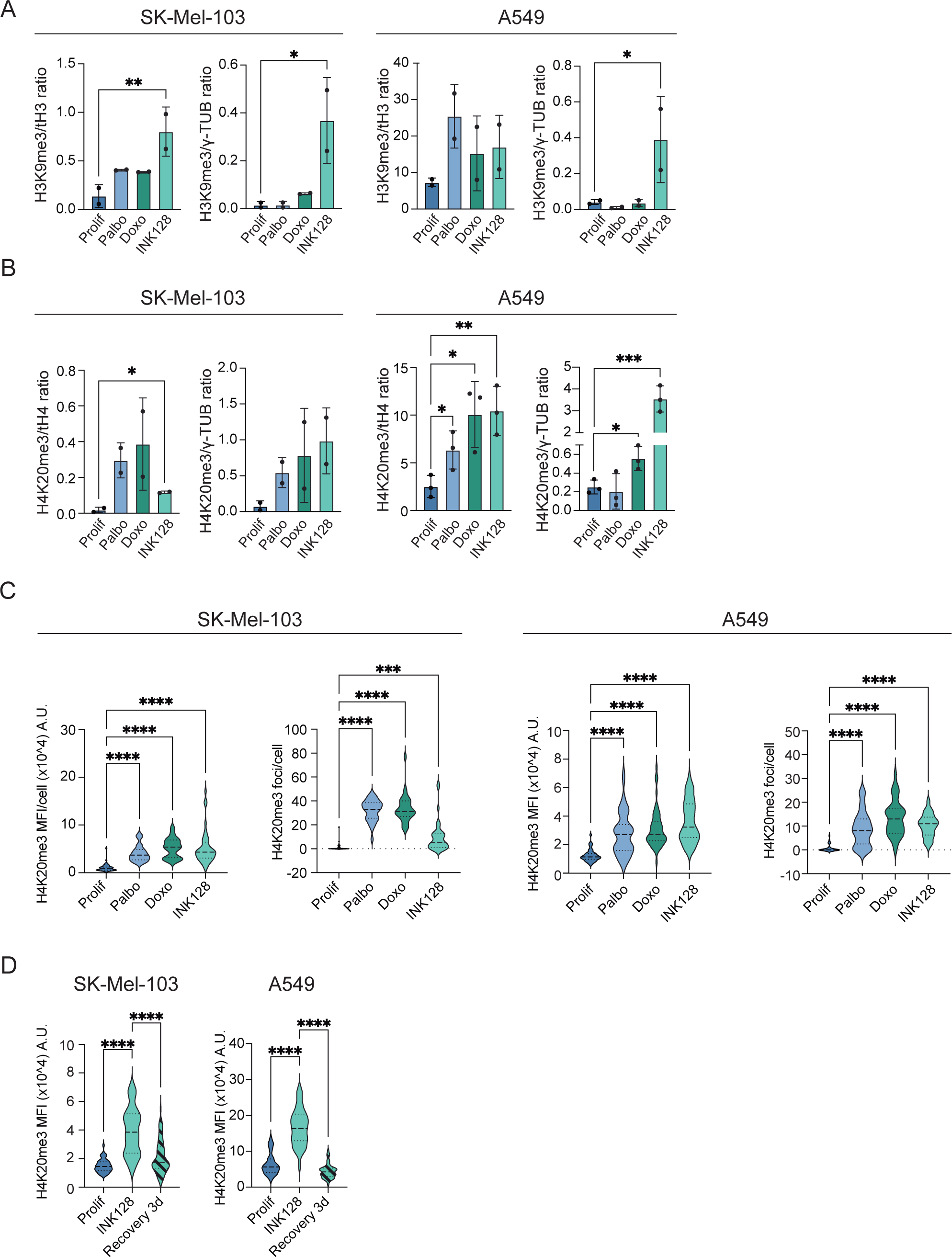
Persister/diapause-like cells present H4K20me3 heterochromatin foci. (A) Immunoblot quantification of H3K9me3 levels normalized on total H3 levels (left) and γ-Tubulin (right) in proliferating, palbo-treated, doxo-treated and INK128-treated SK-Mel-103 and A549 cells. **, P < 0.01; *, P < 0.05; one-way ANOVA compared to proliferating controls (n=2). (B) Immunoblot quantification of H4K20me3 levels normalized on total H4 levels (left) and γ-Tubulin (right) in proliferating, palbo-treated, doxo-treated and INK128-treated SK-Mel-103 and A549 cells. **, P < 0.01; *, P < 0.05; one-way ANOVA compared to proliferating controls (n=2 SK-Mel-103 and n=3 A549). (C) Immunostaining of H4K20me3 (red) and dapi (blue) in proliferating, palbo-treated, doxo-treated and INK128-treated SK-Mel-103 (left) and A549 (right). Quantification of H4K20me3 mean fluorescence intensity (MFI) and foci per nuclei in proliferating, palbo-treated, doxo-treated and INK128-treated SK-Mel-103 and A549. ****, P < 0.0001; one-way ANOVA compared to proliferating control cells. Cells analyzed from 3 independent replicates. Cell numbers were n= 53 (SK-Mel-103 prolif), n= 43 (SK-Mel-103 palbo), n= 38 (SK-Mel-103 doxo), n=44 (SK-Mel-103 INK128); n=46 (A549 prolif), n=51 A549 palbo), n=53 (A549 doxo), n=53 (A549 INK128). (D) Quantification of H4K20me3 mean fluorescence intensity (MFI) in SK-Mel-103 and A549 cells in the following conditions: proliferating, INK128-treated, and cells after 3 days from INK128 withdrawal. ****, P < 0.0001; *, P<0,05; one-way ANOVA. Cell numbers were n=46 (SK-Mel-103 prolif), n=40 (SK-Mel-103 INK128), n=60 (SK-Mel-103 recovery 3d); n=96 (A549 prolif), n=41 A549 INK128), n=77 (A549 recovery 3d).

**Figure S6.**
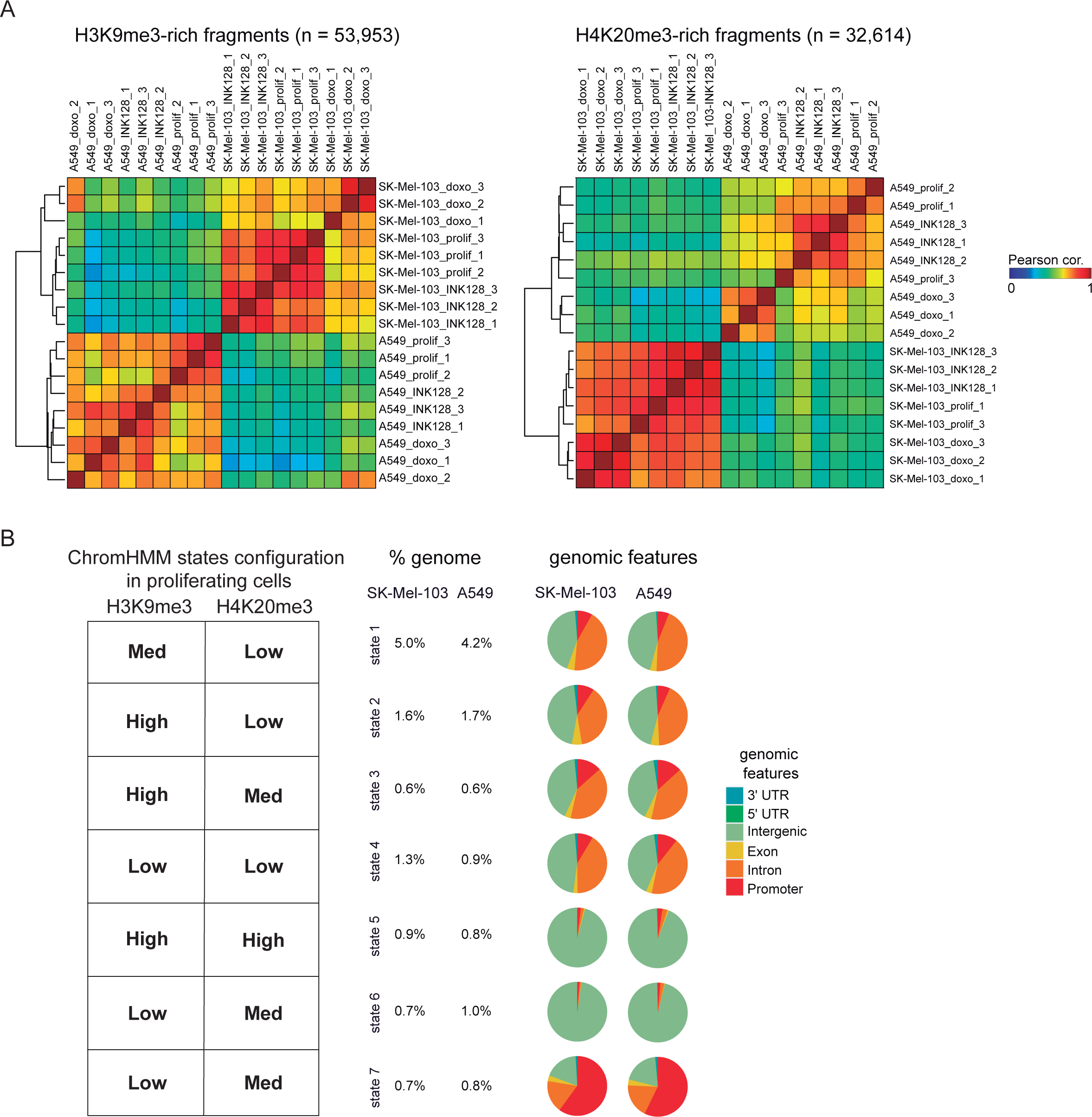
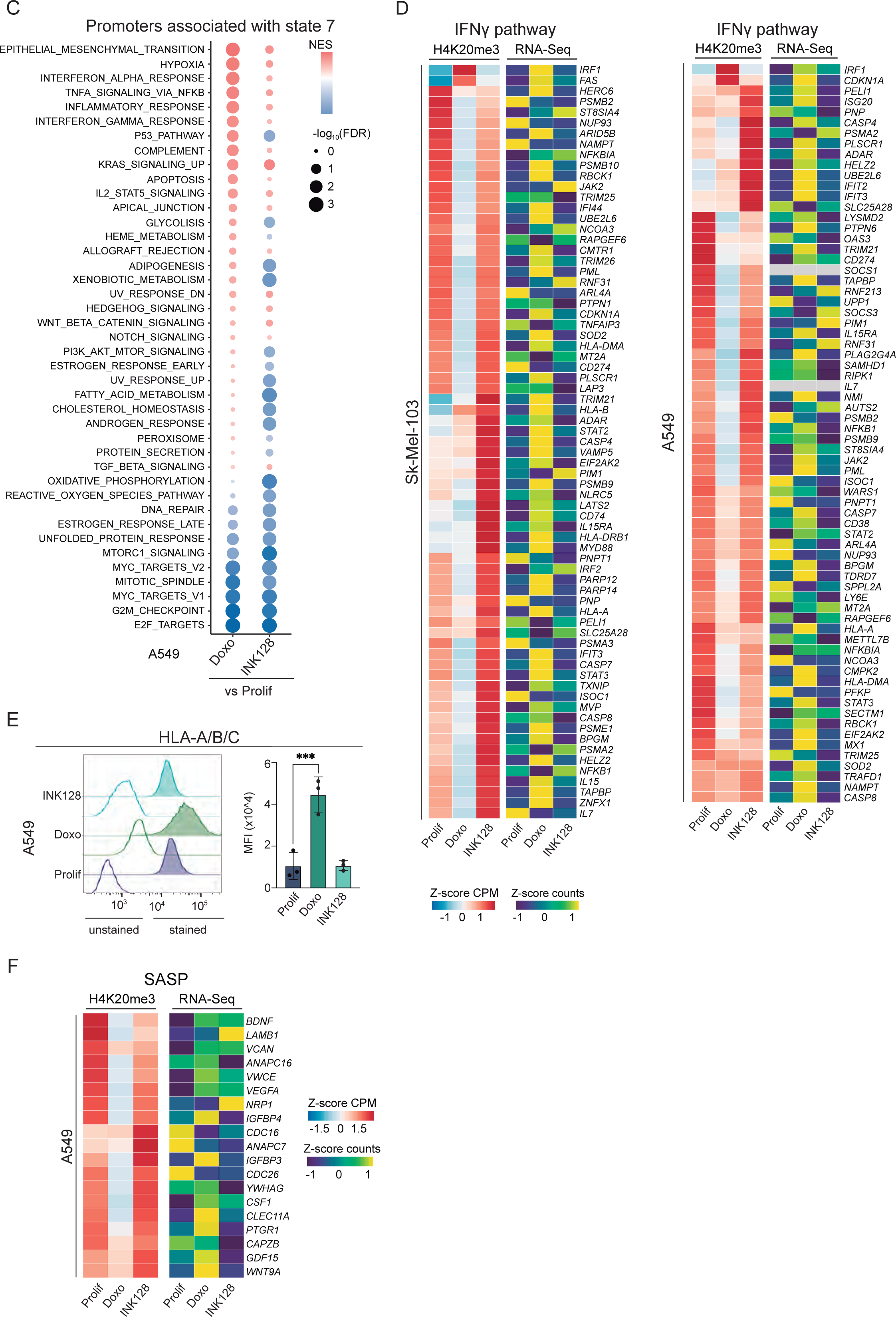
H4K20me3 is differentially associated to IFN and SASP-associated promoter regions in senescent and persister cancer cells. (A) Pair-wise Pearson correlation between replicates based on the enrichment over H3K9me3-rich regions and H4K20me3-rich regions using Deeptools. The regions used in this analysis are the result of intersecting peaks called in control, doxo-treated, and INK128-treated SK-Mel-103 and A549 cells (n=3). (B) Pie charts reflecting the overlapping of ChromHMM states with genomic features in SK-Mel-103 and A549 cells. (C) Dotplots of Pre-ranked GSEA associated to the ChromHMM state 7 of A549 cells, where genes were ranked according to their fold change between treated and controls cells. Normalised Enrichment Score (NES) is colour-coded, while statistical significance (FDR: False Discovery Rate) is encoded in the size of the dots. (D) Heatmaps plotting the levels of H4K20me3 over promoters in gene set ‘Hallmark interferon gamma response’ that intersect with ChromHMM state 7 in proliferating, doxo-treated, INK128-treated SK-Mel-103 and A549 cells. This is accompanied by a heatmap plotting the normalized RNA-Seq counts for these genes, averaging replicates. (E) Flow cytometry analysis of human (HLA-A/B/C) MHC-I expression in proliferating, palbo-treated, doxo-treated and INK128-treated A549 cells. Representative histograms showing the fluorescence signal of each stained sample and its unstained control (uncolored histogram). Quantification of the MFI after autofluorescence subtraction (n=3). ***, P < 0.001; one-way ANOVA compared to proliferating control cells. (F) Heatmaps plotting the levels of H4K20me3 over promoters of SASP genes (previously defined as TIS A549 SASP) that intersect with ChromHMM state 7 in proliferating, doxo-treated and INK128-treated A549 cells. This is accompanied by a heatmap plotting the normalized RNASeq counts for these genes, averaging replicates.

## Experimental procedures

### Cell lines and culture conditions

Human melanoma SKMEL-103, NSCLC A549, H1299, MCF7, MDA-MB-231 cells were obtained from the ATCC. All the cell lines were maintained in standard Dulbecco’s modified Eagle’s medium (DMEM, high glucose, Life Technologies, #12077579) supplemented with 10% heat inactivated fetal bovine serum (FBS) (Gibco, #10270106) and 1% anti-anti (Antibiotic-Antimycotic, Gibco, #15240062).

Mouse embryonic stem cells (mESCs) were provided by Oscar Fernandez-Capetillo (CNIO, Madrid). mESCs were grown on gelatin 1% with ES medium (EM): Dulbecco’s modified Eagle’s medium (DMEM, high glucose, Life Technologies, #12077579) supplemented with 15% foetal bovine serum (FBS) (Gibco, #10270106), 1% anti-anti (Antibiotic-Antimycotic, Gibco, #15240062), LIF (1000 U/ml, Merck Millipore, #ESG1107), 0.1 mM non-essential amino acids (Life Technologies, #11140035), 1% glutamax (Life Technologies, #35050061), and 55 mM β-mercaptoethanol (Merck, #M3148).

All cells were kept in a standard humidified incubator at 37°C and 5% CO_2_. Cells were monthly tested for mycoplasma contamination using standard PCR and only negative cells were used.

### Cell lines treatments

#### Senescence induction

Senescence was induced throughout treatment with the DNA-damaging agent Doxorubicin (100nM in all cell lines except MDA-MB-231 and H1299 50nM) (Sigma, #D1515) and the CDK4/6 inhibitor Palbociclib (1μM in SK-Mel-103, H1299 and MCF6 cells and 5μM in A549 and MDA-MB-231 cells) (PD033299, Absource Diagnostic, #S1116). After 3 days, cells were replenished with complete medium without the drugs. In A549 and MCF7 cells palbociclib was maintained in the media until cell collection. **Diapause-like induction**. Cells were treated with the dual mTOR/PI3K inhibitor INK128 (100nM or 200nM) (Abosurce Diagnostic, #S2811). The inhibitor was maintained in the media until cell collection.

#### Diapause induction

mESCs were treated with INK128 (200nM) for 8 days. The inhibitor was kept in the media until cell collection.

#### Navitoclax treatment

Proliferating, senescent and diapause-like cells were treated with Navitoclax (1μM, 5μM) (Eurodiagnostico, #HY-10087) for 3 days before performing viability assay experiments.

Unless specified, cells were collected 7 days after the beginning of the treatment for performing all the experiments.

### siRNA transfection

All the siRNAs were purchased from siTOOLs. siRNAs have been used at 3nM final concentration. After 7 days from the beginning of the treatment with the senescence and diapause-like inducers, both proliferating, diapause-like and senescent SK-Mel-103 and A549 were transfected with siRNAs for 5 days before performing viability assays. Lipofectamine reagent RNAiMAX (Life Technologies, #13778075) was used at 2μL/mL diluted in Opti-MEM (Life Technologies, #31985062) to perform the transfection.

After 5 days from the transfection, cell viability assays were performed Luminescent assay and Crystal Violet Assay).

### Cell viability assay

Cell viability assay were performed after Navitoclax treatment (72h) or siRNAs transfections (5 days) in control proliferating, senescent and diapause-like SK-Mel-103 and A549 cells using CellTiter-Glo Luminescent Cell Viability Assay (Promega, #G7571) following manufacturer’s instructions or Crystal Violet (Merck, #C6158).

For Cell Titer Glo, raw data were acquired by measuring luminescence in a VICTOR Multilabel Plate Reader (Pelkin Elmer).

For crystal violet staining, cells were first washed once with PBS and then fixed stained through the crystal violet solution (6.179mM crystal violet in 20% methanol solution in H_2_O) for 30 minutes. The solution was then removed from the plates and cells were washed in H_2_O at least 5 times and dry under a chemical hood overnight. Images were taken using an HP scanner. For quantification, the staining was lysed using lysing solution (0.1M sodium citrate, 50% ethanol at pH 4.2). Absorbance was measured using a SYNERGY HTX Absorbance microplate reader (Agilent Technologies) at 570nM.

### Gene expression analysis by RT-qPCR

Total RNA was extracted from cell samples using TRIzol reagent (Invitrogen, #15596018) or the RNeasy Micro Kit (Quiagen). Reverse transcription of up to 1μg of total RNA was performed using the iScript cDNA Synthesis Kit (Bio-Rad, #1725038).

Quantitative real-time PCR (RT-qPCR) was performed using GoTaq PCR Master Mix (Promega, #A6002) and specific primers listed below. The reaction was performed in a QuantStudio 6 Flex thermocycler (Applied Biosystems).

All the reactions were performed in triplicates. *ACTB* and *GAPDH* were used as endogenous normalization controls, and all data were analyzed through the 2^-ΔΔCT^ method (Yuan et al., 2006). The primers used in this study are listed below:

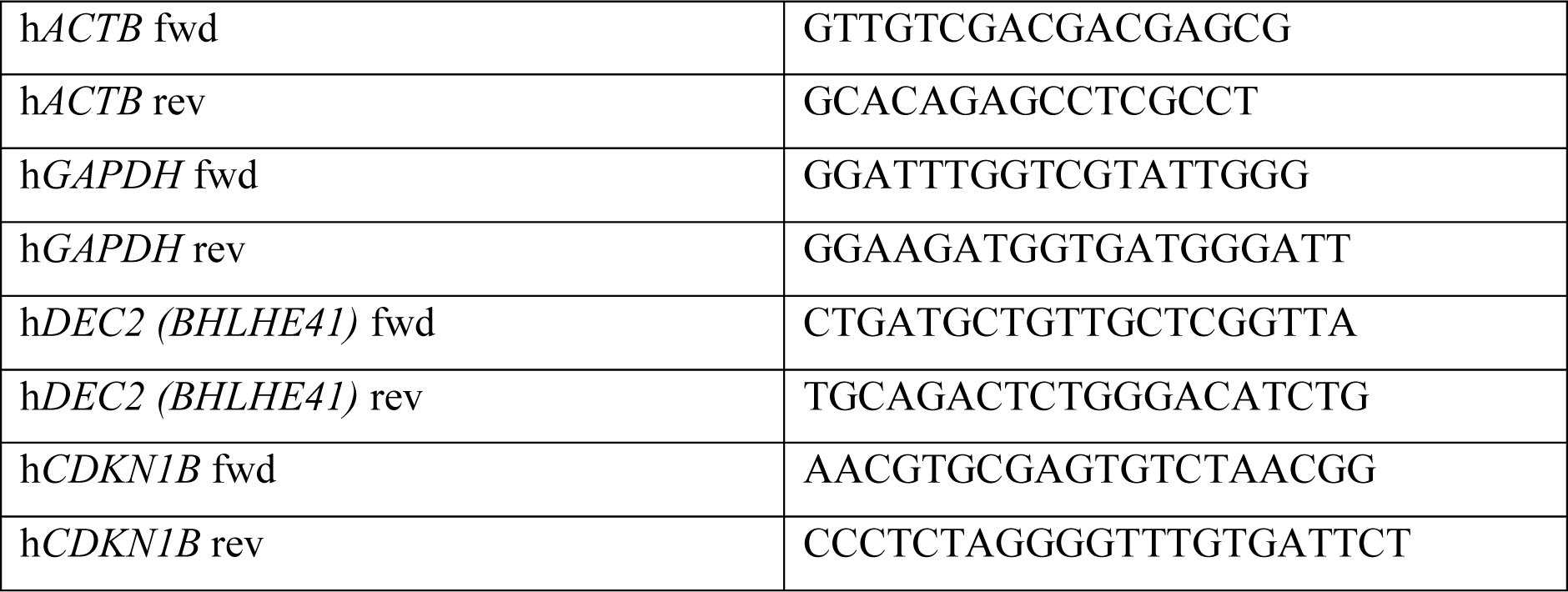

### Immunoblot

Cell pellets were lysed in 50mM Tris-Hcl pH 7.4, 0.5% NP-40, 250mM NaCl, 5mM EDTA, with freshly added 1mM Sodium Orthovanadate, 1mM NaF, protease inhibitors (#87785, Thermofisher) and phosphatase inhibitors (#4906837001, Roche). 15-25μg of proteins were run on NuPage 4-12% gradient Bis-Tris gels in NuPAGE™ MES SDS Running Buffer (20X) and wet-transferred to 0.2μM nitrocellulose membranes (GE Healthcare, #10600001). Blots were blocked in Intercept (TBS) Blocking Buffer (LI-COR, #927-60001) for 30 minutes and then incubated with the following primary antibodies: rabbit Anti-trimethyl-Histone H3 (Lys9) Antibody (1:500, Upstate, #07-442), rabbit Anti-trimethyl-Histone H4 (Lys20) Antibody (1:500, Upstate, #07-749), mouse Anti-Histone H4 Antibody (1:500, Condalab, #61521), mouse anti-Histone H3 (96C10) Antibody (1:500, Cell Signaling, #3638), mouse anti-γ-Tubulin Antibody clone GTU-88 (1:5000, Sigma, #T6557) over-night. The following day the membranes were incubated with the following secondary antibodies: IRDye 800 CW anti-rabbit (1:5000, Licor Odissey, #926-32211) and IRDye 800 CW anti-mouse (1:5000, Licor Odissey, #926-32210).

### Flow cytometry analysis

Cells were digested into single cells by trypsinization (0.25% trypsin-EDTA, Gibco, #25200056). For lysosomal mass measurement, cells were pre-treated with LysoTracker dye (Invitrogen, Thermofisher, #L7528) according to manufacturer’ instructions. Single cells were resuspended in fluorescence-activated cell sorting (FACS) buffer (5mM EDTA and 0.5% BSA in PBS). For apoptosis detection, cells were stained with Annexin-V and Propidium Iodide apoptosis detection kit (Thermofisher, #88-8007-74) according to manufacturer’s instructions. For ROS detection, cells were stained with ROS detection solution from the ROS-ID total ROS detection kit (Enzo Life Sciences, #ENZ-51011) according to manufacturer’s instructions. For blocking, cells were incubated in FACS buffer with 20% FBS for 10 min at 4°C). Cells were stained with HLA-A, B, C antibody (1:100, Biolegend, #311404) for 40 minutes. Cell viability was assessed using propidium iodide (PI, Sigma, #P4864) or Live/Dead Fixable Yellow dye (Invitrogen, #L34967), following the manufacturer’s instructions. Cells were washed once with FACS buffer and run on a Gallios Beckman Coulter flow cytometer (BD Biosciences) or a a FACSAria Fusion (BD Biosciences). The autofluorescence signal from unstained samples was obtained and subtracted from each stained sample in all experiments. Data were analyzed using FlowJo v10 software.

### Cell cycle analysis

Cells were digested into single cells by trypsinization (0.25% trypsin-EDTA, Gibco, #25200056) and resuspended at a concentration of 10^6^ cells/mL in a solution of 30% PBS and 70% ice-cold EtOH for fixation. After at least 2hrs of fixation on ice, cells were resuspended in the following staining solution: 3% PI 500μg/mL in Sodium citrate 38mM (in water), 3% RNAse A 10mg/mL (Life Technologies, #EN0531) in PBS-EDTA 5mM at 4°C. Cells were analyzed by flow cytometry the day after.

### Senescence-associated β-galactosidase staining

After one wash in PBS, cells were fixed in the SA-β-GAL fixing solution (5 nM EGTA, 2 mM MgCl2, 0.2% glutaraldehyde in 0.1 M phosphate buffer) for 15 minutes at room temperature (RT). After washing once with PBS, cells were incubated overnight at 37°C in SA-β-GAL staining solution pH 6 (40 mM citric acid, 5 mM potassium cyanoferrate (II) and (III), 150 mM sodium chloride and 2 mM magnesium chloride, dissolved in 0.1 M phosphate buffer containing 1 mg/mL X-Gal (Melford BioLaboratories, #MB1001) prepared in dimethylformamide (DMF, Sigma, #D4551)). Cells were then washed once with PBS and kept in glycerol solution 0.5%. Pictures were taken using a Nikon Eclipse TS2 bright-field microscope.

### Immunofluorescence

Cells were fixed in 4% paraformaldehyde for 10 minutes, permeabilized in 0.2% Triton X-100 and blocked in 5% FBS and 0.2% Triton X-100. Primary antibodies (rabbit Anti-trimethyl-Histone H3 (Lys9) Antibody (1:500, Upstate, #07-442), rabbit Anti-trimethyl-Histone H4 (Lys20) Antibody (1:500, Upstate, #07-749) were diluted in blocking solution and incubated over night at 4°C. Cells were washed three times with blocking solution for a total of 20 minutes and then incubated with secondary antibody (1:500, Invitrogen, #A31572) for 2h. Cells were washed twice with permeabilization buffer and mounted with Vectashield antifade mounting medium with DAPI (Vector Laboratories, #H1200). For lysotracker staining, cells were incubated with LysoTracker dye (Invitrogen, Thermofisher, #L7528) according to manufacturer’ instructions for 30’, before fixation in 4% PFA.

Confocal images were taken using the super-resolution Airyscan detector of ZEISS Elyra 7 microscope at x100 and x64 magnification or with the Leica SP5 Spectral Confocal Multiphoton at x64 magnification.

### RNA sequencing (RNA-seq)

Total RNA was extracted using Trizol reagent and the Rneasy Mini Kit (Qiagen, #QIA74106), following manufacturer’s insturctions. Total RNA extractions were quantified with a Nanodrop One (Thermo Fisher), and RNA integrity was assessed with the Bioanalyzer 2100 RNA Nano assay (Agilent). Libraries for RNA-seq were prepared at IRB Barcelona Functional Genomics Core Facility. Briefly, mRNA was isolated from 1.2 μg of total RNA and used to generate dual-indexed cDNA libraries with the Illumina stranded mRNA ligation kit (Illumina) and UD Indexes Set A (Illumina). Ten cycles of PCR amplification were applied to all libraries.

Sequencing-ready libraries were quantified using the Qubit dsDNA HS assay (Invitrogen) and quality controlled with the Bioanalyzer 2100 DNA HS assay (Agilent). An equimolar pool was prepared with the thirty-two libraries for SE75 sequencing on a NextSeq550 (Illumina).

Sequencing output was above 1,015 million 75-nt long single-end reads and a minimum of 27.7 million reads were obtained for all samples.

Sequencing-ready libraries were quantified using the Qubit dsDNA HS assay (Invitrogen) and quality controlled with the Bioanalyzer 2100 DNA HS assay (Agilent). An equimolar pool was prepared with the thirty-two libraries for SE75 sequencing on a NextSeq550 (Illumina). Sequencing output was above 1,015 million 75-nt long single-end reads and a minimum of 27.7 million reads were obtained for all samples.

Adapters and low-quality bases (<Q20) were removed from single-end reads with Cutadapt (v4.4)^86^ and TrimGalore (v0.6.10) (https://github.com/FelixKrueger/TrimGalore). Trimmed reads were then mapped to the human reference genome hg38 using the STAR (v2.7.10b) aligner^87^ and the corresponding gene annotation from GENCODE: release 43/ GRCh38.p13. The following parameters were used for mapping of RNA-Seq reads against the reference genome: --outFilterMultimapNmax 1 –outFilterMismatchNmax 999 -- seedSearchStartLmax 50 --outFilterScoreMinOverLread 0.66 --quantMode GeneCounts -- alignEndsType Local. Differential expression analyses were conducted with DESeq2^88^. Lists of genes pre-ranked according to their Log2 fold-change between experimental conditions were submitted to GSEA^89,90^ to test for differential enrichment of GSEA hallmark and in-house SASP gene sets.

Annotations of repetitive elements from the Repeat masker database in the human genome were downloaded from USCS (http://hgdownload.soe.ucsc.edu/goldenPath/mm10/database/) and converted to saf format. The featureCounts function was used to generate a count matrix for the annotated regions. Comparisons between conditions was performed using the DESeq2 package. Repetitive elements were grouped according to their family, as annotated in the original database.

### CRISPR/Cas9 sgRNA screening

To perform the genome-wide screen, we used a CRISPR/Cas9 system previously described^59^. Embryonic stem cells were generated from mice ubiquitously expressing Cas9 under control of the endogenous Cola1a locus and a tetracycline responsive operator transgene; reverse tetracycline-controlled transactivator synthesis is under control of the endogenous ROSA26 locus. ESCs from these mice were infected with a lentiviral library encoding short guide RNAs (sgRNAs) targeting 19,150 mouse genes, with approximately five independent sgRNAs per gene. Addition of doxycycline to the ESCs causes inducible expression of the Cas9 enzyme, which induces editing of the sgRNA targets.

Diapause was induced in mESC through the mTOR/PI3K inhibitor INK128 at 200nM. After 5 days, half of the proliferating cells and half of the diapause cells were sequenced to know the initial diversity of the sgRNAs library. Doxycycline was added to the remaining cells for 3 additional days to induce sgRNA editing before sequencing. Genomic DNA was isolated from cell pellets using a genomic DNA isolation kit (Blood & Cell Culture Midi kit (Qiagen)). After gDNA isolation, sgRNAs were amplified and barcoded by PCR as in Shalem et al., to amplify the DNA fragment containing sgRNA sequences. PCR products were sequenced on a HiSeq 4000 instrument (Illumina) at 50bp reads to a depth of 30M reads per sample.

Reads were preprocessed by removing adapters using Cutadapt v4.1^86^ with parameters “-e 0.2 -a GTTTTAGAGCTAGAAATAGCAAGTTAAAATA-m 18”. Next sequences were trimmed to length 19 to match that of the probes. An artificial genome was created for alignment with Bowtie v0.12.9^91^ using the probes’ sequences and the function bowtie-build with default parameters. Finally, reads were aligned to this genome with parameters -S -t -p 20 -n 1 −l 19. Read counts were imported into R^92^ by reading the sam files and counting the number of occurrences of each probe. The resulting count matrix was normalized using the rlog function from the DESeq2 v1.34.0^88^ R package. A linear model with random and fixed effects was fitted to the normalize probe data for each gene. The condition was used as a fixed covariable while the guides were included as random effects whenever there was more than one. The model was fit with the function lmer from the lme4^88^ package or with the native R lm function if there was only one probe for a given gene. Contrast coefficients and p-values were computed using the glht function from the multcomp^93^ package without any p-value adjustment.

GO gene set collections were downloaded from^94^. Genes quantified in the microarray study were annotated according to the Broad Hallmark^95^. Functional enrichment analyses were performed using a modification of ROAST^96^, a rotation-based approach implemented in the Limma R package, which is especially suitable for small experiments. Such modifications were implemented to accommodate the proposed statistical restandardization^98^ in the ROAST algorithm, which enables its use for competitive testing^99^. The MaxMean^98^ statistic was used for testing gene-set enrichment of the different gene collections. For each gene, the most variable guide within each gene was used in these analyses (median absolute deviation). The results of these analyses were adjusted by multiple comparisons using the Benjamini-Hochberg False Discovery Rate method^100^. All these was performed using the functions of the roastgsa^101^ R package.

### Mass spectrometry analysis

Cells were mixed with 25 μL of tris(2-carboxyethyl)phosphine (TCEP) and 70 μL of 1% formic acid in methanol. Samples were vortexed and left in the freezer for 1 hour, centrifuged for 10 minutes at 12000 rpm and 4°C, and transferred to glass vials for their analysis by LC-MS. LC/MS was performed with a Thermo Scientific Vanquish Horizon UHPLC system interfaced with a Thermo Scientific Orbitrap ID-X Tribrid Mass Spectrometer Waltham, MA).

Metabolites were separated by HILIC chromatography with an InfinityLab Poroshell 120 HILIC-Z (2.1 x 100mm, 2.7µm) column (Agilent technologies). The mobile phase A was 50 mM ammonium acetate in water, and mobile phase B was acetonitrile. Separation was conducted under the following gradient: 0–1 min, isocratic 90% B; 1–6 min decreased to 60% B; 6-6.2 min decreased again to 50% B; 6.2-7 min, isocratic 50% B; 7-7,2 min, increased to 90% B; 7,2–10.5 min, reequilibration column 90% B. The flow was 0.4 mL min-1. The injection volume 5 µL.

Samples were analyzed in targeted SIM mode in positive mode except for THF and Cys which were determined in negative mode. The MS parameters used were: isolation window (m/z), 2; spray voltage, 3500 V; sheath gas, 50; auxiliary gas, 10; Ion transfer tube temperature, 300 °C; vaporizer temperature, 300 °C; Orbitrap resolution, 7500; RF Lens (%), 60; AGC target, 2e5; maximum injection time, 200 ms.

### Cleavage Under Targets and Tagmentation sequencing (CUT&Tag)

CUT&Tag was performed at the Barts Cancer Institute (Queen Mary, University of London), following the published protocol: https://www.protocols.io/view/bench-top-cut-amp-tag-kqdg34qdpl25/v3 The reagents used are listed below:

| Reagents | # |
| --- | --- |
| Beads | Bangs Laboratories BP531-RML |
| Antibody anti-H3K9me3 | Abcam ab176916 |
| Antibody anti-H4K20me3 | Diagenode C15410057 |
| Antibody anti-IgG control | Cell Signaling 3900S |
| Antibody guinea pig anti-rabbit | Antibodies online ABN101961 |
| HEPES | Gibco 15630-056 |
| NaCl | Invitrogen 10255984 |
| Spermidine | S0266 |
| Digitonin | Millipore 300410 |
| EDTA | Millipore 324506-100mL |
| Proteinase K | Thermofisher Scientific EO0491 |
| Potassium Chloride | Sigma-Aldrich P3911 |
| Manganese Chloride | Sigma-Aldrich 203734 |
| Calcium Chloride | Fisher BP510 |
| Potassium Chloride | Sigma-Aldrich P3911 |
| Proteinase Inhibitors tablets | Roche 11873580001 |
| Magnesium Chloride | Invitrogen AM9530G |
| SDS | Sigma Aldrich |
| PCI | Invitrogen 15593049 |
| Chloroform | Sigma 366919-1L |
| Phase-lock tubes | Qiagen 129046 |
| Tris-HCl | AM9855G |
| NEBNext HiFi 2x PCR and Mastermix | NEB M0541S |
| Ampure XP Beads | A63881 |
| Tapestation HSD tape | Agilent 5067-5584 |
| Tapestation buffer | Agilent 5067-5603 |

The indexed primers were described by Buenrostro et al,^102^. The primers used for library preparation are listed below:

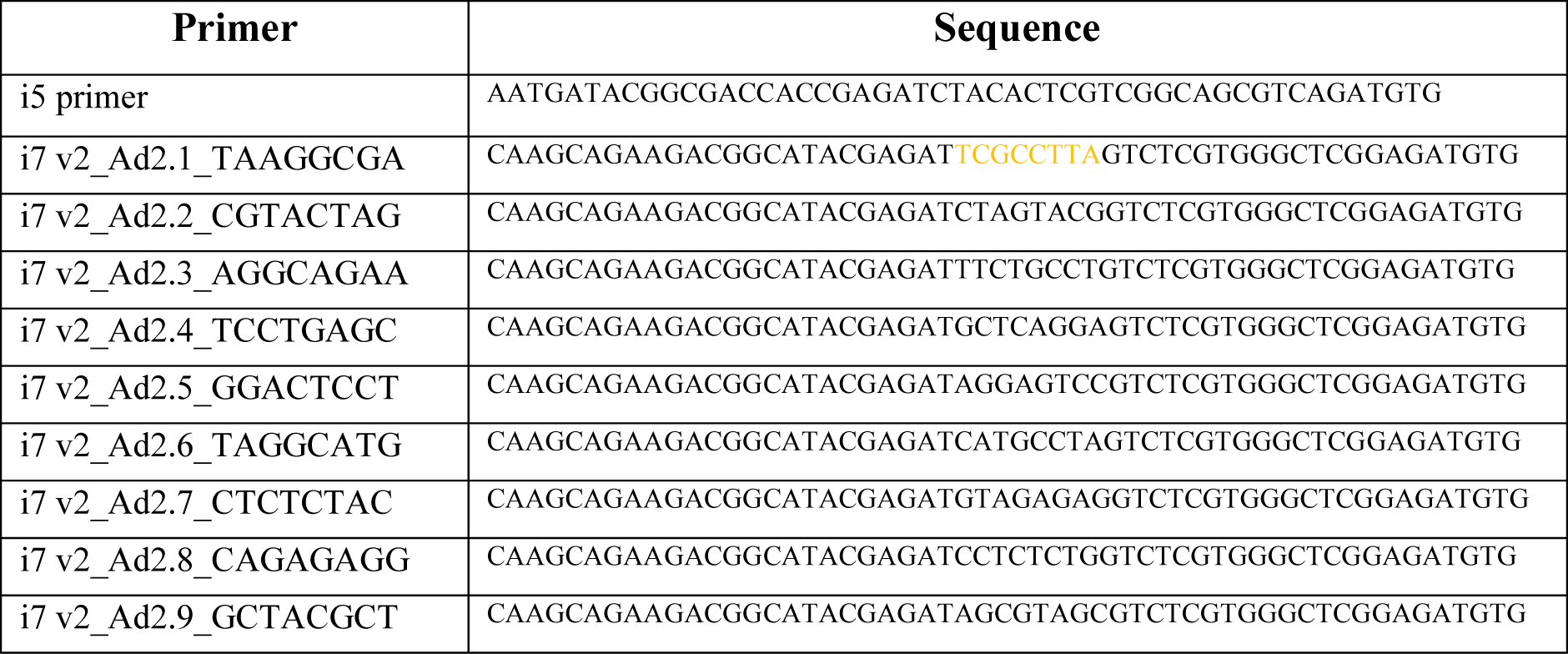
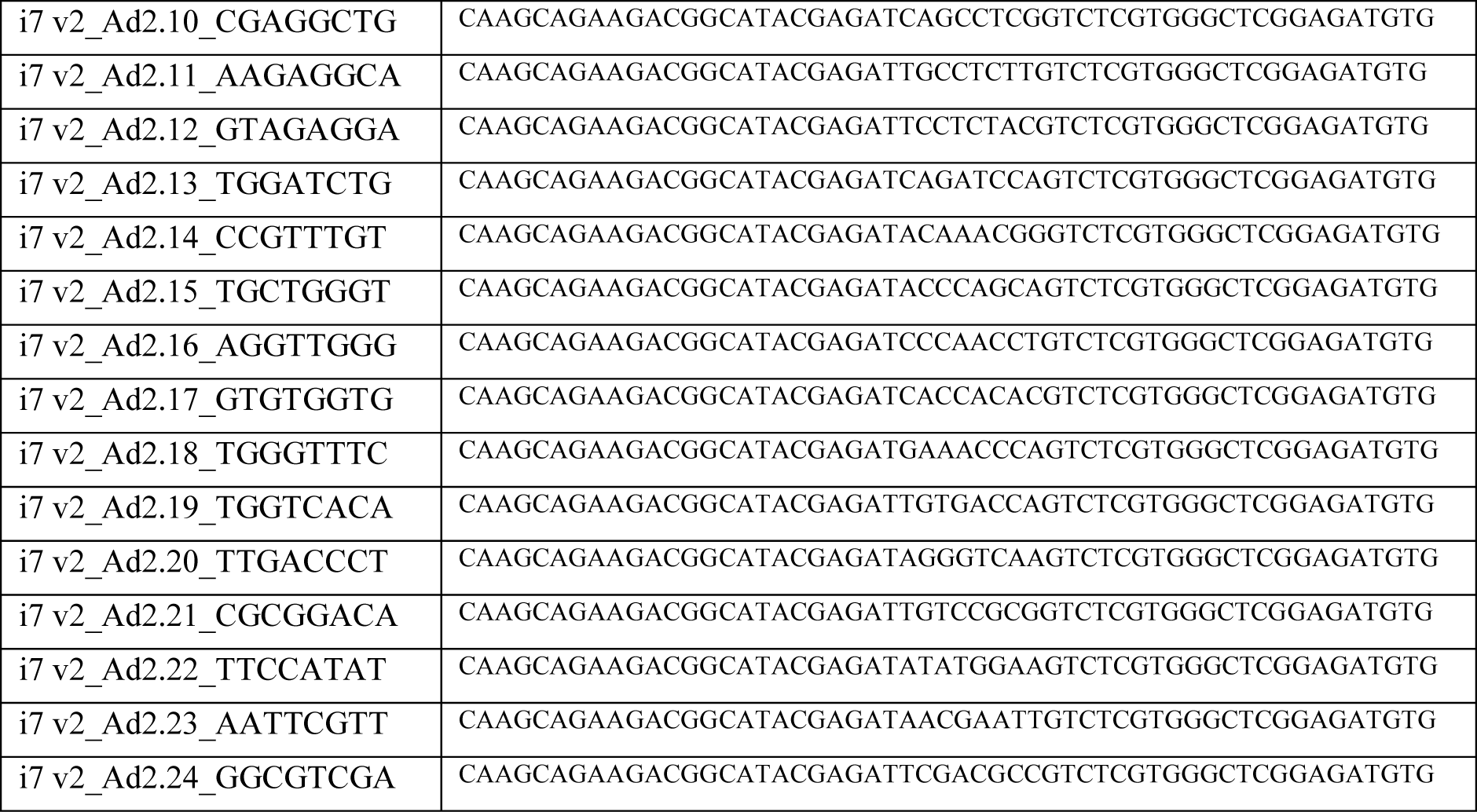

Adapters and low-quality bases (<Q20) were removed from reads with Cutadapt (v4.4)^86^ and TrimGalore (v0.6.10) (https://github.com/FelixKrueger/TrimGalore). Trimmed reads were mapped to the human reference genome hg38 with Bowtie2 (v2.5.1)^103^ using default parameters. PCR duplicates were removed with Picard Tools MarkDuplicates (v1.97). Deduplicated bam files of biological replicates were merged using SAMtools (v1.6) merge^104^. BigWig files were created using DeepTools bamCoverage (v3.0.2)^105^ with the following parameters: --normalizeUsingRPKM –ignoreForNormalization chrX chrY –samFlagInclude 64 –extendReads --binSize 20 --smoothLength 40. Reads overlapping problematic, blacklisted regions (hg38-blacklist.v2.bed)^106^ were excluded from the computation of coverage with the option --blackListFileName. In addition to scaling by library size, in order to remove any composition biases, we calculated a normalizing factor using the BioconductoR csaw::normFactors function^107,108^. This function counts reads in 10-kb genome-wide non-overlapping bins and uses the trimmed mean of M-values (TMM) method to correct for any systematic fold change in the coverage of bins. The normalization factor was passed onto DeepTools bamCoverage with the parameter--scaleFactor.

ChromHMM^71^ was used to identify chromatin fragments with differential enrichment for H3K9me3 and/or H4K20me3 between experimental conditions. The genome was analyzed at 500-bp intervals and a model of 9 states was chosen to characterize both A549 and Sk-Mel-103 cells. 2 ChromHMM states were not included in downstream analyses because they showed no enrichment for neither H3K9me3 or H4K20me3. Genes were assigned to a ChromHMM state according to the percentage of overlap of their promoter regions, defined as TSS +/− 3kb. Enrichment of ChromHMM states for GSEA hallmark gene sets was assessed using a hypergeometric test.

### Pa-Tn5 protein production

To produce double-stranded adaptors with 19mer Tn5 mosaic ends, single stranded DNA oligos were ordered (Eurofins) (sequence information derived from Picelli et al., 2014^109^, listed below:

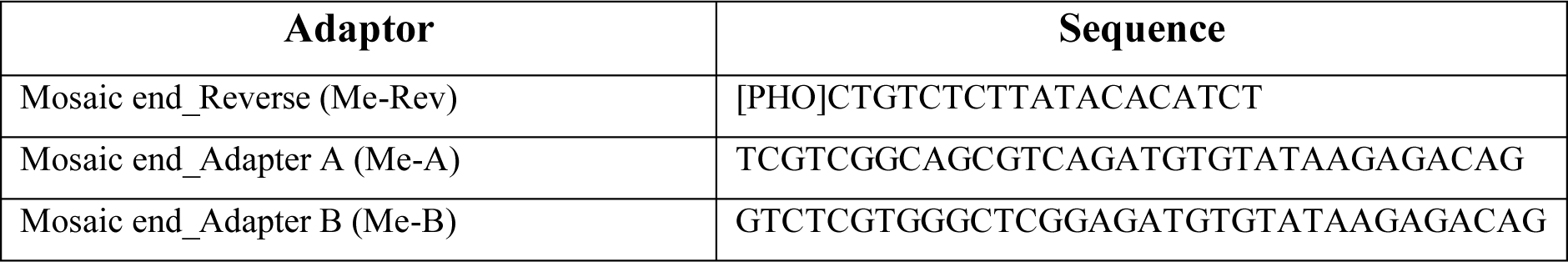

Oligos were resuspended to 200µM in annealing buffer (10mM Tris pH8, 50mM NaCl, 1mM EDTA). To produce double stranded adapters, 20µL Me-Rev and 20µL Me-A were mixed and 20µL Me-Rev and 20µL Me-B were mixed. Tubes were placed in a thermocycler and incubated at 95°C for 2 mins. After the initial incubation, the temperature was reduced by 5°C every 5 minutes until a final 5 mins at 25°C. 3xFLAG-pA-Tn5 purified protein with a concentration of 5.5µM (50% glycerol) was a kind gift from Chema Martin. To assemble the pA-Tn5 adapter transposome, 100µL fusion protein was mixed with 10µL pre-annealed Tn5MEDS-A (200µM) and 10µL Tn5MEDS-B (200µM) oligonucleotides, and incubated at RT for 50 minutes. Produced loaded transposome with a final glycerol concentration of 43%. The transposome was stored at −20°C until *in vitro* enzymatic activity assay or CUT&Tag experiment. pA-Tn5 transposome was incubated with 150ng HMW Lambda DNA (NEB) in digestion buffer (5XTAPS-MgCl2-PEG8000) at 55°C for 7 minutes, prior to incubation with 1% SDS and proteinase K (20mg/mL, NEB) to inactivate and degrade pA-Tn5. Samples were kept on ice, and run on a 0.7% 1X TBE agarose gel. Active pA-Tn5 transposome digested high molecular weight DNA, and produced a smear on the agarose gel. Gel was imaged using UV-fluorescence on Chemidoc (Amersham Imager 600RGB).

### Animal experimentation

Mouse experiments were performed under the Institute for Research in Biomedicine (IRB, Barcelona, Spain) ethical guidelines and regulations. Animals were purchased from Envigo Spain and were housed in strict animal husbandry facilities free of pathogens and micro-organisms. The facilities had controlled temperature environment and at normal 12h day-night light cycles. Mice were randomly selected for the peptide or vehicle treatment and placed in four cages with 4 animals each. Each cage had bedding to provide shelter and privacy as well as unlimited access to food and water. Animals were daily inspected by the qualified personnel and weekly or as needed by the institute’s veterinary doctor.

SK-Mel-103 cells were digested into single cells by trypsinization (0.25% trypsin-EDTA, Invitrogen). Cells were resuspended at a concentration of 10^6^ cells/100μL in a solution of Matrigel (Corning Life Sciences, #354230) and DMEM (1:5) and subcutaneously injected in both flanks of athymic nude mice. When the tumors reached a size of approximately 75 to 100mm^3^, the mice were randomly divided in 6 groups of 4 mice each:

- Untreated
- Palbociclib
- BEZ235
- Navitoclax
- Palbociclib+Navitoclax
- BEZ235+Navitoclax

Palbociclib (100mg/kg in 50mM sodium lactate) (Ambeed, #A295334), BEZ235 (35mg/kg in a solution of PEG400 and PHOSAL 50 PG 1:2) (Medchemtronica, #HY-50673) and Navitoclax (50mg/kg in a solution of PEG400 and PHOSAL 50 PG 1:2) were administered by oral gavage every day for 10 days; for combined treatments, Navitoclax administration started two days after Palbociclib and BEZ235. Xenograft tumors were measured every day using a digital caliper. Tumor volume was calculated using the formula: volume = a x b x b x 0.5, where “a” is the length and “b” is the measured breadth of the tumor lump (skin included). Animals were sacrificed after 10 days from the beginning of the treatment or when tumors reached the size of 1000mm^3^.

### Statistical analysis

Data were analyzed using GraphPad Prism v.9.3.0 software and are represented as mean ± SEM of independent biological replicates. Statistical analyses were performed as described in the figures. Differences were considered significant based on P values (*, P < 0.05; **, P < 0.01; ***, P < 0.001; ****, P < 0.0001).

## Notes

https://www.ncbi.nlm.nih.gov/geo/query/acc.cgi?acc=GSE251660

https://www.ncbi.nlm.nih.gov/geo/query/acc.cgi?acc=GSE246690

https://www.ncbi.nlm.nih.gov/geo/query/acc.cgi?acc=GSE251714

## Bibliography

1. Dhanyamraju, P. K., Schell, T. D., Amin, S. & Robertson, G. P. Drug-Tolerant Persister Cells in Cancer Therapy Resistance. Cancer Research 82, 2503–2514 (2022).

2. Conti, G. D., Dias, M. H. & Bernards, R. Fighting Drug Resistance through the Targeting of Drug-Tolerant Persister Cells. Cancers 13, (2021).

3. Cabanos, H. F. & Hata, A. N. Emerging Insights into Targeted Therapy-Tolerant Persister Cells in Cancer. Cancers 13, 2666 (2021).

4. Wang, B., Kohli, J. & Demaria, M. Senescent Cells in Cancer Therapy: Friends or Foes? Trends in Cancer 6, 838–857 (2020).

5. Vallette, F. M. et al. Dormant, quiescent, tolerant and persister cells: Four synonyms for the same target in cancer. Biochemical Pharmacology 162, 169–176 (2019).

6. Montagner, M. & Sahai, E. In vitro Models of Breast Cancer Metastatic Dormancy. Frontiers in Cell and Developmental Biology 8, (2020).

7. Aguirre-Ghiso, J. A. Translating the Science of Cancer Dormancy to the Clinic. Cancer Research 81, 4673–4675 (2021).

8. Hernandez-Segura, A., Nehme, J. & Demaria, M. Hallmarks of Cellular Senescence. Trends in Cell Biology 28, 436–453 (2018).

9. Gorgoulis, V. et al. Cellular Senescence: Defining a Path Forward. Cell 179, 813–827 (2019).

10. Biran, A. et al. Quantitative identification of senescent cells in aging and disease. Aging Cell 16, 661–671 (2017).

11. Tan, J. X. & Finkel, T. Lysosomes in senescence and aging. EMBO reports 24, e57265 (2023).

12. Dimri, G. P. et al. A biomarker that identifies senescent human cells in culture and in aging skin in vivo. Proceedings of the National Academy of Sciences 92, 9363–9367 (1995).

13. Höhn, A. et al. Happily (n)ever after: Aging in the context of oxidative stress, proteostasis loss and cellular senescence. Redox Biology 11, 482–501 (2017).

14. Sadaie, M. et al. Redistribution of the Lamin B1 genomic binding profile affects rearrangement of heterochromatic domains and SAHF formation during senescence. Genes Dev. 27, 1800–1808 (2013).

15. Narita, M. et al. Rb-Mediated Heterochromatin Formation and Silencing of E2F Target Genes during Cellular Senescence. Cell 113, 703–716 (2003).

16. Shah, P. P. et al. Lamin B1 depletion in senescent cells triggers large-scale changes in gene expression and the chromatin landscape. Genes Dev. 27, 1787–1799 (2013).

17. Birch, J. & Gil, J. Senescence and the SASP: many therapeutic avenues. Genes Dev. 34, 1565–1576 (2020).

18. Faget, D. V., Ren, Q. & Stewart, S. A. Unmasking senescence: context-dependent effects of SASP in cancer. Nat Rev Cancer 19, 439–453 (2019).

19. Chaib, S. et al. The efficacy of chemotherapy is limited by intratumoural senescent cells that persist through the upregulation of PD-L2. 2022.11.04.501681 Preprint at 10.1101/2022.11.04.501681 (2022).

20. Di Micco, R., Krizhanovsky, V., Baker, D. & d’Adda di Fagagna, F. Cellular senescence in ageing: from mechanisms to therapeutic opportunities. Nat Rev Mol Cell Biol 22, 75–95 (2021).

21. Chaib, S., Tchkonia, T. & Kirkland, J. L. Cellular senescence and senolytics: the path to the clinic. Nat Med 28, 1556–1568 (2022).

22. Wang, L., Lankhorst, L. & Bernards, R. Exploiting senescence for the treatment of cancer. Nat Rev Cancer 22, 340–355 (2022).

23. Sharma, S. V. et al. A Chromatin-Mediated Reversible Drug-Tolerant State in Cancer Cell Subpopulations. Cell 141, 69–80 (2010).

24. Álvarez-Varela, A. et al. Mex3a marks drug-tolerant persister colorectal cancer cells that mediate relapse after chemotherapy. Nat Cancer 3, 1052–1070 (2022).

25. Guler, G. D. et al. Repression of Stress-Induced LINE-1 Expression Protects Cancer Cell Subpopulations from Lethal Drug Exposure. Cancer Cell 32, 221–237.e13 (2017).

26. Rehman, S. K. et al. Colorectal Cancer Cells Enter a Diapause-like DTP State to Survive Chemotherapy. Cell 184, 226–242.e21 (2021).

27. Dhimolea, E. et al. An Embryonic Diapause-like Adaptation with Suppressed Myc Activity Enables Tumor Treatment Persistence. Cancer Cell 39, 240–256.e11 (2021).

28. Hangauer, M. J. et al. Drug-tolerant persister cancer cells are vulnerable to GPX4 inhibition. Nature 551, 247–250 (2017).

29. Li, Y. et al. Mitophagy is a novel protective mechanism for drug-tolerant persister (DTP) cancer cells. Autophagy 0, 1–2 (2023).

30. Zhang, Z., Tan, Y., Huang, C. & Wei, X. Redox signaling in drug-tolerant persister cells as an emerging therapeutic target. eBioMedicine 89, (2023).

31. Saudemont, A. & Quesnel, B. In a model of tumor dormancy, long-term persistent leukemic cells have increased B7-H1 and B7.1 expression and resist CTL-mediated lysis. Blood 104, 2124–2133 (2004).

32. Pommier, A. et al. Unresolved endoplasmic reticulum stress engenders immune-resistant, latent pancreatic cancer metastases. Science 360, eaao4908 (2018).

33. Romaniello, D. et al. Senescence-associated reprogramming induced by interleukin-1 impairs response to EGFR neutralization. Cellular & Molecular Biology Letters 27, 20 (2022).

34. Wang, M. et al. EGF Receptor Inhibition Radiosensitizes NSCLC Cells by Inducing Senescence in Cells Sustaining DNA Double-Strand Breaks. Cancer Research 71, 6261–6269 (2011).

35. Rudolf, E., John, S. & Cervinka, M. Irinotecan induces senescence and apoptosis in colonic cells in vitro. Toxicology Letters 214, 1–8 (2012).

36. Haug, K., Kravik, K. L. & Angelis, P. M. D. Cellular Response to Irinotecan in Colon Cancer Cell Lines Showing Differential Response to 5-Fluorouracil. ANTICANCER RESEARCH (2008).

37. Willobee, B. A. et al. Combined Blockade of MEK and CDK4/6 Pathways Induces Senescence to Improve Survival in Pancreatic Ductal Adenocarcinoma. Molecular Cancer Therapeutics 20, 1246–1256 (2021).

38. Schwarze, S. R., Fu, V. X., Desotelle, J. A., Kenowski, M. L. & Jarrard, D. F. The Identification of Senescence-Specific Genes during the Induction of Senescence in Prostate Cancer Cells. Neoplasia 7, 816–823 (2005).

39. Criscione, S. W. et al. The landscape of therapeutic vulnerabilities in EGFR inhibitor osimertinib drug tolerant persister cells. npj Precis. Onc. 6, 1–13 (2022).

40. Tetsu, O., Phuchareon, J., Eisele, D. W., Hangauer, M. J. & McCormick, F. AKT inactivation causes persistent drug tolerance to EGFR inhibitors. Pharmacological Research 102, 132–137 (2015).

41. Kurppa, K. J. et al. Treatment-Induced Tumor Dormancy through YAP-Mediated Transcriptional Reprogramming of the Apoptotic Pathway. Cancer Cell 37, 104–122.e12 (2020).

42. Shen, S., et al. Melanoma Persister Cells Are Tolerant to BRAF/MEK Inhibitors via ACOX1-Mediated Fatty Acid Oxidation. Cell Reports 33, (2020).

43. Bulut-Karslioglu, A. et al. Inhibition of mTOR induces a paused pluripotent state. Nature 540, 119–123 (2016).

44. Iyer, D. P. et al. Delay of human early development via in vitro diapause. 2023.05.29.541316 Preprint at 10.1101/2023.05.29.541316 (2023).

45. Yang, X. et al. Inhibition of DEC2 is necessary for exiting cell dormancy in salivary adenoid cystic carcinoma. Journal of Experimental & Clinical Cancer Research 40, 169 (2021).

46. Bragado, P. et al. TGF-β2 dictates disseminated tumour cell fate in target organs through TGF-β-RIII and p38α/β signalling. Nat Cell Biol 15, 1351–1361 (2013).

47. Scognamiglio, R. et al. Myc Depletion Induces a Pluripotent Dormant State Mimicking Diapause. Cell 164, 668–680 (2016).

48. Brown, J. A. et al. TGF-β-Induced Quiescence Mediates Chemoresistance of Tumor-Propagating Cells in Squamous Cell Carcinoma. Cell Stem Cell 21, 650–664.e8 (2017).

49. Ren, D. et al. Wnt5a induces and maintains prostate cancer cells dormancy in bone. Journal of Experimental Medicine 216, 428–449 (2018).

50. Basisty, N. et al. A proteomic atlas of senescence-associated secretomes for aging biomarker development. PLOS Biology 18, e3000599 (2020).

51. Kurz, D. J., Decary, S., Hong, Y. & Erusalimsky, J. D. Senescence-associated β-galactosidase reflects an increase in lysosomal mass during replicative ageing of human endothelial cells. Journal of Cell Science 113, 3613–3622 (2000).

52. Lee, B. Y. et al. Senescence-associated β-galactosidase is lysosomal β-galactosidase. Aging Cell 5, 187–195 (2006).

53. Zhu, Y. et al. Identification of a novel senolytic agent, navitoclax, targeting the Bcl-2 family of anti-apoptotic factors. Aging Cell 15, 428–435 (2016).

54. Zhu, Y. et al. The Achilles’ heel of senescent cells: from transcriptome to senolytic drugs. Aging Cell 14, 644–658 (2015).

55. Zhu, Y. et al. New agents that target senescent cells: the flavone, fisetin, and the BCL-X_L_ inhibitors, A1331852 and A1155463. Aging 9, 955–963 (2017).

56. Mas-Bargues, C., Borrás, C. & Viña, J. Bcl-xL as a Modulator of Senescence and Aging. International Journal of Molecular Sciences 22, 1527 (2021).

57. Bharti, V. et al. BCL-xL inhibition potentiates cancer therapies by redirecting the outcome of p53 activation from senescence to apoptosis. Cell Reports 41, 111826 (2022).

58. Rahman, M. et al. Selective Vulnerability of Senescent Glioblastoma Cells to BCL-XL Inhibition. Molecular Cancer Research 20, 938–948 (2022).

59. Ruiz, S. et al. A Genome-wide CRISPR Screen Identifies CDC25A as a Determinant of Sensitivity to ATR Inhibitors. Molecular Cell 62, 307–313 (2016).

60. Fox, D. B. et al. NRF2 activation promotes the recurrence of dormant tumour cells through regulation of redox and nucleotide metabolism. Nat Metab 2, 318–334 (2020).

61. Ducker, G. S. & Rabinowitz, J. D. One-Carbon Metabolism in Health and Disease. Cell Metabolism 25, 27–42 (2017).

62. Kerr, S. J. Competing Methyltransferase Systems. Journal of Biological Chemistry 247, 4248–4252 (1972).

63. Barroso, M., Handy, D. E. & Castro, R. The Link Between Hyperhomocysteinemia and Hypomethylation: Implications for Cardiovascular Disease. Journal of Inborn Errors of Metabolism and Screening 5, 2326409817698994 (2017).

64. Ren, W. et al. DNMT1 reads heterochromatic H4K20me3 to reinforce LINE-1 DNA methylation. Nat Commun 12, 2490 (2021).

65. Dnmt1 has de novo activity targeted to transposable elements | Nature Structural & Molecular Biology. https://www.nature.com/articles/s41594-021-00603-8.

66. Direct readout of heterochromatic H3K9me3 regulates DNMT1-mediated maintenance DNA methylation | PNAS. https://www.pnas.org/doi/10.1073/pnas.2009316117.

67. Mancini, M., Magnani, E., Macchi, F. & Bonapace, I. M. The multi-functionality of UHRF1: epigenome maintenance and preservation of genome integrity. Nucleic Acids Res 49, 6053–6068 (2021).

68. Boonsanay, V. et al. Regulation of Skeletal Muscle Stem Cell Quiescence by Suv4-20h1-Dependent Facultative Heterochromatin Formation. Cell Stem Cell 18, 229–242 (2016).

69. Evertts, A. G. et al. H4K20 methylation regulates quiescence and chromatin compaction. Mol Biol Cell 24, 3025–3037 (2013).

70. Nelson, D. M. et al. Mapping H4K20me3 onto the chromatin landscape of senescent cells indicates a function in control of cell senescence and tumor suppression through preservation of genetic and epigenetic stability. Genome Biology 17, 158 (2016).

71. Ernst, J. & Kellis, M. ChromHMM: automating chromatin-state discovery and characterization. Nat Methods 9, 215–216 (2012).

72. Marin, I. et al. Cellular Senescence Is Immunogenic and Promotes Antitumor Immunity. Cancer Discovery 13, 410–431 (2023).

73. Chen, H.-A. et al. Senescence Rewires Microenvironment Sensing to Facilitate Antitumor Immunity. Cancer Discovery 13, 432–453 (2023).

74. Aguirre-Ghiso, J. A. & Sosa, M. S. Emerging Topics on Disseminated Cancer Cell Dormancy and the Paradigm of Metastasis. Annual Review of Cancer Biology 2, 377–393 (2018).

75. Cellular Dormancy in Cancer: Mechanisms and Potential Targeting Strategies. https://www.e-crt.org/journal/view.php?doi=10.4143/crt.2023.468.

76. Laberge, R.-M. et al. MTOR regulates the pro-tumorigenic senescence-associated secretory phenotype by promoting IL1A translation. Nat Cell Biol 17, 1049–1061 (2015).

77. Herranz, N. et al. mTOR regulates MAPKAPK2 translation to control the senescence-associated secretory phenotype. Nat Cell Biol 17, 1205–1217 (2015).

78. Kim, Y.-J. et al. Human prx1 Gene Is a Target of Nrf2 and Is Up-regulated by Hypoxia/Reoxygenation: Implication to Tumor Biology. Cancer Research 67, 546–554 (2007).

79. Tang, C. et al. Peroxiredoxin-1 ameliorates pressure overload-induced cardiac hypertrophy and fibrosis. Biomedicine & Pharmacotherapy 129, 110357 (2020).

80. Bostick, M. et al. UHRF1 Plays a Role in Maintaining DNA Methylation in Mammalian Cells. Science 317, 1760–1764 (2007).

81. Sharif, J. et al. The SRA protein Np95 mediates epigenetic inheritance by recruiting Dnmt1 to methylated DNA. Nature 450, 908–912 (2007).

82. Balint, B., Jepchumba, V. K., Guéant, J.-L. & Guéant-Rodriguez, R.-M. Mechanisms of homocysteine-induced damage to the endothelial, medial and adventitial layers of the arterial wall. Biochimie 173, 100–106 (2020).

83. Chandra, T. et al. Independence of Repressive Histone Marks and Chromatin Compaction during Senescent Heterochromatic Layer Formation. Molecular Cell 47, 203–214 (2012).

84. Xu, J. & Kidder, B. L. H4K20me3 co-localizes with activating histone modifications at transcriptionally dynamic regions in embryonic stem cells. BMC Genomics 19, 514 (2018).

85. Abbas, T. et al. CRL4Cdt2 Regulates Cell Proliferation and Histone Gene Expression by Targeting PR-Set7/Set8 for Degradation. Molecular Cell 40, 9–21 (2010).

86. Martin, M. Cutadapt removes adapter sequences from high-throughput sequencing reads. EMBnet.journal 17, 10–12 (2011).

87. Dobin, A. et al. STAR: ultrafast universal RNA-seq aligner. Bioinformatics 29, 15–21 (2013).

88. Love, M. I., Huber, W. & Anders, S. Moderated estimation of fold change and dispersion for RNA-seq data with DESeq2. Genome Biology 15, 550 (2014).

89. Mootha, V. K. et al. PGC-1α-responsive genes involved in oxidative phosphorylation are coordinately downregulated in human diabetes. Nat Genet 34, 267–273 (2003).

90. Subramanian, A. et al. Gene set enrichment analysis: A knowledge-based approach for interpreting genome-wide expression profiles. Proceedings of the National Academy of Sciences 102, 15545–15550 (2005).

91. Langmead, B., Trapnell, C., Pop, M. & Salzberg, S. L. Ultrafast and memory-efficient alignment of short DNA sequences to the human genome. Genome Biology 10, R25 (2009).

92. R: A Language and Environment for Statistical Computing | BibSonomy. https://www.bibsonomy.org/bibtex/7469ffee3b07f9167cf47e7555041ee7.

93. Hothorn, T., Bretz, F. & Westfall, P. Simultaneous Inference in General Parametric Models. Biometrical Journal 50, 346–363 (2008).

94. The Gene Ontology Consortium et al. The Gene Ontology knowledgebase in 2023. Genetics 224, iyad031 (2023).

95. Liberzon, A. et al. The Molecular Signatures Database (MSigDB) hallmark gene set collection. Cell Syst 1, 417–425 (2015).

96. Wu, D. et al. ROAST: rotation gene set tests for complex microarray experiments. Bioinformatics 26, 2176–2182 (2010).

97. Ritchie, M. E. et al. limma powers differential expression analyses for RNA-sequencing and microarray studies. Nucleic Acids Research 43, e47 (2015).

98. Efron, B. & Tibshirani, R. On testing the significance of sets of genes. Ann. Appl. Stat. 1, (2007).

99. Goeman, J. J. & Bühlmann, P. Analyzing gene expression data in terms of gene sets: methodological issues. Bioinformatics 23, 980–987 (2007).

100. Benjamini, Y. & Hochberg, Y. Controlling the False Discovery Rate: A Practical and Powerful Approach to Multiple Testing. Journal of the Royal Statistical Society: Series B (Methodological) 57, 289–300 (1995).

101. Mestres, A. C., Llergo, A. B. & Attolini, C. S.-O. A comparison of rotation-based scores for gene set analysis. 2021.03.23.436604 Preprint at 10.1101/2021.03.23.436604 (2021).

102. Buenrostro, J. D. et al. Single-cell chromatin accessibility reveals principles of regulatory variation. Nature 523, 486–490 (2015).

103. Langmead, B. & Salzberg, S. L. Fast gapped-read alignment with Bowtie 2. Nat Methods 9, 357–359 (2012).

104. Danecek, P. et al. Twelve years of SAMtools and BCFtools. Gigascience 10, giab008 (2021).

105. Ramírez, F., Dündar, F., Diehl, S., Grüning, B. A. & Manke, T. deepTools: a flexible platform for exploring deep-sequencing data. Nucleic Acids Research 42, W187–W191 (2014).

106. Amemiya, H. M., Kundaje, A. & Boyle, A. P. The ENCODE Blacklist: Identification of Problematic Regions of the Genome. Sci Rep 9, 9354 (2019).

107. Lun, A. T. L. & Smyth, G. K. De novo detection of differentially bound regions for ChIP-seq data using peaks and windows: controlling error rates correctly. Nucleic Acids Research 42, e95 (2014).

108. Lun, A. T. L. & Smyth, G. K. csaw: a Bioconductor package for differential binding analysis of ChIP-seq data using sliding windows. Nucleic Acids Research 44, e45 (2016).

109. Picelli, S. et al. Tn5 transposase and tagmentation procedures for massively scaled sequencing projects. Genome Res. 24, 2033–2040 (2014).

