## Supplementary tables for "Persister cancer cells are characterized by H4K20me3 heterochromatin that defines a low inflammatory profile"

| Genes | SK-Mel-103 INK128/Prolif FC | A549 INK128/Prolif FC |
|---|---|---|
| *GSTM3* | -11,896652 | -1,6924377 |
| *F2RL2* | -6,9331557 | -2,5886472 |
| *CCDC160* | -6,5587293 | -8,8609957 |
| *SCDP1* | -5,7792466 | -4,6193715 |
| *TM4SF18* | -5,5612484 | -2,4436962 |
| *PRRG4* | -4,9419815 | -2,7264564 |
| *SCD* | -4,9313462 | -3,9751441 |
| *ANKRD1* | -4,9110621 | -4,6069915 |
| *BCAT1* | -4,9105612 | -4,9231524 |
| *SRGN* | -4,3921115 | -1,962996 |
| *OSBPL6* | -4,1133789 | -1,6864024 |
| *STYK1* | -3,8850434 | -3,7384768 |
| *CHAC1* | -3,5467054 | -5,52457 |
| *UCHL3* | -3,5417703 | -2,1522389 |
| *ENPP1* | -3,3828802 | -2,3817351 |
| *AOX1* | -3,1338236 | -1,8270435 |
| *PGAM4* | -3,0834325 | 1,67222648 |
| *PGAM1* | -3,0197616 | -2,8446207 |
| *SAMHD1* | -2,9758001 | -1,8953392 |
| *DNAJA1* | -2,9614723 | -2,0375992 |
| *PGAM1P2* | -2,9069351 | -2,9776996 |
| *ITGAE* | -2,892908 | -2,905691 |
| *FAM98A* | -2,8579639 | -2,1529735 |
| *TMED5* | -2,713668 | -1,6593968 |
| *PRDX1* | -2,6824639 | -1,9996379 |
| *CLU* | -2,668422 | -3,2195025 |
| *ADI1* | -2,6678704 | -2,5681357 |
| *ADGRE1* | -2,6484342 | -1,6145656 |
| *EIF2S2P4* | -2,6176393 | -2,3959445 |
| *PCCB* | -2,5742218 | -1,6516901 |
| *AKR1C1* | -2,5316441 | -1,9574642 |
| *TIMM17A* | -2,4720405 | -2,3575826 |
| *CNN3* | -2,4167958 | -1,9287264 |
| *GPT2* | -2,4023652 | -2,0066864 |
| *FKBP14* | -2,361605 | -2,8612558 |
| *RBM24* | -2,3612968 | -2,6444713 |
| *HSPA4* | -2,3497953 | -2,0467277 |
| *HIGD1AP1* | -2,3329953 | -2,1290639 |
| *VDAC1P4* | -2,3288108 | -2,4753858 |
| *RDH11* | -2,3188372 | -1,7436929 |

| | | |
|---|---|---|
| *SLC31A1* | -2,3169027 | -2,2592305 |
| *ME1* | -2,3156445 | -1,7987431 |
| *AIFM1* | -2,273268 | -2,0076601 |
| *SLC25A30* | -2,2646275 | -2,0550995 |
| *LDLRAP1* | -2,2348323 | -1,5773321 |
| *TPI1P2* | -2,2217983 | -2,0401086 |
| *AKR1C2* | -2,2012124 | -1,8511689 |
| *DCUN1D5* | -2,1943314 | -2,7006908 |
| *MRPL50* | -2,1833276 | -1,6153347 |
| *MAGEA6* | -2,1584168 | -1,7292968 |
| *GCNT2* | -2,1172331 | -2,985141 |
| *TOMM34* | -2,1038455 | -1,6251865 |
| *AP1S3* | -2,0700299 | -1,8722033 |
| *HYOU1* | -2,0623465 | -1,6013213 |
| *NSFP1* | -2,0344346 | -2,0350566 |
| *HMOX2* | -2,0327236 | -1,6810431 |
| *RAB23* | -2,012737 | -1,8136035 |
| *VDAC1P2* | -2,0085069 | -2,191307 |
| *NDUFB3* | -1,9914865 | -1,6864719 |
| *NAP1L5* | -1,9738097 | -1,6353563 |
| *DLD* | -1,9612687 | -1,6411063 |
| *UQCR10* | -1,9522292 | -1,6346784 |
| *OAZ1* | -1,9423861 | -2,0674807 |
| *MYDGF* | -1,9327821 | -1,8257903 |
| *VDAC1P1* | -1,9306007 | -2,1126201 |
| *VDAC1P6* | -1,9060815 | -2,0587063 |
| *UQCRFS1* | -1,9047805 | -1,9911702 |
| *POMP* | -1,898697 | -1,7984497 |
| *AAGAB* | -1,8967351 | -1,9154207 |
| *CAPZA1* | -1,8950066 | -1,7704973 |
| *SPRTN* | -1,8831634 | -2,1561614 |
| *BAK1P1* | -1,8803394 | -1,7076897 |
| *TSPAN13* | -1,8789401 | -3,4280506 |
| *HK1* | -1,875145 | -1,786189 |
| *AKIRIN1* | -1,8722568 | -1,8939286 |
| *VDAC1* | -1,8522554 | -1,980915 |
| *MPV17L2* | -1,846096 | -2,2527265 |
| *TRUB1* | -1,8021012 | -1,7046434 |
| *SLC6A9* | -1,7859359 | -3,060799 |
| *TNFSF9* | -1,7856436 | -2,1658627 |
| *HIGD1A* | -1,7856074 | -1,7142336 |
| *SPTLC3* | -1,7839965 | -2,6454622 |

| | | |
|---|---|---|
| *VDAC1P3* | -1,778333 | -2,172605 |
| *PSMB5* | -1,7637945 | -1,9695576 |
| *SYNGR2* | -1,75437 | -1,6189576 |
| *PLEKHB2* | -1,7415573 | -1,6133588 |
| *SYBU* | -1,736114 | -1,717313 |
| *AP2S1* | -1,7111012 | -2,0613328 |
| *UBE2L3* | -1,7104391 | -2,0648502 |
| *PRR13P5* | -1,6904893 | -1,8443051 |
| *ANXA6* | -1,6874265 | -2,1073458 |
| *USB1* | -1,686832 | -1,7613666 |
| *ZBTB8OS* | -1,6828463 | -1,5836728 |
| *PRR13* | -1,678916 | -1,7948811 |
| *SPR* | -1,6787473 | -1,7287913 |
| *TAF13* | -1,6663201 | -2,1347666 |
| *MSRB1* | -1,6620643 | -1,6075614 |
| *SNHG9* | -1,6566437 | -2,3001818 |
| *FBXO33* | -1,6494546 | -1,6287821 |
| *BAK1* | -1,6396177 | -1,9364122 |
| *GDE1* | -1,6391779 | -1,8525386 |
| *PKM* | -1,6279668 | -1,7889959 |
| *NNMT* | -1,6264894 | -1,9036062 |
| *SEC61B* | -1,6217001 | -1,5940368 |
| *SMIM3* | -1,6156551 | -1,6611333 |
| *RIPK2* | -1,5994108 | -1,6333596 |
| *VTA1* | -1,5947649 | -1,8574996 |
| *YWHAG* | -1,5736151 | -1,5644326 |
| *KIF3C* | -1,5706388 | -1,8413549 |
| *SEPHS2* | -1,5640373 | -1,8577836 |
| *RP2* | -1,560089 | -2,0977272 |
| *TMEM167A* | -1,5568779 | -1,9590007 |
| *QDPR* | -1,5568404 | -1,5939022 |
| *LSM1* | -1,5366313 | -1,7908667 |
| *CDC42* | -1,5156383 | -1,575014 |

**Table S1. Differentially expressed genes commonly downregulated in INK128 treated A549 and SK-Mel-103 cells with a p value < 0.05.**

| Genes | SK-Mel-103 INK128/Prolif FC | A549 INK128/Prolif FC |
|---|---|---|
| *CPEB1-AS1* | 24,2342795 | 5,53645618 |
| *ID2-AS1* | 20,1951415 | 3,89517065 |
| *AMY1C* | 17,9996967 | 18,0147421 |
| *KLF3-AS1* | 17,8083872 | 16,034119 |
| *AMY1B* | 17,5306686 | 11,4415277 |
| *AMY1A* | 16,8995157 | 10,6765263 |
| *AMY2A* | 14,2604424 | 19,3622899 |
| *AMY2B* | 13,7347458 | 17,8622349 |
| *DET1* | 13,4741439 | 3,2556709 |
| *CARF* | 12,8002879 | 5,84156183 |
| *FAM167B* | 12,2406551 | 5,6834603 |
| *LINC01422* | 11,9902889 | 18,3837079 |
| *FOXO1* | 11,8847781 | 3,1608634 |
| *ATP6AP1L* | 9,61268584 | 5,26345592 |
| *CCDC18-AS1* | 9,24101403 | 6,11731831 |
| *POLR2J4* | 8,39310629 | 5,40319964 |
| *PLA2G6* | 8,26266994 | 4,97340325 |
| *SALL4* | 7,91889603 | 4,1701324 |
| *MUC20* | 7,76716949 | 5,23421555 |
| *INTS6-AS1* | 7,72290359 | 4,15086472 |
| *PRRT2* | 7,60857476 | 7,18467882 |
| *ACRBP* | 6,86673907 | 15,2413831 |
| *TFAP2E* | 6,77476204 | 4,73311017 |
| *SBK1* | 6,42062349 | 2,99292837 |
| *HSF4* | 6,37273969 | 3,95909458 |
| *ADIRF-AS1* | 6,13501687 | 2,07978674 |
| *TMEM9B-AS1* | 6,07867151 | 5,20873746 |
| *LINC01089* | 5,51330068 | 1,72041454 |
| *TAS2R4* | 5,32036097 | 3,53183687 |
| *SPATA25* | 5,13052635 | 2,39949219 |
| *ARTN* | 4,93704632 | 7,85898724 |
| *GABPB1-AS1* | 4,8684129 | 2,63824214 |
| *PAXBP1-AS1* | 4,86336239 | 2,11093479 |
| *GABBR1* | 4,82927447 | 4,43192781 |
| *CNKSR1* | 4,82079753 | 2,51001945 |
| *MTMR9LP* | 4,78844131 | 2,46177297 |
| *PITPNM3* | 4,61664916 | 6,20000418 |
| *KCNIP2* | 4,5680724 | 3,30760667 |
| *FAAH* | 4,50640539 | 2,86470875 |
| *TPTE2P1* | 4,49759486 | 3,6770412 |
| *FNBP1P1* | 4,46613163 | 2,44729645 |

| | | |
|---|---|---|
| *RAB4B* | 4,35704312 | 2,40030932 |
| *TPT1-AS1* | 4,08996196 | 2,18316183 |
| *LYPD1* | 3,86791454 | 6,42759755 |
| *PAN3-AS1* | 3,85605476 | 2,01560047 |
| *CACNG8* | 3,82421129 | 1,69823733 |
| *RPL12P47* | 3,64501041 | 3,77426341 |
| *RAD51-AS1* | 3,63899819 | 2,9235331 |
| *PAN2* | 3,63493681 | 1,80362563 |
| *INE1* | 3,4339479 | 3,2631199 |
| *CROCCP3* | 3,42722712 | 2,45650393 |
| *FZD1* | 3,29078875 | 1,59228184 |
| *CSPG4P12* | 3,28206478 | 2,67680227 |
| *MORF4L2-AS1* | 3,21935086 | 1,91537909 |
| *CD22* | 3,20380042 | 3,50163016 |
| *DRICH1* | 3,15753634 | 5,63021183 |
| *ADAM8* | 3,14730785 | 2,63296608 |
| *JMJD7-PLA2G4B* | 3,13833491 | 2,95599262 |
| *ANKRD23* | 3,06638352 | 2,84710383 |
| *ITGA10* | 3,05892707 | 2,61347919 |
| *APBB3* | 3,05685283 | 3,24340949 |
| *SLC25A25-AS1* | 3,04886115 | 2,61617377 |
| *UBAP1L* | 3,01663075 | 2,19957634 |
| *EFCAB13* | 2,96270534 | 2,29666788 |
| *PILRA* | 2,93241945 | 2,09971804 |
| *FAM229A* | 2,93042906 | 3,26024704 |
| *LENG8-AS1* | 2,90311935 | 3,451622 |
| *MAPK13* | 2,85111436 | 5,54173682 |
| *ZNF337* | 2,84393409 | 1,68950568 |
| *ATG16L2* | 2,8412671 | 3,65119794 |
| *ZNF34* | 2,83768368 | 1,81034035 |
| *MIR186* | 2,82916886 | 12,5293747 |
| *KCNQ1OT1* | 2,80576493 | 2,09401832 |
| *RPS4XP16* | 2,7700254 | 2,7082101 |
| *HCG25* | 2,70696866 | 2,28583571 |
| *TRIM66* | 2,69294451 | 2,34492991 |
| *ALOX12-AS1* | 2,68168341 | 1,80903109 |
| *ST3GAL3* | 2,65833182 | 2,51302639 |
| *SPRY2* | 2,65778982 | 2,36823774 |
| *ZNF333* | 2,65562986 | 2,6894615 |
| *CCDC57* | 2,64687404 | 2,12210163 |
| *NAIP* | 2,6432458 | 2,92226854 |
| *TPT1P4* | 2,63002737 | 2,85194479 |

| | | |
|---|---|---|
| *TPH1* | 2,52180904 | 12,8407759 |
| *SH3BP5-AS1* | 2,50652842 | 2,87099045 |
| *MAN2C1* | 2,48381189 | 1,58702229 |
| *GOLGA8A* | 2,44223077 | 4,05901083 |
| *RNU6-26P* | 2,42264998 | 4,97461775 |
| *RPL12P12* | 2,41931141 | 2,51465406 |
| *SLC25A21-AS1* | 2,39588583 | 2,76460607 |
| *SZT2* | 2,38365001 | 1,70143274 |
| *LENG8* | 2,34710454 | 2,58704794 |
| *NPIPB15* | 2,33523629 | 3,2379756 |
| *ERMARD* | 2,31025594 | 1,7358938 |
| *ANKRD20A17P* | 2,30657236 | 3,05956899 |
| *CCNL2* | 2,3032551 | 2,26119381 |
| *CATSPER2* | 2,28688564 | 2,06247528 |
| *PPP1R3E* | 2,28494708 | 2,3895674 |
| *RPL32P3* | 2,28423427 | 1,68453437 |
| *RPL12P8* | 2,25497819 | 2,3059021 |
| *CAPRIN2* | 2,2445327 | 2,36373504 |
| *WNK4* | 2,22741662 | 2,41613191 |
| *RPS18* | 2,21202686 | 2,28015772 |
| *RPL13P12* | 2,2101826 | 2,10209748 |
| *RPL12P41* | 2,20517645 | 3,04971932 |
| *CDK5RAP3* | 2,19003198 | 3,04508173 |
| *RPL12P4* | 2,18955489 | 2,51293399 |
| *RPL12P17* | 2,18647617 | 2,39554442 |
| *EEF1A1P38* | 2,18509911 | 2,08425928 |
| *RPL12* | 2,18224747 | 2,48417265 |
| *RPL13* | 2,16564005 | 2,00932378 |
| *RPL37* | 2,16284933 | 2,10362285 |
| *TRIM52* | 2,15730138 | 1,75124134 |
| *N4BP2L2-IT2* | 2,14020476 | 2,54684458 |
| *RPL12P6* | 2,10823944 | 2,52956322 |
| *CLDN15* | 2,10504254 | 1,83728275 |
| *LINC00342* | 2,10229434 | 6,38901918 |
| *EEF1A1P25* | 2,0999088 | 2,01144701 |
| *PABPC1L* | 2,09523744 | 2,01604535 |
| *MYH3* | 2,09397143 | 2,13039596 |
| *RPL37P2* | 2,08750181 | 2,21716583 |
| *DNHD1* | 2,07522757 | 2,14743807 |
| *PHC1* | 2,07250684 | 1,93170214 |
| *WDR27* | 2,06008938 | 1,95384961 |
| *RPL36AP13* | 2,05543296 | 2,14853707 |

| | | |
|---|---|---|
| *POLM* | 2,05376804 | 1,80033914 |
| *GAS5* | 2,05086612 | 2,95128175 |
| *RPL12P38* | 2,04314291 | 2,51550277 |
| *NPIPB1P* | 2,01654745 | 2,39272306 |
| *RPL37P23* | 2,00704253 | 2,18306111 |
| *OGT* | 1,99385392 | 1,92778348 |
| *RPS18P12* | 1,99384761 | 2,04165366 |
| *NPIPA7* | 1,99119658 | 2,01991766 |
| *ANKRD10-IT1* | 1,98248866 | 2,29964591 |
| *RPL28* | 1,98144442 | 2,53104638 |
| *NPIPA3* | 1,96447342 | 1,92511217 |
| *RPL3P4* | 1,95647722 | 2,13664472 |
| *RPL29P12* | 1,95543672 | 2,10855564 |
| *NPIPA5* | 1,95177086 | 1,92750327 |
| *NPIPA1* | 1,95031618 | 1,90240094 |
| *RPL36AP26* | 1,94253962 | 1,86721476 |
| *CROCC* | 1,94136687 | 2,1362374 |
| *EEF1A1P12* | 1,94077202 | 1,91522493 |
| *RPS4XP13* | 1,93461096 | 1,83352687 |
| *RPL31* | 1,93209745 | 2,12347794 |
| *NPIPB11* | 1,93037885 | 2,47052029 |
| *GTF2IP13* | 1,92832736 | 2,12489305 |
| *PARP6* | 1,91221705 | 1,72136651 |
| *DNASE1L2* | 1,91028426 | 2,52022614 |
| *RPL4P2* | 1,90450402 | 1,95530039 |
| *EPB41L4A-AS1* | 1,89790007 | 3,18506038 |
| *RPL31P49* | 1,8960404 | 2,40523474 |
| *GIGYF1* | 1,89592951 | 2,23987372 |
| *C14orf93* | 1,88628708 | 2,40480075 |
| *RFX3* | 1,88409394 | 1,97282182 |
| *MDM4* | 1,88111102 | 1,67761315 |
| *EEF1A1P9* | 1,88001664 | 1,82688466 |
| *ZBTB26* | 1,87322276 | 1,83152815 |
| *RPL13AP3* | 1,86963332 | 2,05929017 |
| *EZH1* | 1,86699189 | 2,21909936 |
| *NFYC-AS1* | 1,86060231 | 2,17961906 |
| *NPIPP1* | 1,85042187 | 2,21104527 |
| *EEF1A1P29* | 1,8445573 | 1,83278301 |
| *ZNF839* | 1,843048 | 1,53229868 |
| *NPIPB4* | 1,83494767 | 2,29841691 |
| *RPL4P3* | 1,83221565 | 1,84637753 |
| *NPIPB3* | 1,82986307 | 2,25291764 |

| | | |
|---|---|---|
| *CREB3L4* | 1,82891921 | 2,35852527 |
| *HOXB-AS1* | 1,8266154 | 2,08563977 |
| *PSMA3-AS1* | 1,82124718 | 2,20784803 |
| *IQCH-AS1* | 1,82077198 | 2,33031786 |
| *RPS27* | 1,81941922 | 2,05916139 |
| *NPIPB5* | 1,81916414 | 2,33833055 |
| *MAGI2-AS3* | 1,81015507 | 2,20411274 |
| *ANKDD1A* | 1,80054771 | 1,8799092 |
| *ZBTB12* | 1,79969598 | 1,9127619 |
| *KANTR* | 1,79787388 | 1,9853246 |
| *RPL10P3* | 1,78513931 | 2,1119778 |
| *LIME1* | 1,78500943 | 1,83942897 |
| *RPL4P4* | 1,7829766 | 1,74302007 |
| *RPL3* | 1,78050098 | 1,88144378 |
| *RPL18A* | 1,77861071 | 1,84934655 |
| *EEF1A1P11* | 1,77667424 | 1,8035803 |
| *RPS6* | 1,77200135 | 1,79421894 |
| *MSANTD2* | 1,7677111 | 1,83083941 |
| *ANXA2R* | 1,76568712 | 1,66712376 |
| *ANKMY1* | 1,75988528 | 1,62863711 |
| *RPL18AP3* | 1,7490507 | 1,85838692 |
| *PIM1* | 1,74766427 | 3,08987371 |
| *RPS4X* | 1,74610672 | 1,74536286 |
| *DDX39B* | 1,74440436 | 1,71249274 |
| *SPTB* | 1,74189502 | 2,77985027 |
| *VMP1* | 1,74023315 | 1,96008816 |
| *RPL29* | 1,73738987 | 1,77817799 |
| *EEF1A1P14* | 1,73032175 | 1,82264002 |
| *RPL13AP6* | 1,72886336 | 1,917851 |
| *DMTF1* | 1,72718484 | 1,88987874 |
| *PPP1R15A* | 1,72581599 | 2,33979423 |
| *USP32P3* | 1,72245628 | 2,86315585 |
| *RPL10* | 1,72240392 | 1,88761753 |
| *LRRC37A17P* | 1,71696937 | 1,69940924 |
| *ACVR2B* | 1,71567473 | 3,01098149 |
| *RPS4XP3* | 1,70523389 | 1,72638469 |
| *RPL4P5* | 1,70386352 | 1,68077617 |
| *GUSBP11* | 1,70361439 | 2,36484164 |
| *RPL29P11* | 1,70042507 | 1,7447621 |
| *STRADA* | 1,69704774 | 1,62589377 |
| *RPL4* | 1,68611108 | 1,69577586 |
| *RPL9P32* | 1,68049015 | 1,76019619 |

| | | |
|---|---|---|
| *RPS27P29* | 1,67916371 | 2,08590891 |
| *RPS16* | 1,67705319 | 1,71788555 |
| *RPS3AP49* | 1,67131274 | 1,8474765 |
| *RPS8* | 1,66745247 | 1,85189089 |
| *THUMPD3-AS1* | 1,66053025 | 1,68296976 |
| *RPL23AP43* | 1,65971417 | 1,8806341 |
| *RPL23AP18* | 1,64552919 | 1,66536341 |
| *EEF1A1P7* | 1,64510603 | 1,62586736 |
| *RPS3AP47* | 1,64139269 | 1,68165385 |
| *RPS3AP26* | 1,64022946 | 1,77202126 |
| *RPL3P2* | 1,63510676 | 1,72345501 |
| *RPS15AP17* | 1,63486796 | 1,99214037 |
| *CCDC144B* | 1,63082239 | 2,13324857 |
| *KLF7* | 1,62987171 | 2,35505806 |
| *RPS3AP25* | 1,62843685 | 1,72512115 |
| *RPS3AP5* | 1,62240096 | 1,70759976 |
| *LRRC37A2* | 1,61300641 | 1,99913129 |
| *ZNF891* | 1,61119161 | 1,82183111 |
| *RPS3AP6* | 1,60794749 | 1,71100085 |
| *RPS3A* | 1,6071584 | 1,69960289 |
| *EEF1A1P19* | 1,59932223 | 1,66485517 |
| *LRRC37A3* | 1,59809169 | 2,45173733 |
| *EEF1A1* | 1,59506113 | 1,69388223 |
| *GTF2IP20* | 1,59448314 | 2,40941891 |
| *LRRC37A* | 1,58400855 | 1,96160487 |
| *RPL3P7* | 1,58270507 | 1,77166157 |
| *RPL9* | 1,58158717 | 1,69154891 |
| *RPL23AP74* | 1,5813112 | 1,70212635 |
| *HSPBAP1* | 1,57937918 | 1,74357788 |
| *RPL26P19* | 1,56949012 | 2,18188302 |
| *RPL9P7* | 1,56872442 | 1,69681715 |
| *EEF1A1P6* | 1,56631885 | 1,65756877 |
| *LRRC37A4P* | 1,55236363 | 1,65401092 |
| *EEF1A1P5* | 1,54804357 | 1,63287737 |
| *MYO9A* | 1,54802053 | 1,5891383 |
| *EEF2* | 1,5407837 | 1,9479163 |
| *RPL23AP42* | 1,53744015 | 1,70520604 |
| *RPL23A* | 1,53358374 | 1,72262166 |
| *UBALD2* | 1,53167876 | 2,10542168 |
| *RPL23AP2* | 1,51864274 | 1,71343423 |
| *TBCE* | 1,34874722 | 2,65485572 |

**Table S2. Differentially expressed genes commonly upregulated in INK128-treated A549 and SK-Mel-103 cells with a p value < 0.05.**

| Genes | Prolif doxy/prolif vehicle | Genes | INK128 doxy/INK128 vehicle |
|---|---|---|---|
| *Clec2l* | -2,2259618 | *Clec2l* | -1,8359515 |
| *Susd2* | -2,4260122 | *Susd2* | -1,228338214 |
| *Heatr1* | -2,215191 | *Heatr1* | -2,458532519 |
| *Gm5480* | -1,9523131 | *Gm5480* | -1,664293992 |
| *BC068157* | -1,9208461 | *BC068157* | -3,093179932 |
| *Rdh11* | -1,9147214 | *Rdh11* | -1,840681791 |
| *C3* | -1,9090531 | *C3* | -2,003610212 |
| *Mal* | -1,8144853 | *Mal* | -1,07870848 |
| *Agpat4* | -1,7164383 | *Agpat4* | -1,346233225 |
| *Snrpe* | -1,6850692 | *Snrpe* | -1,028515047 |
| *Hcrtr1* | -1,6499643 | *Hcrtr1* | -1,062663983 |
| *Olfr976* | -1,6278538 | *Olfr976* | -1,636135808 |
| *Zfp280d* | -1,5704605 | *Zfp280d* | -1,949860241 |
| *Efcab4b* | -1,5664993 | *Efcab4b* | -1,088080674 |
| *Wfdc3* | -1,5601252 | *Wfdc3* | -1,926303142 |
| *Snrpc* | -1,5506222 | *Snrpc* | -1,492322326 |
| *Eif3b* | -1,5320306 | *Eif3b* | -1,956813922 |
| *Wfdc12* | -1,5272456 | *Wfdc12* | -1,106651344 |
| *Med16* | -1,5254836 | *Med16* | -3,518514751 |
| *Xkr6* | -1,4578114 | *Xkr6* | -1,156082548 |
| *Lta4h* | -1,4312302 | *Lta4h* | -1,290630698 |
| *Sec16b* | -1,4184383 | *Sec16b* | -1,671232626 |
| *Gpr88* | -1,4037135 | *Gpr88* | -1,087086381 |
| *Kpna3* | -1,3657215 | *Kpna3* | -1,496088262 |
| *Olfr299* | -1,3656401 | *Olfr299* | -1,705725762 |
| *Rexo2* | -1,3506939 | *Rexo2* | -1,39943368 |
| *Ptpdc1* | -1,3408801 | *Ptpdc1* | -2,94442154 |
| *Pax5* | -1,3216299 | *Pax5* | -1,516207537 |
| *Prss16* | -1,2896296 | *Prss16* | -1,832350982 |
| *Btbd9* | -1,2880855 | *Btbd9* | -1,560822986 |
| *4933404M02Rik* | -1,2828483 | *4933404M02Rik* | -1,146714539 |
| *Hist1h3e* | -1,2691891 | *Hist1h3e* | -1,19979842 |
| *Epha8* | -1,2575906 | *Epha8* | -1,490733459 |
| *Mrpl15* | -1,2485118 | *Mrpl15* | -1,061468816 |
| *Clec1b* | -1,2427276 | *Clec1b* | -1,078148035 |
| *Vmn1r57* | -1,229581 | *Vmn1r57* | -1,35591202 |
| *Cend1* | -1,2240933 | *Cend1* | -1,621798242 |
| *Fam186b* | -1,2184624 | *Fam186b* | -1,453671095 |
| *Vmn1r204* | -1,2137116 | *Vmn1r204* | -1,435176415 |
| *Rgs7* | -1,2101355 | *Rgs7* | -1,136740054 |
| *Lsm1* | -1,2042396 | *Lsm1* | -1,661337237 |

| | | | |
|---|---|---|---|
| *Tut1* | -1,1800495 | *Tut1* | -1,035008877 |
| *Mepce* | -1,1746234 | *Mepce* | -1,766245455 |
| *Fcgr2b* | -1,1683708 | *Fcgr2b* | -1,15213707 |
| *Cdca4* | -1,1542511 | *Cdca4* | -1,886232182 |
| *Auh* | -1,1433256 | *Auh* | -1,578545271 |
| *Olfr996* | -1,139033 | *Olfr996* | -1,262447732 |
| *Ppp1r3f* | -1,1380471 | *Ppp1r3f* | -1,813839453 |
| *Oas1g* | -1,1326693 | *Oas1g* | -1,296471528 |
| *Sh2d2a* | -1,1257063 | *Sh2d2a* | -1,202159359 |
| *Rbck1* | -1,1233365 | *Rbck1* | -1,938783539 |
| *Fkbp2* | -1,1174667 | *Fkbp2* | -1,32639463 |
| *Tubb3* | -1,1166952 | *Tubb3* | -1,532399618 |
| *Sstr1* | -1,1051363 | *Sstr1* | -1,43014548 |
| *Dpf2* | -1,1019453 | *Dpf2* | -2,570751454 |
| *Eif2b5* | -1,097712 | *Eif2b5* | -1,159108712 |
| *Sip1* | -1,0883433 | *Sip1* | -1,660304424 |
| *1810065E05Rik* | -1,0808824 | *1810065E05Rik* | -1,599729611 |
| *Serpinb6a* | -1,0775348 | *Serpinb6a* | -1,046032122 |
| *Fezf2* | -1,0732216 | *Fezf2* | -1,259542943 |
| *Zcchc10* | -1,0652578 | *Zcchc10* | -1,056214747 |
| *Cdh17* | -1,060942 | *Cdh17* | -1,110243622 |
| *Gm17677* | -1,057955 | *Gm17677* | -1,111893186 |
| *4933403F05Rik* | -1,0536968 | *4933403F05Rik* | -1,009618691 |
| *Gm2016* | -1,0500275 | *Gm2016* | -2,043087348 |
| *BC018507* | -1,0495755 | *BC018507* | -1,563248543 |
| *Pkp4* | -1,0489108 | *Pkp4* | -1,249156146 |
| *Id3* | -1,0434105 | *Id3* | -1,994185612 |
| *Tfdp2* | -1,0399493 | *Tfdp2* | -1,101853746 |
| *2410137M14Rik* | -1,0392727 | *2410137M14Rik* | -1,641237032 |
| *Evx1* | -1,038284 | *Evx1* | -2,368618341 |
| *Cyp3a25* | -1,0345478 | *Cyp3a25* | -1,101174989 |
| *Kank4* | -1,0320466 | *Kank4* | -1,658151004 |
| *Foxo4* | -1,0318349 | *Foxo4* | -1,106607655 |
| *Rpl41* | -1,0307546 | *Rpl41* | -1,054224302 |
| *Gm12886* | -1,0234699 | *Gm12886* | -1,338620613 |
| *Zscan4e* | -1,0211416 | *Zscan4e* | -1,454396864 |
| *Syde2* | -1,0182281 | *Syde2* | -1,549960879 |
| *Epo* | -1,0090712 | *Epo* | -1,849161898 |
| *Kdelr2* | -1,0042682 | *Kdelr2* | -1,473868438 |
| *Amd2* | -1,0032321 | *Amd2* | -1,030559403 |
| *Neurl1a* | -1,0017544 | *Neurl1a* | -1,770012011 |
| *Prl2c5* | -1,0003802 | *Prl2c5* | -1,129555299 |

| Genes | Prolif doxy/prolif vehicle |
|---|---:|
| *Prmt7* | -2,548594122 |
| *Gm6792* | -2,191432098 |
| *Dppa2* | -1,926612158 |
| *Bcl2a1c* | -1,896883151 |
| *1700001C19Rik* | -1,825070136 |
| *Zfp354c* | -1,810242074 |
| *Arl6ip6* | -1,75432396 |
| *Klra10* | -1,642107571 |
| *Pde3b* | -1,623887866 |
| *Vmn2r33* | -1,597875518 |
| *2810432D09Rik* | -1,594384238 |
| *Bnip3* | -1,575495692 |
| *Defb8* | -1,550500853 |
| *Tsc22d2* | -1,542891035 |
| *Qpct* | -1,520667635 |
| *Krtap17-1* | -1,51585507 |
| *Emp3* | -1,514883208 |
| *Cul4b* | -1,485861849 |
| *Sgta* | -1,4842016 |
| *Zfp518a* | -1,48348241 |
| *Gm6484* | -1,477639462 |
| *Gnl1* | -1,475979548 |
| *Olfr1380* | -1,462849751 |
| *Fgd2* | -1,448106166 |
| *Gm12789* | -1,444683717 |
| *Cenpk* | -1,444355556 |
| *Selk* | -1,438566816 |
| *Vmn1r159* | -1,425589419 |
| *Zfp652* | -1,42224486 |
| *Six4* | -1,4096216 |
| *Oxtr* | -1,409541997 |
| *Wbp1* | -1,406348611 |
| *Gm9781* | -1,396583289 |
| *Entpd2* | -1,384511148 |
| *Fam164c* | -1,381015749 |
| *Zbed4* | -1,379740664 |
| *Spata20* | -1,37923664 |
| *Trim43b* | -1,37484012 |
| *Rpl17* | -1,361476389 |
| *Taf5l* | -1,358609783 |
| *Iqck* | -1,352186016 |

| | |
|---|---|
| *Olfr1392* | -1,345600261 |
| *Pxn* | -1,343596739 |
| *Snai1* | -1,340625067 |
| *Ammecr1* | -1,339081457 |
| *Gm17252* | -1,336291493 |
| *H13* | -1,332824825 |
| *Vwf* | -1,32793923 |
| *Snrpd3* | -1,325942699 |
| *Gm14393* | -1,321007662 |
| *Kcnk3* | -1,318543582 |
| *Alg10b* | -1,317549282 |
| *Aasdhppt* | -1,316632084 |
| *Sh3tc2* | -1,311178493 |
| *Gm4858* | -1,310979025 |
| *Ppp1r9a* | -1,309778146 |
| *Xrcc1* | -1,306666811 |
| *Agxt2l2* | -1,303088352 |
| *Ufsp1* | -1,293519005 |
| *Fads3* | -1,289737159 |
| *4930572J05Rik* | -1,289679299 |
| *Megf9* | -1,288902906 |
| *Mageb16* | -1,288023121 |
| *Usp20* | -1,285717002 |
| *Rfwd2* | -1,284112966 |
| *Rtkn* | -1,26946346 |
| *Zfp114* | -1,263522484 |
| *Cdr1* | -1,259074971 |
| *Gnb1l* | -1,257841395 |
| *Creg2* | -1,254414239 |
| *Cdh15* | -1,253150455 |
| *Dusp6* | -1,252408415 |
| *Defb9* | -1,252403018 |
| *St3gal4* | -1,252224418 |
| *Ifrd2* | -1,251189135 |
| *Ikzf1* | -1,248910528 |
| *Rft1* | -1,248780313 |
| *Il17a* | -1,247951146 |
| *Foxred1* | -1,240535358 |
| *Wibg* | -1,229358173 |
| *Efna1* | -1,228214761 |
| *Syt1* | -1,225554942 |
| *Gnaq* | -1,223041614 |

| | |
|---|---|
| *Hopx* | -1,222990775 |
| *Mmp23* | -1,221703748 |
| *Rbm28* | -1,218317814 |
| *Hist1h3c* | -1,218046772 |
| *Fam83g* | -1,217817927 |
| *Gm10100* | -1,217657351 |
| *Prr23a* | -1,217369297 |
| *Il5* | -1,215450036 |
| *Plekhg2* | -1,20916463 |
| *Tmem54* | -1,206159138 |
| *Ghitm* | -1,205562882 |
| *A630055G03Rik* | -1,201791529 |
| *A930005I04Rik* | -1,197058485 |
| *1700030K09Rik* | -1,195988574 |
| *Ttc14* | -1,194612853 |
| *Cables1* | -1,191536145 |
| *Hpdl* | -1,18397314 |
| *H2-T3* | -1,178812143 |
| *Fndc7* | -1,177818144 |
| *Sap30* | -1,17743254 |
| *Tspyl2* | -1,177298903 |
| *Map2k2* | -1,173194738 |
| *Rnf144b* | -1,171645073 |
| *Npepl1* | -1,167245534 |
| *Stra8* | -1,164391085 |
| *Gfm2* | -1,164032625 |
| *Batf* | -1,161012926 |
| *Gm6760* | -1,159156474 |
| *Ccdc12* | -1,156647015 |
| *Neurod2* | -1,15613372 |
| *Ocln* | -1,155111297 |
| *Efr3b* | -1,154089835 |
| *Pin4* | -1,152107539 |
| *1810031K17Rik* | -1,14733296 |
| *Cntn6* | -1,147058362 |
| *Rnf183* | -1,14450468 |
| *Cenpn* | -1,14421669 |
| *Gm8267* | -1,143422278 |
| *Hhla1* | -1,142085589 |
| *Dclk3* | -1,134491282 |
| *Ucn3* | -1,132637959 |
| *Fpr-rs6* | -1,13120663 |

| | |
|---|---|
| *Cys1* | -1,127147423 |
| *Rabggtb* | -1,125197856 |
| *Rela* | -1,123575798 |
| *Naa50* | -1,121963397 |
| *Slc31a1* | -1,121122619 |
| *1810062G17Rik* | -1,12083318 |
| *Pgc* | -1,119913069 |
| *Nat15* | -1,11850996 |
| *Slc6a5* | -1,118116883 |
| *Usp49* | -1,117027524 |
| *Prrt2* | -1,11701729 |
| *Pgbd5* | -1,116347917 |
| *Gpn3* | -1,115113726 |
| *Tmem143* | -1,110507448 |
| *Gm6320* | -1,105787516 |
| *Pdcd2* | -1,105285865 |
| *Gm19345* | -1,103149972 |
| *Zc3h12c* | -1,102303941 |
| *Sco1* | -1,101917762 |
| *Papln* | -1,101637661 |
| *Ms4a8a* | -1,098502678 |
| *Aldh1a2* | -1,098204466 |
| *E2f5* | -1,096031283 |
| *Cep120* | -1,096028889 |
| *Fam19a4* | -1,095572729 |
| *Ano8* | -1,095498349 |
| *Atpaf2* | -1,095280055 |
| *Vnn3* | -1,09479856 |
| *Slc22a18* | -1,091709329 |
| *Ms4a3* | -1,089346423 |
| *Wsb2* | -1,088408326 |
| *Ephb3* | -1,088309849 |
| *Purb* | -1,084395775 |
| *Spert* | -1,08230107 |
| *Ctla2b* | -1,080688461 |
| *Cldn12* | -1,073565191 |
| *Cnot7* | -1,071330177 |
| *Cacna1a* | -1,070546033 |
| *Tceal5* | -1,068623554 |
| *Dnm1l* | -1,067485061 |
| *Acacb* | -1,066810698 |
| *Bod1* | -1,065378399 |

| | |
|---|---|
| *Gxylt2* | -1,064226907 |
| *Col9a1* | -1,06314608 |
| *Zfp13* | -1,061609883 |
| *1700125D06Rik* | -1,05917414 |
| *Tmem43* | -1,057408176 |
| *Arcn1* | -1,057345374 |
| *Lrrn1* | -1,057113311 |
| *Apoa4* | -1,054016658 |
| *Fsbp* | -1,051700313 |
| *Zfp938* | -1,05158862 |
| *Eif4e1b* | -1,05122624 |
| *Slc13a5* | -1,050989924 |
| *Adam11* | -1,050796305 |
| *Naf1* | -1,050356489 |
| *AA986860* | -1,048635739 |
| *Srp68* | -1,047933654 |
| *Vmn1r170* | -1,046612585 |
| *Olfr113* | -1,046169235 |
| *Cldn3* | -1,043010686 |
| *Fam161b* | -1,041133664 |
| *Eif3l* | -1,041119312 |
| *Leap2* | -1,0406304 |
| *Ppara* | -1,039709366 |
| *Trim28* | -1,038519356 |
| *Rrp9* | -1,038337041 |
| *Lrrc8b* | -1,037629616 |
| *Megf6* | -1,03624078 |
| *Eef1a2* | -1,035310416 |
| *Zfp956* | -1,035013125 |
| *Pex11a* | -1,034514677 |
| *Gpatch1* | -1,034395626 |
| *Nudcd3* | -1,034324193 |
| *1700101E01Rik* | -1,033598513 |
| *Cyp2f2* | -1,029728086 |
| *Setd8* | -1,027306431 |
| *Dtl* | -1,026521152 |
| *Gabpa* | -1,026406762 |
| *Mocs3* | -1,025547845 |
| *Gm597* | -1,022653583 |
| *Mlxipl* | -1,022481751 |
| *2410091C18Rik* | -1,02223465 |
| *Cmpk2* | -1,021331842 |

| | |
|---|---|
| *Brd9* | -1,021004685 |
| *D4Wsu53e* | -1,019458138 |
| *Capns1* | -1,016317287 |
| *Sohlh1* | -1,015844141 |
| *Rbl1* | -1,0157282 |
| *Elovl1* | -1,014787002 |
| *4732415M23Rik* | -1,014585757 |
| *Dis3l* | -1,012754679 |
| *Aurka* | -1,012286822 |
| *2700078E11Rik* | -1,011732003 |
| *Trat1* | -1,010884642 |
| *Fes* | -1,01013627 |
| *Fbxl6* | -1,008940252 |
| *Olfr314* | -1,00831335 |
| *Ccdc11* | -1,008279004 |
| *Lcp2* | -1,004653077 |
| *Krtap5-4* | -1,003338169 |
| *Gm8298* | -1,002343369 |
| *Arl4d* | -1,001943266 |
| *Mau2* | -1,001511201 |

| Genes | INK128 doxy/INK128 vehicle |
|---|---|
| *Prdx1* | -3,33404503 |
| *H2afz* | -3,086989037 |
| *Vmn1r121* | -2,952586346 |
| *Rassf5* | -2,684912477 |
| *Rbm14* | -2,539009795 |
| *9230110F15Rik* | -2,523205952 |
| *Rnf24* | -2,492343092 |
| *Hmox1* | -2,470016295 |
| *Tuba1b* | -2,456671762 |
| *Rybp* | -2,429233757 |
| *Rassf1* | -2,406218593 |
| *Edaradd* | -2,364712838 |
| *Supt5h* | -2,328265506 |
| *Ppp2r3c* | -2,317056982 |
| *Cartpt* | -2,271414806 |
| *Notch1* | -2,257691848 |
| *Cnot8* | -2,24667952 |
| *Pip5kl1* | -2,24025078 |
| *Rnf2* | -2,222507232 |
| *Iars2* | -2,211772257 |
| *Ctbp1* | -2,208471161 |
| *Slc2a13* | -2,173618323 |
| *Wt1* | -2,168956329 |
| *Ptpn3* | -2,163807143 |
| *Itpa* | -2,13811481 |
| *Rangrf* | -2,136085522 |
| *Parl* | -2,120782713 |
| *Prep* | -2,100831763 |
| *Osm* | -2,091714096 |
| *Acpp* | -2,09160796 |
| *Mrpl47* | -2,085277653 |
| *Aacs* | -2,074726517 |
| *4930504O13Rik* | -2,066313944 |
| *Olfr490* | -2,054816913 |
| *Dnajb4* | -2,039886936 |
| *Smap2* | -2,037069004 |
| *2410002F23Rik* | -2,004040241 |
| *Papd4* | -2,003188494 |
| *Kcnf1* | -2,002713541 |
| *Adamts13* | -1,992898901 |
| *Zfp873* | -1,987623598 |

| | |
|---|---|
| *1110038D17Rik* | -1,961210802 |
| *Glt28d2* | -1,951640325 |
| *Klraq1* | -1,946514647 |
| *Vsig2* | -1,930679189 |
| *Rab8b* | -1,902672455 |
| *Ccnb1* | -1,902516789 |
| *0610010K14Rik* | -1,896155111 |
| *Slurp1* | -1,893911486 |
| *Uhrf1* | -1,889388343 |
| *Camta1* | -1,888199933 |
| *Hnrnpk* | -1,884385151 |
| *Pcgf6* | -1,883867522 |
| *Prrt3* | -1,882980459 |
| *Cenpw* | -1,879739957 |
| *Cbx7* | -1,877834929 |
| *Olfr204* | -1,873803083 |
| *Plekhn1* | -1,869737163 |
| *Car9* | -1,85627259 |
| *Vipr2* | -1,852142081 |
| *Lmbrd1* | -1,847108334 |
| *Rnf170* | -1,846711312 |
| *Slc7a6os* | -1,831433536 |
| *Gm12769* | -1,828979688 |
| *AU021034* | -1,821453382 |
| *Olfr347* | -1,818223364 |
| *Lrrc51* | -1,815878874 |
| *Iqcg* | -1,813287855 |
| *Nccrp1* | -1,813048604 |
| *Btaf1* | -1,802820587 |
| *Tfeb* | -1,786328982 |
| *Esam* | -1,784646856 |
| *Narfl* | -1,783696306 |
| *Manf* | -1,783235558 |
| *Olfr1066* | -1,782839303 |
| *2810405K02Rik* | -1,773085745 |
| *Ubc* | -1,772641422 |
| *Sc5d* | -1,767040809 |
| *Zdhhc13* | -1,755263672 |
| *Otub1* | -1,753049943 |
| *Ccdc58* | -1,744378235 |
| *Scube3* | -1,74334633 |
| *Popdc3* | -1,742763804 |

| | |
|---|---|
| *Sirt2* | -1,742479557 |
| *Neurog1* | -1,738538339 |
| *Ncoa1* | -1,737842474 |
| *Olfr1121* | -1,736352598 |
| *Bloc1s4* | -1,735311861 |
| *Gm8909* | -1,725737848 |
| *Atl2* | -1,719983408 |
| *Meig1* | -1,71764986 |
| *1810019J16Rik* | -1,715122094 |
| *A430105I19Rik* | -1,713169208 |
| *Efhc1* | -1,705701575 |
| *Khdc3* | -1,702650935 |
| *Ak5* | -1,698084287 |
| *Tecta* | -1,692718308 |
| *Cacng5* | -1,691624596 |
| *Gna15* | -1,690444592 |
| *Tcf15* | -1,690160062 |
| *Sh2d1a* | -1,68891983 |
| *Htr1a* | -1,688251584 |
| *4933427D06Rik* | -1,685993607 |
| *Tob1* | -1,680346129 |
| *Fdx1* | -1,674290514 |
| *Map3k5* | -1,669122039 |
| *C630004H02Rik* | -1,669108384 |
| *Slc23a1* | -1,668668786 |
| *Kcmf1* | -1,667651956 |
| *Dnmt1* | -1,661907078 |
| *Uxt* | -1,658322023 |
| *2310003H01Rik* | -1,656405264 |
| *Taok2* | -1,654456951 |
| *Gm4345* | -1,65015333 |
| *Gjd3* | -1,642569707 |
| *Fasl* | -1,622216025 |
| *Ino80b* | -1,617746143 |
| *Olfr978* | -1,615103406 |
| *Ppp1r14d* | -1,611695471 |
| *Akr1b10* | -1,607786559 |
| *Dazl* | -1,607401119 |
| *Fst* | -1,606566543 |
| *Tmem95* | -1,606110827 |
| *Agmo* | -1,594476201 |
| *AW146020* | -1,593542827 |

| | |
|---|---|
| *Cxcr4* | -1,59217405 |
| *Slc43a3* | -1,591484448 |
| *Igsf9* | -1,590894272 |
| *Polg2* | -1,590768844 |
| *Fam96b* | -1,59015865 |
| *Gm10696* | -1,584760666 |
| *Ceacam15* | -1,578431887 |
| *Spaca5* | -1,573827442 |
| *Serpina3m* | -1,573377653 |
| *Elk3* | -1,572768345 |
| *Pias1* | -1,5724617 |
| *Det1* | -1,572401608 |
| *9030624G23Rik* | -1,570612848 |
| *Asb3* | -1,565518767 |
| *Dlg2* | -1,565280443 |
| *Nfe2l2* | -1,562282038 |
| *Sparcl1* | -1,559460831 |
| *Svopl* | -1,557476217 |
| *Cst12* | -1,556839985 |
| *Tns1* | -1,555243329 |
| *Pdcd1* | -1,553461597 |
| *Stx16* | -1,551950962 |
| *Tmem50b* | -1,551736811 |
| *Ddx28* | -1,551097112 |
| *Vmn2r112* | -1,545501205 |
| *Paqr5* | -1,545261771 |
| *Olfr291* | -1,541262371 |
| *Eif4ebp3* | -1,540263806 |
| *2810453I06Rik* | -1,537687813 |
| *Aicda* | -1,536086779 |
| *Pgap3* | -1,535624739 |
| *Gcsh* | -1,535191701 |
| *2010005H15Rik* | -1,534097692 |
| *Olfr1416* | -1,533914987 |
| *Igtp* | -1,52981674 |
| *Fam111a* | -1,527867654 |
| *Tac2* | -1,526857331 |
| *4930529M08Rik* | -1,523709336 |
| *Iglon5* | -1,522004451 |
| *Egfl7* | -1,520996055 |
| *Palmd* | -1,517216275 |
| *Gm221* | -1,513883195 |

| | |
|---|---|
| *Acer3* | -1,513467571 |
| *4930550L24Rik* | -1,512932502 |
| *Ly6g6d* | -1,510529637 |
| *1700013F07Rik* | -1,508835478 |
| *Mipol1* | -1,506838096 |
| *Fam118a* | -1,506339649 |
| *Pla2g4a* | -1,504637433 |
| *Efs* | -1,502646006 |
| *Cbll1* | -1,499234728 |
| *Gm15114* | -1,49897147 |
| *Gpr126* | -1,498792612 |
| *Mmp8* | -1,497803252 |
| *Clk3* | -1,496460664 |
| *Amelx* | -1,493858985 |
| *Mtr* | -1,490825772 |
| *Esyt2* | -1,49011521 |
| *Chic1* | -1,48948086 |
| *Chchd7* | -1,489294632 |
| *Nt5m* | -1,483385061 |
| *Sfpi1* | -1,481574915 |
| *Cox5a* | -1,480758122 |
| *Hrg* | -1,480747752 |
| *Inca1* | -1,479364549 |
| *Try10* | -1,478274081 |
| *Gk5* | -1,476486911 |
| *Eid2* | -1,475120308 |
| *H2-M5* | -1,47487466 |
| *Nup43* | -1,47248365 |
| *Olfr157* | -1,471770127 |
| *Rasgrp2* | -1,467538054 |
| *Pth2r* | -1,467157943 |
| *Dctn6* | -1,4650938 |
| *Rab32* | -1,465072699 |
| *Frzb* | -1,464910126 |
| *Hao1* | -1,458350531 |
| *Prss29* | -1,456537222 |
| *Bptf* | -1,455853171 |
| *Eif3g* | -1,455739383 |
| *Stc1* | -1,454553718 |
| *Cbr2* | -1,452687597 |
| *Snx21* | -1,451203704 |
| *Kng2* | -1,448650211 |

| | |
|---|---|
| *Agbl3* | -1,447013248 |
| *Mical3* | -1,446901147 |
| *Spice1* | -1,446794265 |
| *Fhl1* | -1,443886725 |
| *Tbc1d7* | -1,443565696 |
| *Krt35* | -1,443538859 |
| *Ndufa3* | -1,443179216 |
| *Hapln3* | -1,437148438 |
| *Apba2* | -1,436517629 |
| *Gm13283* | -1,435441628 |
| *Tmem223* | -1,434130066 |
| *Rpp25* | -1,432933832 |
| *Pold4* | -1,429854206 |
| *Eif3d* | -1,424923855 |
| *Dync1h1* | -1,424541414 |
| *Kctd19* | -1,42376343 |
| *Rc3h1* | -1,422296742 |
| *Serpind1* | -1,421128123 |
| *2200002D01Rik* | -1,421040162 |
| *Gm14743* | -1,420707112 |
| *Mrpl28* | -1,419593681 |
| *Olfr324* | -1,416036836 |
| *Nos3* | -1,414890218 |
| *Cyr61* | -1,413834673 |
| *Pttg1* | -1,413208513 |
| *Tecr* | -1,413125029 |
| *Echdc3* | -1,412603201 |
| *Map3k12* | -1,411489026 |
| *Dkc1* | -1,411305698 |
| *Olfr980* | -1,410358337 |
| *Zfp689* | -1,407910692 |
| *Ccl25* | -1,406644816 |
| *Larp6* | -1,406294323 |
| *Thsd7a* | -1,403092472 |
| *Naa25* | -1,402244904 |
| *Farsb* | -1,398996306 |
| *Cplx3* | -1,39842402 |
| *Olfr371* | -1,396397982 |
| *Cln5* | -1,395011355 |
| *Lasp1* | -1,391430395 |
| *mar-03* | -1,390086582 |
| *Gm13247* | -1,389391109 |

| | |
|---|---|
| *Arhgdib* | -1,386262958 |
| *Pcca* | -1,385642039 |
| *Bms1* | -1,385613946 |
| *Eif4g1* | -1,385304029 |
| *Zrsr1* | -1,380279385 |
| *Nhlrc2* | -1,379147454 |
| *Eogt* | -1,378743471 |
| *Col6a3* | -1,377850567 |
| *Camkk2* | -1,376813217 |
| *Bmp8b* | -1,376813124 |
| *Hsd11b2* | -1,376718112 |
| *Lrrc30* | -1,375414679 |
| *Olfr1295* | -1,374991077 |
| *Abcc4* | -1,374975559 |
| *Diablo* | -1,374140892 |
| *Bhmt* | -1,373281483 |
| *Ift52* | -1,372635783 |
| *Tsen34* | -1,371748821 |
| *Txnrd1* | -1,369860611 |
| *Klhl38* | -1,368032082 |
| *Jarid2* | -1,365253412 |
| *Klra4* | -1,363227229 |
| *Lpar6* | -1,360535376 |
| *Gimap4* | -1,360230534 |
| *Fam168b* | -1,359676981 |
| *Diras2* | -1,358710164 |
| *Las1l* | -1,357996889 |
| *Ncapg2* | -1,357895306 |
| *Epb4.1l4a* | -1,357615686 |
| *Cdcp1* | -1,356219102 |
| *Gtf2b* | -1,354108508 |
| *1600029D21Rik* | -1,353538206 |
| *Rrn3* | -1,349991038 |
| *Cpt1a* | -1,344426508 |
| *Pfdn5* | -1,34438152 |
| *Lrrc10b* | -1,343599362 |
| *Olfr66* | -1,342923028 |
| *Ntrk2* | -1,341497306 |
| *Aifm2* | -1,341412562 |
| *Oxsm* | -1,339083456 |
| *Dlg1* | -1,337159077 |
| *Nucb1* | -1,336313323 |

| | |
|---|---|
| *Fam59a* | -1,335782457 |
| *Olfr385* | -1,335419666 |
| *Uts2d* | -1,334551306 |
| *Col8a1* | -1,333097934 |
| *Spp1* | -1,332973487 |
| *Fbxl5* | -1,332648804 |
| *Olfr575* | -1,331937562 |
| *Gm906* | -1,330619826 |
| *Ildr1* | -1,327628293 |
| *Defb45* | -1,320450979 |
| *Zfp961* | -1,319070426 |
| *Plin5* | -1,319064561 |
| *Olfr725* | -1,317878625 |
| *Zbtb41* | -1,31700075 |
| *Gatad2a* | -1,316380172 |
| *Grhl1* | -1,316079052 |
| *Aven* | -1,308427089 |
| *Mtx2* | -1,308134876 |
| *Inf2* | -1,305523318 |
| *1810006K21Rik* | -1,303392993 |
| *Sema3d* | -1,301370801 |
| *Casc4* | -1,300737464 |
| *Sod1* | -1,298537699 |
| *Xkr7* | -1,298181936 |
| *Sdr39u1* | -1,297864843 |
| *Enam* | -1,296962245 |
| *Ndufa4* | -1,296891905 |
| *D730048J04Rik* | -1,294800162 |
| *9130008F23Rik* | -1,292325389 |
| *Pop7* | -1,292189086 |
| *Pkn3* | -1,291551319 |
| *Olfr945* | -1,291242471 |
| *Brox* | -1,287975374 |
| *Tmem186* | -1,287403822 |
| *Zbtb8os* | -1,287200299 |
| *Uba3* | -1,286957338 |
| *Mup21* | -1,286226343 |
| *Pfn2* | -1,285523005 |
| *Eps15l1* | -1,284282065 |
| *Specc1l* | -1,284199991 |
| *Slc25a39* | -1,283583505 |
| *Api5* | -1,282585906 |

| | |
|---|---|
| *Olfr700* | -1,282158168 |
| *Srpx* | -1,281852382 |
| *Vsig8* | -1,280462536 |
| *Lrriq3* | -1,279881694 |
| *Duox2* | -1,279631298 |
| *U2af1l4* | -1,279510622 |
| *E2f2* | -1,277415592 |
| *Pyhin1* | -1,276718936 |
| *Nostrin* | -1,276339209 |
| *Myom3* | -1,276252224 |
| *Cnn3* | -1,276137049 |
| *Ranbp6* | -1,275209179 |
| *Fbxw17* | -1,274967 |
| *Cops3* | -1,274851938 |
| *Efna5* | -1,274781346 |
| *Xpc* | -1,274532387 |
| *Tex261* | -1,273682371 |
| *Lgmn* | -1,272367797 |
| *Npy* | -1,270776985 |
| *Mgat4c* | -1,270751025 |
| *Mael* | -1,268969858 |
| *Ier3ip1* | -1,268384309 |
| *Mfsd2b* | -1,268038345 |
| *Igbp1* | -1,266852769 |
| *Evpl* | -1,265833241 |
| *Nlrp4f* | -1,263717797 |
| *2610109H07Rik* | -1,262002481 |
| *4930403N07Rik* | -1,259641122 |
| *Cenpv* | -1,258652727 |
| *Gtpbp6* | -1,258337832 |
| *Fancg* | -1,257401828 |
| *Phlda3* | -1,256265946 |
| *Gm3336* | -1,256261053 |
| *Parm1* | -1,256086104 |
| *Gbgt1* | -1,255753789 |
| *Gm8787* | -1,25483148 |
| *Dzip1l* | -1,253902765 |
| *Aqp7* | -1,252512866 |
| *Sfrp2* | -1,250753592 |
| *Arf5* | -1,25034454 |
| *Ect2* | -1,248760636 |
| *Olfr1209* | -1,246035689 |

| | |
|---|---|
| *Tuft1* | -1,245796679 |
| *Spef2* | -1,244788169 |
| *Olfr670* | -1,244517101 |
| *Krt12* | -1,243936889 |
| *Ktn1* | -1,2434541 |
| *Celf3* | -1,241094726 |
| *Mtmr6* | -1,240833712 |
| *Col5a3* | -1,240455052 |
| *Spinkl* | -1,239216128 |
| *Gpx7* | -1,23916919 |
| *Zfpm1* | -1,238660566 |
| *Pi4ka* | -1,237732368 |
| *Scn3b* | -1,23735587 |
| *Mrgprx2* | -1,236989673 |
| *Fcrl5* | -1,235867202 |
| *Myh9* | -1,235328656 |
| *Atp5f1* | -1,234963953 |
| *1110002B05Rik* | -1,231897976 |
| *Olfr221* | -1,230743835 |
| *Mpv17* | -1,229893134 |
| *Mtf2* | -1,229659114 |
| *Atad2b* | -1,228982432 |
| *Tfg* | -1,228626772 |
| *Sidt2* | -1,227894905 |
| *Capza1* | -1,227836027 |
| *Ren1* | -1,227533786 |
| *Maml3* | -1,226934087 |
| *Aldh3b1* | -1,226813792 |
| *Nup133* | -1,225403531 |
| *Mocs1* | -1,22525317 |
| *Dvl1* | -1,225249904 |
| *Cadps* | -1,225128131 |
| *Mamld1* | -1,225110646 |
| *Olfr860* | -1,224953934 |
| *Tcf4* | -1,224436969 |
| *Slc25a27* | -1,222644019 |
| *Zc3h7a* | -1,221816463 |
| *Olfr483* | -1,220893524 |
| *Polr1c* | -1,220749877 |
| *Klf11* | -1,220338491 |
| *Zbtb9* | -1,217940755 |
| *Gm3238* | -1,217531803 |

| | |
|---|---|
| *Tas2r134* | -1,217333863 |
| *Herc2* | -1,217044526 |
| *E2f6* | -1,216596423 |
| *Erich1* | -1,216595743 |
| *Prmt2* | -1,216180957 |
| *Olfr1061* | -1,215367805 |
| *Slc25a47* | -1,214130421 |
| *Atxn10* | -1,213877755 |
| *Nat10* | -1,213846933 |
| *Ccdc126* | -1,213445262 |
| *Rprd2* | -1,212988179 |
| *Prosapip1* | -1,212111207 |
| *Fgf11* | -1,212096844 |
| *Acsl1* | -1,211859181 |
| *Ube2o* | -1,210985845 |
| *Plekhb2* | -1,210958426 |
| *Cox5b* | -1,210508061 |
| *Olfr1202* | -1,209641542 |
| *Tufm* | -1,209638626 |
| *Herpud1* | -1,209560286 |
| *Acer2* | -1,209113028 |
| *Olfr1507* | -1,207928948 |
| *Fbxl20* | -1,207638676 |
| *Ccl27a* | -1,206946671 |
| *Pitpnc1* | -1,205830578 |
| *Mettl7a1* | -1,205410078 |
| *Kynu* | -1,20368559 |
| *Als2cr12* | -1,203109415 |
| *Ttn* | -1,201029147 |
| *Dnajc12* | -1,200235019 |
| *Gtpbp10* | -1,20005688 |
| *Nlgn2* | -1,19852276 |
| *Gbp6* | -1,198286202 |
| *Oas1h* | -1,19648979 |
| *Cpd* | -1,196336025 |
| *Chmp3* | -1,195427459 |
| *Hivep2* | -1,195351278 |
| *Ccdc103* | -1,194716424 |
| *Gm11487* | -1,193601032 |
| *Snca* | -1,192052685 |
| *Ticam2* | -1,191483112 |
| *Pex11b* | -1,191014161 |

| | |
|---|---|
| *Dtna* | -1,190883298 |
| *C030006K11Rik* | -1,190679859 |
| *Tas2r139* | -1,190672924 |
| *Pspc1* | -1,190217179 |
| *Tead2* | -1,190103898 |
| *Smg7* | -1,188825693 |
| *Inpp4a* | -1,187289194 |
| *Trps1* | -1,186174387 |
| *2310033P09Rik* | -1,186135953 |
| *Aph1c* | -1,185803196 |
| *Hist1h2bk* | -1,185798563 |
| *Uso1* | -1,185472431 |
| *Dtx1* | -1,185447259 |
| *Olfr10* | -1,185339441 |
| *Il21r* | -1,185222581 |
| *Rarb* | -1,184951647 |
| *Mst1* | -1,184871793 |
| *Alyref2* | -1,183863329 |
| *Polr3f* | -1,18208534 |
| *Pramef8* | -1,182077459 |
| *Phldb1* | -1,182000842 |
| *Zglp1* | -1,180705844 |
| *Mgst1* | -1,180053373 |
| *Dpagt1* | -1,178039796 |
| *Tmsb15a* | -1,176520555 |
| *Mmab* | -1,175512407 |
| *Bbox1* | -1,175190246 |
| *Efnb1* | -1,174955406 |
| *Slc1a6* | -1,173359913 |
| *Ep300* | -1,172768843 |
| *Evi2a* | -1,171034326 |
| *Ccdc67* | -1,170784317 |
| *Hmgxb4* | -1,170757407 |
| *Nsf* | -1,168953129 |
| *Map6d1* | -1,168324032 |
| *Arsa* | -1,167828651 |
| *Trmt2b* | -1,166546329 |
| *Fgf21* | -1,165474104 |
| *Cpne5* | -1,165316863 |
| *Gm10345* | -1,16432319 |
| *Cage1* | -1,163050094 |
| *Olfr541* | -1,162582516 |

| | |
|---|---|
| *Dysf* | -1,162357251 |
| *Olfr1494* | -1,162214241 |
| *Pias2* | -1,161241888 |
| *Gfm1* | -1,160999126 |
| *2310021P13Rik* | -1,160625527 |
| *Hsf5* | -1,160274524 |
| *Mccc2* | -1,159596323 |
| *Gsdmc2* | -1,159191965 |
| *Rassf7* | -1,159188003 |
| *Gmfb* | -1,158051486 |
| *Fkbp14* | -1,157859909 |
| *Grrp1* | -1,157034177 |
| *Adcyap1* | -1,156985473 |
| *Ccl6* | -1,156081283 |
| *Pot1a* | -1,154735218 |
| *Ube2h* | -1,154020628 |
| *Raly* | -1,153929753 |
| *Peg10* | -1,153834266 |
| *Fbxo41* | -1,153718693 |
| *Zfp119b* | -1,153597164 |
| *Olfr694* | -1,153341662 |
| *Cyp4f18* | -1,153133056 |
| *Cks2* | -1,150052216 |
| *Tmem41a* | -1,147827616 |
| *Ints3* | -1,147417568 |
| *Vps41* | -1,146853673 |
| *Ppap2a* | -1,146804283 |
| *Prss23* | -1,145450881 |
| *Zfand2a* | -1,145328051 |
| *Tti1* | -1,14375122 |
| *Prss45* | -1,143055992 |
| *Kdm3b* | -1,142744878 |
| *Hspa12b* | -1,141656425 |
| *Olfr285* | -1,140444142 |
| *Paqr4* | -1,140049873 |
| *Amac1* | -1,139879365 |
| *Zar1l* | -1,138906606 |
| *Txk* | -1,137552103 |
| *Ddx41* | -1,137378069 |
| *Zfp187* | -1,137138179 |
| *Pdzd3* | -1,136555357 |
| *Sec24a* | -1,136554921 |

| | |
|---|---|
| *Psma8* | -1,136246744 |
| *6430527G18Rik* | -1,134535006 |
| *Slc12a4* | -1,132558746 |
| *Dhx16* | -1,13221206 |
| *Pnrc2* | -1,132151702 |
| *Gm5150* | -1,13212985 |
| *Igfn1* | -1,132083374 |
| *Erf* | -1,13170217 |
| *Gdpd1* | -1,131457429 |
| *Slc26a5* | -1,129174898 |
| *Arid5b* | -1,128170153 |
| *Eif5a* | -1,126190041 |
| *Kcna1* | -1,125479757 |
| *Dhrs7* | -1,124968124 |
| *Npy5r* | -1,123925165 |
| *Tpbg* | -1,122842778 |
| *Kcnip4* | -1,122397286 |
| *Tnfaip3* | -1,122087609 |
| *Rab11b* | -1,121622636 |
| *1700109H08Rik* | -1,12154542 |
| *Rprd1b* | -1,12100935 |
| *Aldh3a2* | -1,120917963 |
| *Map4k1* | -1,120332978 |
| *Myh3* | -1,120075222 |
| *Wdr77* | -1,119320754 |
| *Mrgpra1* | -1,119193428 |
| *Parp11* | -1,119161521 |
| *Adrb2* | -1,118806713 |
| *Cntnap4* | -1,118446601 |
| *Ikzf2* | -1,118148484 |
| *Timm13* | -1,117576609 |
| *Tmem131* | -1,117405486 |
| *Msto1* | -1,113712506 |
| *Nlrp5* | -1,113266127 |
| *2400001E08Rik* | -1,113125954 |
| *Ttc8* | -1,113006147 |
| *Mto1* | -1,112945189 |
| *Ppp3cc* | -1,112247123 |
| *Dock10* | -1,111302283 |
| *Rasa4* | -1,110311829 |
| *Klk14* | -1,109533623 |
| *Olfr228* | -1,108996011 |

| | |
|---|---|
| *Olfr747* | -1,107393033 |
| *4921524L21Rik* | -1,107299701 |
| *Fam126a* | -1,106411345 |
| *Cmtm3* | -1,105782587 |
| *Nxph4* | -1,105767734 |
| *Cnnm3* | -1,103860285 |
| *Reg1* | -1,103521703 |
| *Grm3* | -1,10348329 |
| *Pdap1* | -1,103299616 |
| *Gmcl1* | -1,102436029 |
| *Gm806* | -1,102241923 |
| *Olfr1382* | -1,101009249 |
| *Hira* | -1,100251202 |
| *Actr3* | -1,10005513 |
| *Cep290* | -1,099699649 |
| *Gm97* | -1,099686246 |
| *Pdrg1* | -1,099539277 |
| *Vbp1* | -1,099033985 |
| *Tssk2* | -1,098879177 |
| *Foxk2* | -1,098369386 |
| *Vgll1* | -1,097957291 |
| *2010109A12Rik* | -1,097820443 |
| *Dab2ip* | -1,097214079 |
| *Bphl* | -1,096846506 |
| *Iqcd* | -1,096454475 |
| *Gm5415* | -1,095072331 |
| *BC027072* | -1,094845481 |
| *Tbx22* | -1,094634622 |
| *Matn3* | -1,094558984 |
| *Rragc* | -1,094189202 |
| *Nphp4* | -1,09356743 |
| *Acyp2* | -1,092898221 |
| *Cyp4v3* | -1,09241078 |
| *Paox* | -1,091378255 |
| *Nrxn3* | -1,091291349 |
| *Kcnj8* | -1,091185898 |
| *Rasl10a* | -1,090807992 |
| *Hspa12a* | -1,090291808 |
| *Itgb2* | -1,089534406 |
| *Siglec5* | -1,089252529 |
| *Olfr661* | -1,088678458 |
| *Myef2* | -1,088535727 |

| | |
|---|---|
| *Rbm47* | -1,087587468 |
| *Skint3* | -1,085777024 |
| *Olfr1383* | -1,085286526 |
| *Prl7c1* | -1,085057212 |
| *Vmn1r5* | -1,084733299 |
| *Slc10a2* | -1,084374026 |
| *Rassf4* | -1,083970051 |
| *Mmp3* | -1,083947408 |
| *Hook1* | -1,083919033 |
| *Creb3l1* | -1,083279727 |
| *Olfr450* | -1,083267234 |
| *Mtfp1* | -1,083150297 |
| *Olfr342* | -1,082059281 |
| *Fam163a* | -1,081799623 |
| *Fermt2* | -1,081767194 |
| *Slc41a1* | -1,08079222 |
| *1700125H20Rik* | -1,080515582 |
| *Psmd5* | -1,079874587 |
| *2610028H24Rik* | -1,079677101 |
| *Ung* | -1,079319029 |
| *4930525M21Rik* | -1,079018853 |
| *Olfr1246* | -1,078860374 |
| *Cct8l1* | -1,078003424 |
| *Mat2a* | -1,077577994 |
| *Qk* | -1,076992361 |
| *Hdhd2* | -1,076853369 |
| *Vhl* | -1,076751126 |
| *4933434E20Rik* | -1,076520408 |
| *2610528E23Rik* | -1,076210979 |
| *Cdkn2d* | -1,076051418 |
| *Itgb1* | -1,075785084 |
| *Tfdp1* | -1,075604104 |
| *Olfr1049* | -1,075538369 |
| *Tmigd1* | -1,075388535 |
| *Clcn2* | -1,075256729 |
| *Sgcd* | -1,074924296 |
| *Ros1* | -1,074726346 |
| *Ildr2* | -1,074174311 |
| *Eefsec* | -1,073750424 |
| *4930578I06Rik* | -1,073363423 |
| *Il2rg* | -1,073187477 |
| *Rcbtb2* | -1,072776936 |

| | |
|---|---|
| *Olfr429* | -1,072707629 |
| *Cln6* | -1,072061531 |
| *Numbl* | -1,071728242 |
| *Vasn* | -1,071345493 |
| *Trim9* | -1,070643018 |
| *Sptlc1* | -1,070062481 |
| *Vars2* | -1,06964196 |
| *Olfr920* | -1,069598139 |
| *Slco3a1* | -1,069155063 |
| *Slc30a4* | -1,068764158 |
| *Slc22a4* | -1,068315498 |
| *Plscr4* | -1,067785977 |
| *Cacnb3* | -1,067482932 |
| *Bbip1* | -1,065618324 |
| *Rnh1* | -1,065598897 |
| *Tas2r113* | -1,065305756 |
| *Vmn1r236* | -1,0652086 |
| *Trpv4* | -1,064937666 |
| *Zhx1* | -1,064899784 |
| *Rhoq* | -1,064033849 |
| *Spty2d1* | -1,063705214 |
| *BC057079* | -1,063302035 |
| *Vnn1* | -1,063282911 |
| *Olfr585* | -1,063209565 |
| *Mc3r* | -1,062954396 |
| *4933416C03Rik* | -1,061720669 |
| *Incenp* | -1,061594508 |
| *Olfr1226* | -1,061256589 |
| *Ahcyl2* | -1,060890147 |
| *Acot1* | -1,060126335 |
| *Fhl5* | -1,060060396 |
| *Tdpoz2* | -1,058550218 |
| *1500032L24Rik* | -1,058361951 |
| *Tyrobp* | -1,057860148 |
| *Tacr1* | -1,057697932 |
| *LOC100861668* | -1,057269808 |
| *Gzmn* | -1,057162406 |
| *Chka* | -1,056536395 |
| *Dpy19l2* | -1,05630736 |
| *Gm14378* | -1,056031519 |
| *Mov10l1* | -1,055449484 |
| *F13a1* | -1,054926112 |

| | |
|---|---|
| *Hps5* | -1,054678147 |
| *Cdcp2* | -1,053680932 |
| *Mcph1* | -1,052689235 |
| *Rnasel* | -1,05206687 |
| *Qars* | -1,051817184 |
| *Os9* | -1,051775962 |
| *Olfr1254* | -1,051608758 |
| *Vgll3* | -1,051367443 |
| *Cryba2* | -1,050566336 |
| *Cyp2c37* | -1,049803914 |
| *Olfr1243* | -1,049563283 |
| *Rara* | -1,048627076 |
| *Gpkow* | -1,048508976 |
| *Fam196a* | -1,047376609 |
| *Olfr921* | -1,045679719 |
| *Gpr37l1* | -1,045362862 |
| *Vmn1r77* | -1,045085533 |
| *Naa11* | -1,044522871 |
| *2410003K15Rik* | -1,043234853 |
| *Ppif* | -1,042980759 |
| *Srsf6* | -1,042803485 |
| *BC029214* | -1,04249253 |
| *Gm14124* | -1,041115013 |
| *Syn1* | -1,040951317 |
| *Olfr609* | -1,040493971 |
| *Gpha2* | -1,03854738 |
| *Pqlc3* | -1,037227929 |
| *Rnf125* | -1,036750668 |
| *Nxf3* | -1,036670464 |
| *Tmem53* | -1,036615792 |
| *Slc37a1* | -1,036419815 |
| *Grtp1* | -1,035783257 |
| *Ephb2* | -1,035736553 |
| *Alkbh8* | -1,035362441 |
| *4430402I18Rik* | -1,03278074 |
| *Mrpl30* | -1,032412231 |
| *Hexa* | -1,03201864 |
| *Isg15* | -1,031763978 |
| *Olfr1402* | -1,031676433 |
| *Nfx1* | -1,031465315 |
| *Prom2* | -1,030837347 |
| *Sart3* | -1,030482053 |

| | |
|---|---|
| *Scaf4* | -1,030321652 |
| *Nup93* | -1,028361845 |
| *Ppp1r1b* | -1,027816146 |
| *Olfr345* | -1,027419491 |
| *Tcn2* | -1,027217452 |
| *Ptdss2* | -1,026474205 |
| *Ccdc150* | -1,026292594 |
| *Klk12* | -1,026290446 |
| *Apol10b* | -1,025660772 |
| *Wnt8a* | -1,023516154 |
| *Itgb2l* | -1,022269016 |
| *Gpr65* | -1,022238563 |
| *Tmem30c* | -1,022063821 |
| *Arhgap30* | -1,02191134 |
| *H2bfm* | -1,021608547 |
| *Cacnb4* | -1,021023869 |
| *Tjp1* | -1,020712013 |
| *Mybph* | -1,020642946 |
| *Mob4* | -1,02062948 |
| *Olfr746* | -1,020296053 |
| *P2rx1* | -1,020017647 |
| *Edf1* | -1,01984055 |
| *Psma2* | -1,019555374 |
| *Fam72a* | -1,018872619 |
| *Lyrm2* | -1,018691653 |
| *1700020A23Rik* | -1,018455549 |
| *Fbxo33* | -1,018319285 |
| *Obfc1* | -1,01820196 |
| *Fam78b* | -1,018015827 |
| *Mettl8* | -1,017680814 |
| *Sep-04* | -1,01758159 |
| *Saal1* | -1,017370222 |
| *Prl3d2* | -1,016877179 |
| *Zfp68* | -1,016418497 |
| *Hmgn5* | -1,01625975 |
| *Tmem184c* | -1,015718716 |
| *Edar* | -1,015495335 |
| *Hnrnpm* | -1,014786246 |
| *Ufd1l* | -1,01474349 |
| *Nup188* | -1,013406292 |
| *Fam189b* | -1,013180873 |
| *9930012K11Rik* | -1,012458577 |

| | |
|---|---|
| *D630039A03Rik* | -1,012373815 |
| *Hif1a* | -1,011739771 |
| *Gpc2* | -1,01096625 |
| *Avil* | -1,010891387 |
| *Zscan12* | -1,010297237 |
| *Olfr1318* | -1,009808297 |
| *Pate2* | -1,008612355 |
| *Gabrp* | -1,008540961 |
| *Usp4* | -1,008485512 |
| *Antxr1* | -1,007622605 |
| *Sep-05* | -1,007517748 |
| *Agrp* | -1,007115908 |
| *Pard6g* | -1,006424976 |
| *Nktr* | -1,0064023 |
| *Mtap* | -1,006261503 |
| *Jtb* | -1,005444952 |
| *Asb10* | -1,005015513 |
| *Tex13a* | -1,00488603 |
| *Kcna2* | -1,0044124 |
| *Vmn2r53* | -1,003235138 |
| *Gtf2h1* | -1,002962691 |
| *Olfr1140* | -1,00224651 |
| *H2afb1* | -1,002022146 |
| *Pgm5* | -1,00112037 |
| *Fgf4* | -1,001084138 |
| *Prima1* | -1,000630362 |
| *Ms4a4c* | -1,000177673 |

**Table S3**. **Survival genes identified by whole genome CRISPR/Cas9 screening in INK128-treated mESCs and vehicle mESCs.**

Genes were identified by calculating the fold change between the sgRNAs abundance in doxycycline-treated cells (INK128-treated or vehicle) versus their untreated counterparts.
